# MESSAR: Automated recommendation of metabolite substructures from tandem mass spectra

**DOI:** 10.1101/134189

**Authors:** Youzhong Liu, Aida Mrzic, Pieter Meysman, Thomas De Vijlder, Edwin P. Romijn, Dirk Valkenborg, Wout Bittremieux, Kris Laukens

**Author notes:** Authors contributed equally and should be considered co-first authors (Youzhong Liu, Aida Mrzic).

## Abstract

Despite the increasing importance of non-targeted metabolomics to answer various life science questions, extracting biochemically relevant information from metabolomics spectral data is still an incompletely solved problem. Most computational tools to identify tandem mass spectra focus on a limited set of molecules of interest. However, such tools are typically constrained by the availability of reference spectra or molecular databases, limiting their applicability to identify unknown metabolites. In contrast, recent advances in the field illustrate the possibility to expose the underlying biochemistry without relying on metabolite identification, in particular via substructure prediction. We describe an automated method for substructure recommendation motivated by association rule mining. Our framework captures potential relationships between spectral features and substructures learned from public spectral libraries. These associations are used to recommend substructures for any unknown mass spectrum. Our method does not require any predefined metabolite candidates, and therefore it can be used for the partial identification of unknown unknowns. The method is called MESSAR (MEtabolite SubStructure Auto-Recommender) and is implemented in a free online web service available at messar.biodatamining.be.

**Author Summary:** Mass spectrometry is one of most used techniques to detect and identify metabolites. However, learning metabolite structures directly from mass spectrometry data has always been a challenging task. Thousands of mass spectra from various biological systems still remain unanalyzed simply because no current bioinformatic tools are able to generate structural hypotheses. By manually studying mass spectra of standard compounds, chemists discovered that metabolites that share common substructures can also share spectral features. As data scientists, we believe that such relationships can be unraveled from massive structure and spectra data by machine learning. In this study, we adapted “association rule mining”, traditionally used in market basket analysis, to structural and spectral data, allowing us to investigate all spectral features - metabolite substructures relationships. We further collected all statistically sound relationships into a database and used them to assign substructral hypotheses to unexplored spectra. We named our approach MESSAR, MEtabolite SubStructure Auto-Recommender, available to the metabolomics and mass spectrometry community as a free and open web service.

## Introduction

Metabolomics is an emerging “omics” science involving the high-throughput analysis of metabolites or small biomolecules, with highly relevant applications in drug and biomarker discovery [1, 2]. One standard method for metabolite analysis is mass spectrometry (MS), preceded by a separation technique, such as gas chromatography (GC) or liquid chromatography (LC). Advances in MS instrumentation enable the simultaneous detection and quantification of thousands of metabolites in a biological sample. Chemical identification of these metabolites is a key step towards biochemical interpretation of studied samples. To obtain structural information, tandem MS (MS/MS) is applied to record the fragment *m*/*z* of targeted molecules. Structure elucidation from MS/MS data has always been a challenging and time-consuming task with a vast number of potentially interesting metabolites that are still unknowns. The main reason is that current MS/MS databases (spectral libraries) only contain a limited number of historical spectra, far below the number of metabolites in reality [3, 4].

Advances in computational tools have led to a considerable extension of the search space that can be examined and have resulted in an improvement of the identification accuracy by using massive molecular databases (for example, PubChem currently contains over 100 million compounds [5]). These tools start by filtering the molecular database using the precursor *m*/*z* of the unknown spectra, yielding up to thousands of structure candidates.

To subsequently score and rank these candidates two categories of algorithms have been proposed. First, *in silico* fragmentation tools simulate theoretical spectra for each candidate metabolite and compare those with the query spectrum [6–9]. Second, machine learning (ML) methods learn intermediate representations, such as molecular fingerprints [10] from historical spectrum–structure relationships. These representations are then used to score spectrum–candidate matches. A typical example hereof is the CSI:FingerID tool [11]. However, both types of algorithms still have certain limitations. For example, *in silico* fragmentation tools only cover simple fragmentations and cannot accurately simulate complex rearrangement reactions [12]. Additionally, none of these tools can identify “unknown unknowns”, i.e. compounds that have not been structurally described yet and are therefore not present in any molecular database. As a result typically only a fraction of compounds can be identified correctly [13].

Recently a third category of computational tools has been introduced in non-targeted metabolomics. With a focus on the *de novo* identification of unknowns, these tools aim to predict substructures rather than the full metabolite structure. The basic concept for this strategy is that metabolites often share substructures, resulting in similar patterns in their MS/MS spectra. Typical spectral features are product ions, neutral losses, or mass differences [14–17].

One important tool to explore spectral similarity is the Global Natural Products Social Molecular Networking (GNPS) resource [18]. GNPS consists of a large metabolite network where metabolites with similar MS/MS spectra are connected so that structurally annotated metabolites can be used for the identification of their neighbors. However, such a network-based approach may fail to connect metabolite pairs if they have a low spectral similarities despite sharing important substructures such as small functional groups. Additionally, the annotation of neighboring nodes still requires manual intervention.

MS2LDA is a recent framework proposed by van der Hooft et al. [20–22]. It decomposes unlabeled MS/MS spectra into patterns of co-occurring fragments and losses, referred to as “Mass2Motifs”, which are indicative of biological substructures. These Mass2Motifs patterns are automatically extracted from complex MS/MS spectra using unsupervised text mining techniques. However, the extracted motifs have to be structurally annotated based on expert knowledge, which requires extensive domain expertise and is time-consuming.

Here we introduce a new method for the recommendation of substructures for MS/MS spectra, working independently from molecular databases. Our tool, called MESSAR (MEtabolite SubStructure Auto-Recommender), is inspired by the concept of association rule mining (ARM). ARM has been designed to discover interesting relations based on frequently co-occurring items, and it has previously been used to find relations between unassigned mass spectra [23–25].

We use a collection of labeled spectra (reference spectra with a known corresponding molecular structure from a spectral library) to mine co-occurring patterns of the form “MS/MS features + substructures”, yielding “MS/MS feature(s) → substructure” rules. These rules capture recurring patterns found in mass spectra and assign them potential substructures. As such, they can be used to partially replace expert-driven annotations in tools such as MS2LDA.

The current MESSAR model is a database of 8378 “MS/MS feature(s) → substructure” rules derived from the GNPS spectral library. All rules were statistically evaluated on training and independent testing spectra, and compared with rules generated from a decoy GNPS spectral library. When annotating a new spectrum, MESSAR identifies all of its spectral features that match the rule database, after which any rules suggesting similar substructures are aggregated and maximal common substructures (MCS) are reported.

MESSAR is currently designed for positive ion mode LC-MS/MS data. It is available as a free online web service at messar.biodatamining.be.

## Materials and Methods

### Training spectral libraries

MESSAR generates rules from target and decoy GNPS spectral libraries built by Scheubert et al. (Fig 1A). According to the data descriptions [26], the target library consists of 4138 positive ion high-quality labeled spectra acquired on Q-TOF instruments. For each spectrum, Scheubert et al. have computed a fragmentation tree that annotates a subset of fragments with molecular formulas and removed non-annotated peaks that usually represent isotopic peaks, chemical noise, … [11]. Meanwhile, neutral exact masses were assigned to formula annotated fragments. The decoy library was a randomized version of the target library. The decoy process kept the library labels (molecular structures) and randomized the corresponding mass spectra through re-rooted fragmentation trees (Fig 1A). Such process mimics very “noisy” experimental spectra, therefore extracted patterns can probably reflect spurious feature-substructure relations in the target library. For both target and decoy libraries, mass spectra representing the same structure were combined, and duplicated fragments were removed. The final training data consists of 3146 target and decoy spectra (S1 Data).

**Fig 1.**
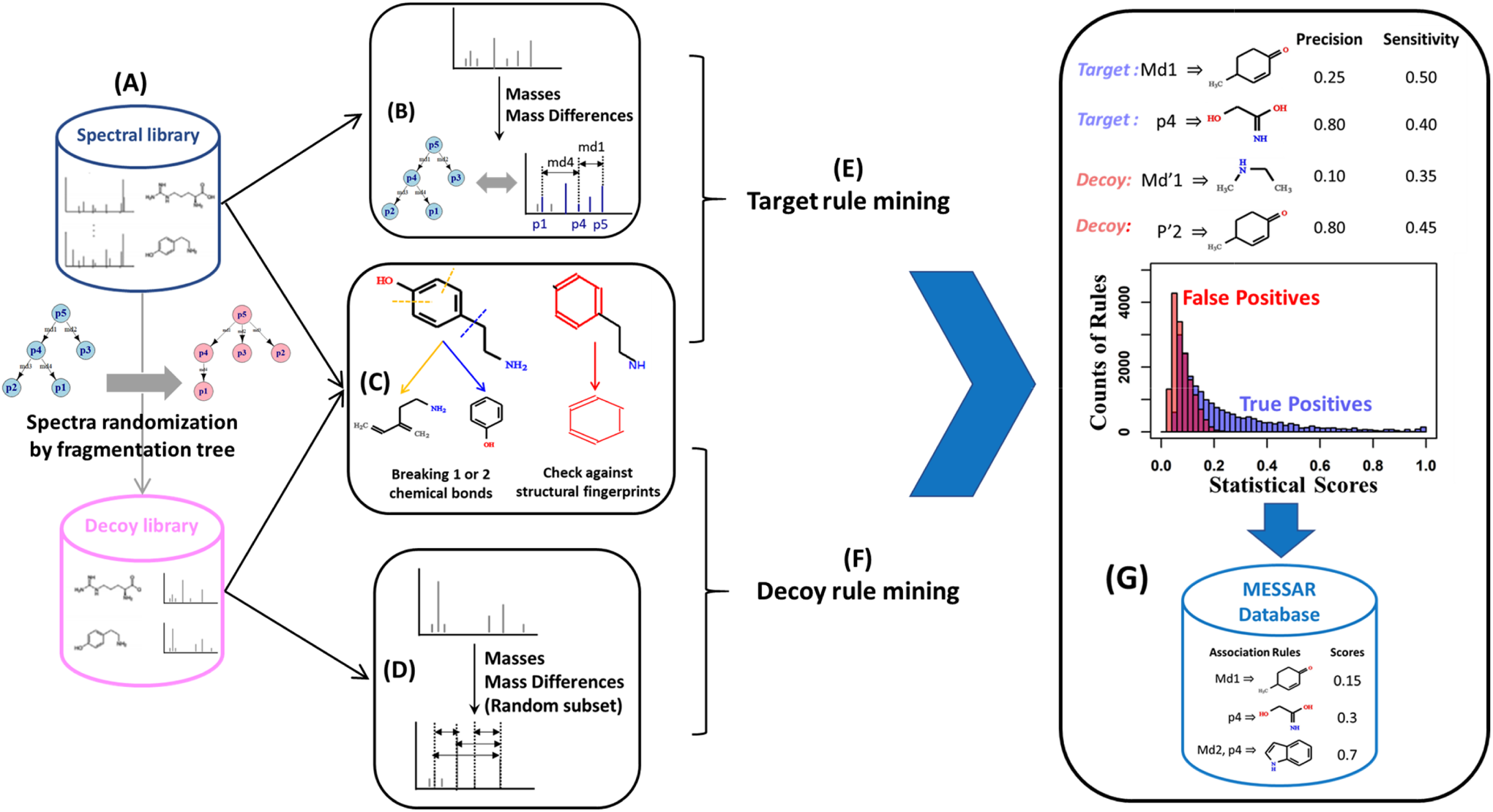
Workflow for target and decoy rule generation. Based on training data from an LC-MS/MS spectral library substructure recommendations are generated. Our method consists of (A) retrieving training MS/MS spectra, (B) extracting spectral features (neutral exact masses and mass differences) from the target library, (C) generating molecular substructures, (D) extracting spectral features from the decoy library, (E) rule mining using substructures and target spectral features, (F) rule mining using substructures and decoy spectral features, (G) statistical evaluation of target and decoy rules, filtering target rules and saving valid rules to the final database.

### Spectral feature extraction

Both fragment masses and distances between fragments (mass differences) were interesting spectral features for model training. However, as an example, if a training spectrum has 20 mass peaks, extracted spectral features consist of 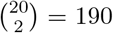 mass differences. If we use all mass differences, rule generation can be computationally expensive and highly dominated by mass differences with no structural information. Therefore, we only used edges of fragmentation trees as target spectral features since they represent losses due to fragmentation reactions and probably hold structural information (Fig 1B). For the decoy spectrum of the same molecule, we extracted a random subset of 20% mass differences (Fig 1D). In this way, a similar amount of spectral features was used to generate target and decoy rules.

### Substructure generation

For every metabolite in the spectral library a set of substructures is created by combining two approaches: i) by checking the presence of predefined substructures, ii) through “breaking of retrosynthetically interesting chemical substructures” (BRICS) algorithm [27] (RDKit https://www.rdkit.org/ in Python). For i), we took 1483 CSI:FingerID molecular fingerprints (all types except for ECFP4) and converted them into substructures [11]. The SMILES code of corresponding substructures are available in S1 Data. For the second approach, we disconnected one or two chemical bonds in the metabolite and collected the resulting two or three substructures obtained from every iteration (Fig 1C). CHON substructures that contain less than five carbon and oxygen atoms were considered trivial and therefore discarded. All substructures were represented by SMILES codes in the rule database.

### Rule generation, statistical analysis and filtering

After combining substructures with extracted spectral features, association rules were mined separately for target and decoy libraries (Fig 1EF). Details about ARM can be found in S1 Text.

A MESSAR rule with shape *X* ⇒ *Y* describes the potential dependency of a substructure (*Y*) on a spectral feature pattern/feature set (*X* can contain up to three co-occurring masses and mass differences). Therefore, each rule can be considered as a binary classifier that decides whether a metabolite contains a substructure according to the corresponding spectral feature presence/absence. The predictive power of the rules was evaluated based on a confusion matrix, as follows:

We calculated two statistical metrics for each target and decoy rule, namely *precision* and *sensitivity* (Fig 1G). A perfect*precision* score of 1.0 means that the metabolite always contains substructure *Y* if we detect the spectra feature set *X* whereas a *sensitivity* of 1.0 means that all metabolites containing substructure *Y* must have feature set *X* in their spectra:

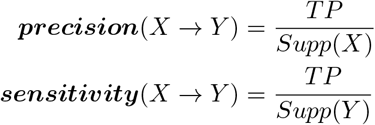

Target rules were filtered by defining a *sensitivity* threshold for 1% FDR (Fig 1G). Based on the target-decoy method, the simple FDR associated with a particular *sensitivity* threshold is the ratio between the number of accepted decoy rules (above the threshold) and target rules [31]. The simple FDR was estimated using the *prozor* package in R (https://github.com/protviz/prozor).

### Test data

The predictive power of individual target rules was first evaluated on an independent “MASSBANK” data set (S2 Data). This test set contained 5164 labeled TOF-MS/MS spectra derived from the MASSBANK spectral library (http://mona.fiehnlab.ucdavis.edu/). Compounds behind these spectra are metabolites, drugs and natural products.

The second test data set “MASSBANK CASMI” (S2 Data) was used to assess substructure recommendation for unknown spectra. This test set consists of 185 labeled TOF-MS/MS data, including 34 drugs and 126 metabolites from MASSBANK, as well as 25 spectra from the open contest CASMI 2017 (http://casmi-contest.org/2017).

When creating both test sets, we discarded all compounds that overlapped with the MESSAR training set based on the first block of InChiKey. “MASSBANK CASMI” does not contain any compounds used for CSI:FingerID model training (https://bio.informatik.uni-jena.de/software/sirius/). Mass spectra representing the same structure in each test set were combined, and duplicate fragments were removed.

### Maximum common substructure

When using MESSAR rules to annotate unknown spectra, we computed the maximum common substructures (MCSs) of a matched rule set in order to extract meaningful core substructures and to reduce the uncertainty of prediction. We first screened all mtached rule pairs to identify “analogous rules” in which the head of rules had a Tanimoto similarity score higher than 0.5 [28]. We extracted from all analogous rule pairs maximum common substructures using the *rcdk* package in R. For motif annotation, the most frequent MCS of a rule set was reported. For unknown spectra annotation, each extracted MCS was scored by summing the *sensitivity* of all relevant rules.

### MESSAR webtool

The MESSAR web tool recommends substructures for an input spectrum based on the pre-trained rule database (Fig 1). MESSAR expects as input a list of MS/MS peaks and the precursor *m*/*z*. MESSAR then uses the fragments and the computed mass differences to query its database. The matched rules are ranked by their *sensitivity*. Users could use these rules to generate structural hypotheses. In addition, rules that suggest similar (or identical) substructure can be aggregated, leading to a ranked list of substructure recommendations. Three rule aggregation algorithms are available in the web tool: i)“Exhaustive”: maximum common substructures computed from all rules; ii) “Fast”: MCS calculation on 20 most sensitive rules; iii) “Naive”: simply combining rules that suggest identical substructures. For all three algorithms, the score of each substructure is the sum of *sensitivity* of all responsible rules. We recommend users to try out all three algorithms to discover the most reliable and meaningful substructures, but we only present results based on the “Exhaustive” algorithm in this paper. The client interface of the web tool was developed using the R Shiny framework.

### Substructure recommendation by CSI:FingerID and MS2LDA

CSI:FingerID (Windows GUI SIRIUS-4.0.1) was downloaded from https://bio.informatiuni-jena.de/software/sirius/. The “MASSBANK CASMI” data was submitted to the GUI as an mgf format file. The precursor formula was not available for fragmentation tree building, and all elements were considered (classical CHNOPS along with Br, Cl, F and I since they might be present in drug compounds). The annotation tolerance was set to 20 ppm. CSI:FingerID predicted five types of molecular fingerprints that can be translated into substructures (1483 substructures in total), namely CDK, PubChem CACTVS, Klekota-Roth, FP3 and MACCS. Substructures were ranked by probability. If CSI:FingerID was unsure about the elemental formula and assigned the same substructure multiple probabilities, the average probability was used. We only considered recommendations with a probability above 0.5.

The same mgf file was submitted to http://ms2lda.org/ for M2M searching and annotation. The minimum intensity of MS2 peaks was set at 1, and the width of MS2 bins at 0.005 Da. The motifs found were further inferred using predefined 500 GNPS motifs. The results are available online at http://ms2lda.org/basicviz/summary/951/.

## Results and Discussion

First, we briefly describe the model training (rule generation) procedure and illustrate the statistical significance of generated rules on training and test spectra. Second, the biochemical relevance of these rules is revealed through a comparison with MS2LDA patterns. Third, MESSAR was validated on two independent sets of test spectra for the prediction power of individual rules and all rules together. The performance of MESSAR for unknown spectra annotation was compared with CSI:FingerID. Finally, the usefulness of the MESSAR output was evaluated alongside CSI:FingerID and MS2LDA through 185 test spectra.

### Outline of MESSAR rule generation

MESSAR predicts molecular substructures from an MS/MS spectrum based on significant “MS/MS feature(s) → substructure”-like patterns (rules) derived from existing labeled spectra. To find such patterns, the classical ARM algorithm was applied on a subset of the GNPS spectral library. The algorithm discovers rules among labeled spectra based on user-defined parameters (S1 Text). An overview of the rule generation procedure from the spectral library is depicted in Fig 1. This includes the following steps described in the Materials and Methods section: (A) retrieving training data: target and decoy GNPS spectral libraries used by passatutto software [26], (B) extracting spectral features from the target library, (C) generating molecular substructures of training compounds, (D) extracting spectral features from the decoy library, (E) ARM on target features and substructures, (F) generating random rules from the decoy database, (G) statistical evaluation of target and decoy rules, filtering target rules by FDR estimation with the target-decoy method. All target rules above the score threshold chosen were saved in the MESSAR rule database.

### Statistical properties of MESSAR rules

Using the procedures described in S1 Text, we have generated 20747 and 15480 rules from target and decoy spectral libraries, respectively (before FDR-based filtering in step G). In the target database we can observe several expected substructure recommendations. For example, mass features in rules 1–3 (Table 2) reflect the molecular weight of recommended substructure. We found about 3% such rules that captured the direct link between masses/mass differences and the presence/loss of substructures (S1 File). The remaining 97% rules describe potential latent associations between spectral features and substructures (Table 2, Rules 4–9, S1 File) and cannot be explained directly.

In total, the 20747 target rules covered 215 sets of spectral features and 732 substructures. The same set of spectral features can recommend several different substructures (Rules 4-6), and reciprocally, the same substructure can be associated with multiple feature patterns (Rules 7-9). In the first scenario, the recommended substructures are usually very much alike, which can be explained by the presence of the same spectral feature(s) in similar training molecules with a minor substructure difference (S2 Fig). The second scenario is consistent with the concept of Mass2Motifs [20], that is, a complex substructure is indeed associated to co-occurring molecular fragments and losses (spectral patterns).

However, not all target MESSAR rules are meaningful because ARM can find spectral features-substructure associations that are present by chance. Since a direct biochemical evaluation (e.g. via molecular weight) is not feasible for most rules, we verified whether the statistical measures, *precision* and *sensitivity*, can be used to select meaningful rules. The most meaningful rule can be selected based on the highest *precision* or *sensitivity* (Table 2). Using the entire training set, we created confusion matrices for each rule and evaluated the two statistical measures. The decoy rules served as negative controls and were evaluated in the same way as the target rules (Fig 1G)

First, there was no correlation between the two metrics (Fig 2A). Points representing target/decoy rules can be separated based on sensitivity. We further compared the distributions of *precision* and *sensitivity* (Fig 2BC). It can be seen that the *precision* of most target and decoy rules lies between 0 and 0.4, and target/decoy distributions overlap over the entire *precision* range. In contrast, the *sensitivity* of the decoy rules centers around 0.07 and ranges mostly between 0 and 0.2, while target rules display a skewed Poisson-like distribution around 0.16 with a long right tail up to 1 (Fig 2C). Moreover, there was little overlap between target/decoy distributions.

**Fig 2.**
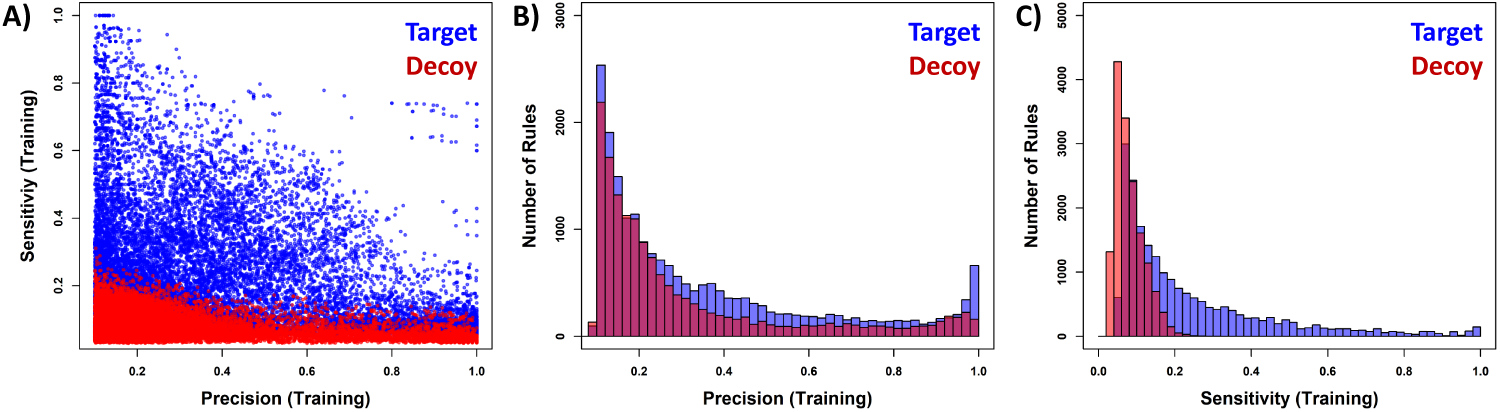
Statistical evaluation of MESSAR rules. (A) scatter plot between precision and sensitivity, (B) precision and (C) sensitivity distribution of target and decoy rules.

In summary, *sensitivity* appears to be more suitable to distinguish meaningful rules from spurious patterns. It will be the only metrics used for FDR control and model validation (substructure recommendation) in the manuscript. After filtering rules for statistical soundness and for 1% FDR (threshold defined based on the shape of the curve in S3 Fig), target rules with a *sensitivity* higher than 0.20 were kept. The final MESSAR database consists of 8378 association rules (S1 File).

### Comparison between MESSAR rules and MS2LDA patterns

MS2LDA [20, 22] is an unsupervised tool that discovers patterns across fragmentation spectra. As it operates within a similar scope as MESSAR, a detailed comparison is warranted. The major difference between MESSAR and MS2LDA is that MS2LDA requires frequent patterns of spectral features (fragments and neutral losses) extracted from raw MS/MS spectra to be manually annotated by MS experts. These two steps result in a set of annotated “Mass2Motifs” (M2Ms) that couple spectral features to descriptive sub-structures (e.g. “Amine loss - Indicative for free NH2 group in fragmented molecule”), comparable to MESSAR rules. Moreover, spectral feature–substructure associations in M2Ms are highly confident, and as such they can be used as a ground truth to assess the biochemical relevance of MESSAR rules.

We compared the 8378 MESSAR rules with the 500 positive ion mode M2Ms derived from GNPS [20]. Each M2M consists of up to 200 motif features (fragments and losses), sorted by their probabilities (Fig 3 left panel). The fifty most probable features were searched against the MESSAR target rules using a 20 ppm mass window around the MESSAR spectral features, and feature types were required to agree.

**Fig 3.**
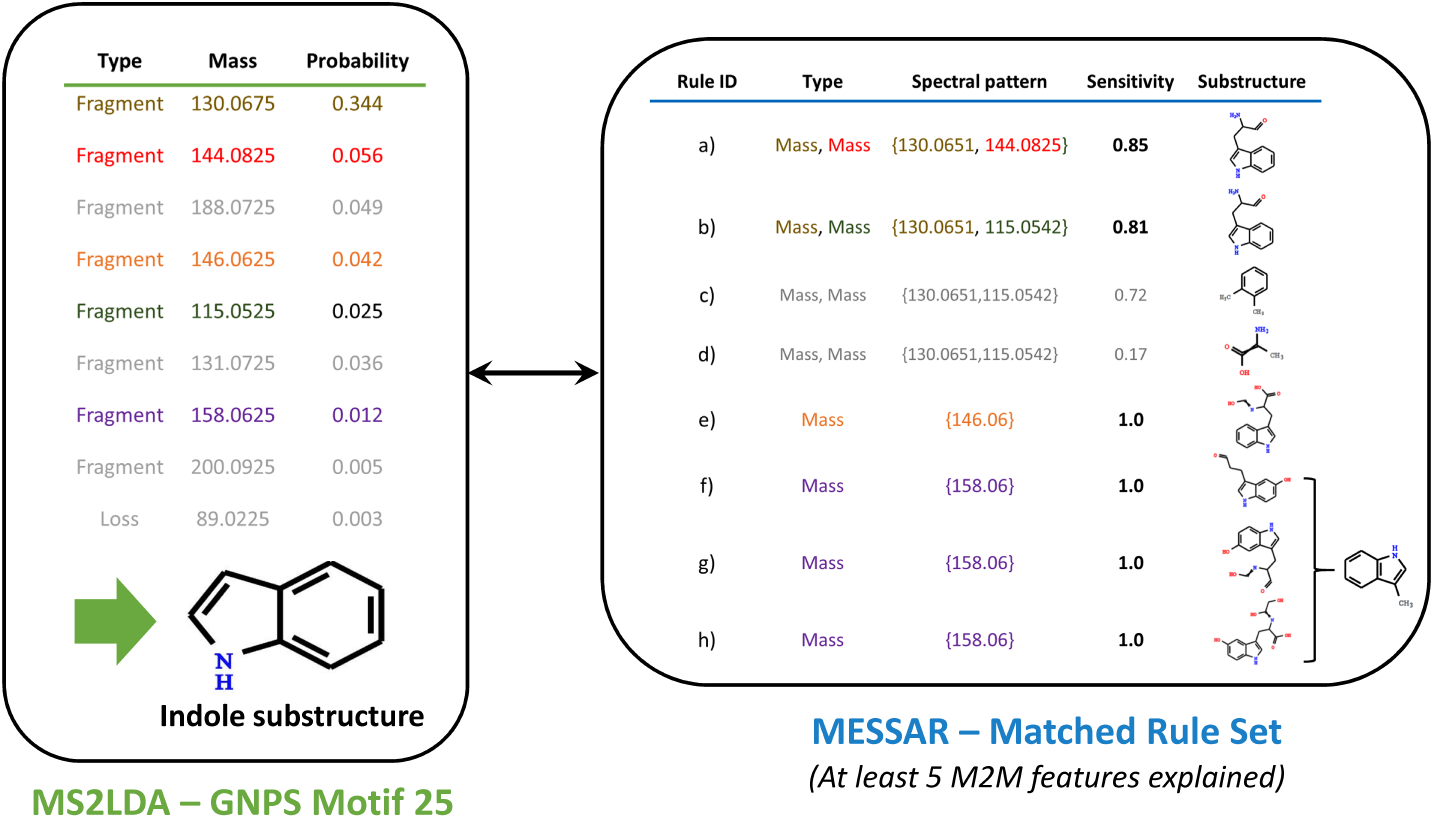
Example of comparison between Mass2Motifs and MESSAR rules. The fifty most probable features (fragments or losses) of the Motif 25 were searched against the MESSAR target rules. A 20 ppm mass window was used and feature types were required to agree (“Fragment” = “Mass”, “Loss” = “MDiff”). Sensitivity was used as the tie-breaker to select the most meaningful rules, and MCS was computed if multiple selected rules have the same sensitivity. The MESSAR substructure recommendations for Motif 25 were compared with the annotation by MS experts.

In practice, M2M features can never completely overlap with MESSAR rule features. One reason is that some M2M features can be isotopic peaks, noise,…, while such features were removed before training MESSAR rules. To understand what a “significant overlap” is, we performed two negative control experiments: i) searching “random motifs” consisting of only features taken from other motifs against MESSAR target rules; ii) searching motif features against 15480 unfiltered decoy rules. In i) and ii), 4 and 29 motifs overlapped with MESSAR rules by 5 or more features (or 10% for M2Ms with fewer than 50 features), respectively, while 77 actual M2Ms shared more than 5 common features with target rules (S2 File). Based on the negative control experiments, we can consider that there was a significant overlap between these 77 M2Ms and MESSAR rules.

We further investigated the underlying biochemical link based on these 77 significant M2M-rule matches. Briefly, we used MESSAR rules to annotate their matched motifs. The substructure annotation was achieved based on the set of MESSAR rules associated with a M2M via common spectral features (Fig 3). If several matched rules were associated with the same feature set, *sensitivity* was used as tiebreaker to select the most important rule. For instance, among rules b), c) and d) in Fig 3, rule b) with the highest *sensitivity* was selected, and only the substructure recommended by b) was kept for evaluation. If *sensitivity* could not break the tie (such as rules f, g and h in Fig 3), we reported the MCS of substructures predicted from all such rules. The MESSAR prediction for all 77 M2Ms can be found in S3 data.

Among these 77 M2Ms, 28 have also received expert annotations in [20]. Using these motifs, we validated the biochemical relevance of MESSAR rules by comparing substructures predicted with ground-truth expert annotation (S1 Table). Interestingly, the MESSAR-predicted substructures showed a striking similarity to expert knowledge, ranging from simple (e.g ethyl phenol of Motif 21) to complex (e.g. indole substructure of Motif 25, 26, 194) substructures. According to experts’ knowledge, ground-truth annotations of 26 motifs (out of 28) were identical or very similar to the substructure predicted by at least one matched MESSAR rule. On average, 40% of *sensitivity*-selected rules correctly predicted the motif substructure (exact number of correct rules in S2 File). In addition, matched rules can capture structural similarity between motifs. For example, most rules matching with motifs 1, 32, 39, 50 and 274, which are all steroid-related, correctly recommended the steroid core (S1 Table).

The MESSSAR rules also recommended substructures for the remaining 49 M2M (out of 77) that were not annotated in [20]. The examples in S2 Table and S3 Data show the ability of MESSAR to assign meaningful substructures, allowing biochemical interpretations of unknown M2Ms.

Overall, we found a significant overlap between 8378 MESSSAR rules and 77 M2Ms. The matched and *sensitivity*-selected rule set not only validated most expert annotations of M2Ms but also recommended substructures for unknown M2Ms. However, in broad terms, MESSAR and MS2LDA are strongly complementary, since rules and M2Ms show completely different formats: MESSAR rules link spectral features (with exact masses) with specific substructures, while M2Ms are spectral patterns derived from raw experimental spectra with descriptive substructure annotation (Fig 3). Although strong biochemical links were revealed from overlapping rules and motifs, the remaining rules and motifs can not be compared directly. We will further illustrate their complementarity in following sections.

### Validating MESSAR rules using authentic standards

MESSAR rules were characterized by their *sensitivity* calculated based on the entire training set. This metrics describes the probability that a feature set (*X*) is detected if the substructure (*Y*) is in the training molecule. Target-decoy comparison shows the ability of *sensitivity* to select non-random rules. In the MS2LDA-MESSAR comparison, rules with the highest *sensitivity* correctly predicted motif substructures from overlapped motif features. To demonstrate that the rest of rules were not only beyond random but also meaningful and reliable, 8378 MESSAR rules were individually validated on an independent test set of 5164 MASSBANK standards (S2 Data).

For each rule *X* ⇒ *Y*, we counted how many test spectra contain the feature set *X* while the substructure *Y* is present in the corresponding chemical standards, in other words, the number of true positives (Table 1). Consistent with the model training procedure, 4743 rules with *TP* ≥ 5 were statistically evaluated (S1 Text). We could not evaluate the rest of rules due to their rare occurrence in the test set. In addition to the *TP*s, we counted the frequency of substructure (*Supp*(*Y*)) for each rule, from which *sensitivity* was reported (Fig 4, S3 File).

**Fig 4.**
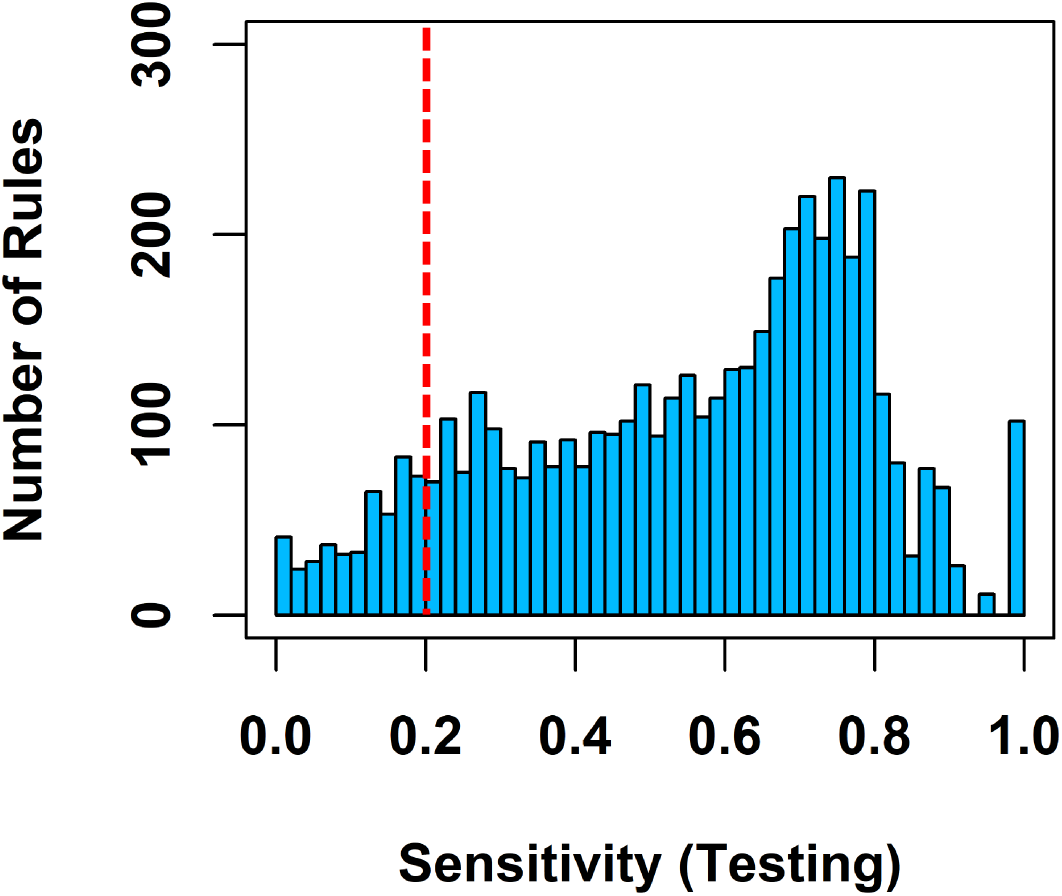
Statistical evaluation of MESSAR rules on testing data. Sensitivity distribution of 4743 evaluated rules. The red dash line indicates the 0.2 threshold used to define spurious rules.

**Table 1.** Confusion matrix for the MESSAR rule *X* → *Y*.

| | $Y$ | $\neg Y$ |
| --- | --- | --- |
| $X$ | True Positive (TP) | False Positive (FP) |
| $\neg X$ | False Negative (FN) | True Negative (TN) |

$Supp(X) = TP + FP$ , Number of spectra that contain the set of spectral feature(s) $X$ ; $Supp(Y) = TP + FN$ , Number of compounds that contain the substructure $Y$ .

**Table 2.**
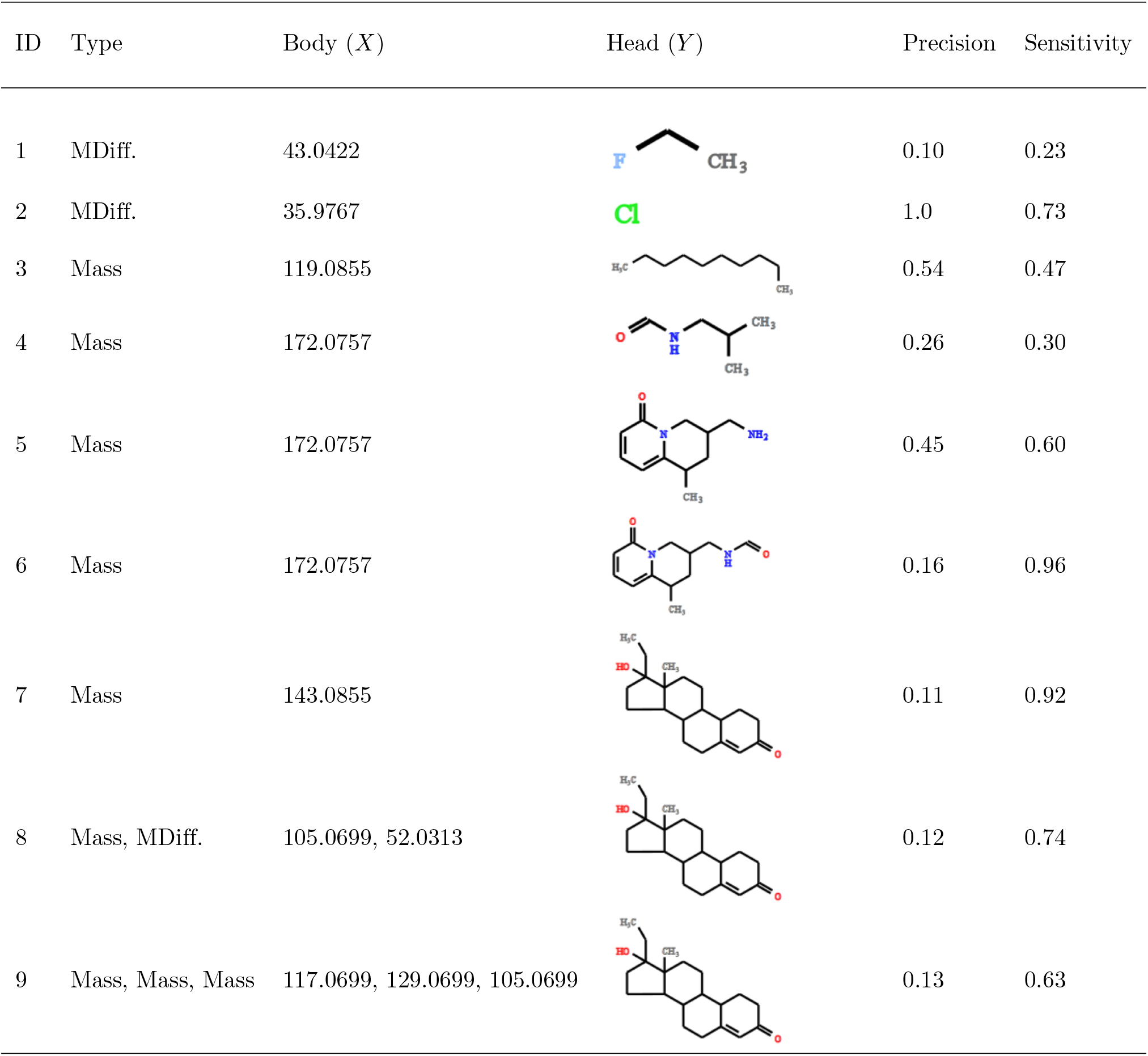
Examples of MESSAR rules from the target database.

| ID | Type | Body (X) | Head (Y) | Precision | Sensitivity |
| --- | --- | --- | --- | --- | --- |
| 1 | MDiff. | 43.0422 |  | 0.10 | 0.23 |
| 2 | MDiff. | 35.9767 |  | 1.0 | 0.73 |
| 3 | Mass | 119.0855 |  | 0.54 | 0.47 |
| 4 | Mass | 172.0757 |  | 0.26 | 0.30 |
| 5 | Mass | 172.0757 |  | 0.45 | 0.60 |
| 6 | Mass | 172.0757 |  | 0.16 | 0.96 |
| 7 | Mass | 143.0855 |  | 0.11 | 0.92 |
| 8 | Mass, MDiff. | 105.0699, 52.0313 |  | 0.12 | 0.74 |
| 9 | Mass, Mass, Mass | 117.0699, 129.0699, 105.0699 |  | 0.13 | 0.63 |

In Fig 4, half of evaluated rules (2364 out of 4743 rules) had a *sensitivity* higher than 0.6, while only 10% (463 out of 4743) might be spurious due to a *sensitivity* lower than 0.2. This result was not biased by the rule filtering step (Fig 1G) since we have only applied a 0.2 filter on training data-derived *sensitivity*. A global high *sensitivity* indicates that MESSAR rules are meaningful and powerful in capturing characteristic spectral features of a substructure. As an example, with a *sensitivity* of 1, the rule 21850 (S3 File) implies that the fragment 160.0757 appears in all testing compounds that involve cytisine substructure.

### Substructure recommendation for unknown spectra

We can use the entire rule set to annotate unknown spectra. This functionality is available to users in the MESSAR web tool. The intermediate steps of the substructure prediction procedure are illustrated in Fig 5A using the example of Ochratoxin B ethyl ester (Inchikey: XXAVUHHKDMGGBR-UHFFFAOYSA-N, Challenge ID: 38 in S2 Data “MASSBANK CASMI”). We queried the rule database based on the peaks and mass differences extracted from the test spectrum. With a 20 ppm mass window, the query resulted in 202 matched rules predicting 192 substructures. According to expert knowledge, most rules recommended benzene rings, aromatic amines and indole-related substrutures (S4 File). Such conclusion can be drawn after examining all matched rules. For ease of interpretation, we proposed an additional simplification step through analogous rules aggregation so that final recommended substructures were MCSs of rules head (*Y*). Accordingly, the score of each MCS was the sum of *sensitivity* of all responsible rules. We have aggregated 202 rules into 26 ranked substructures (S4 File). This last step, available in our web tool, is optional but strongly recommended.

**Fig 5.**
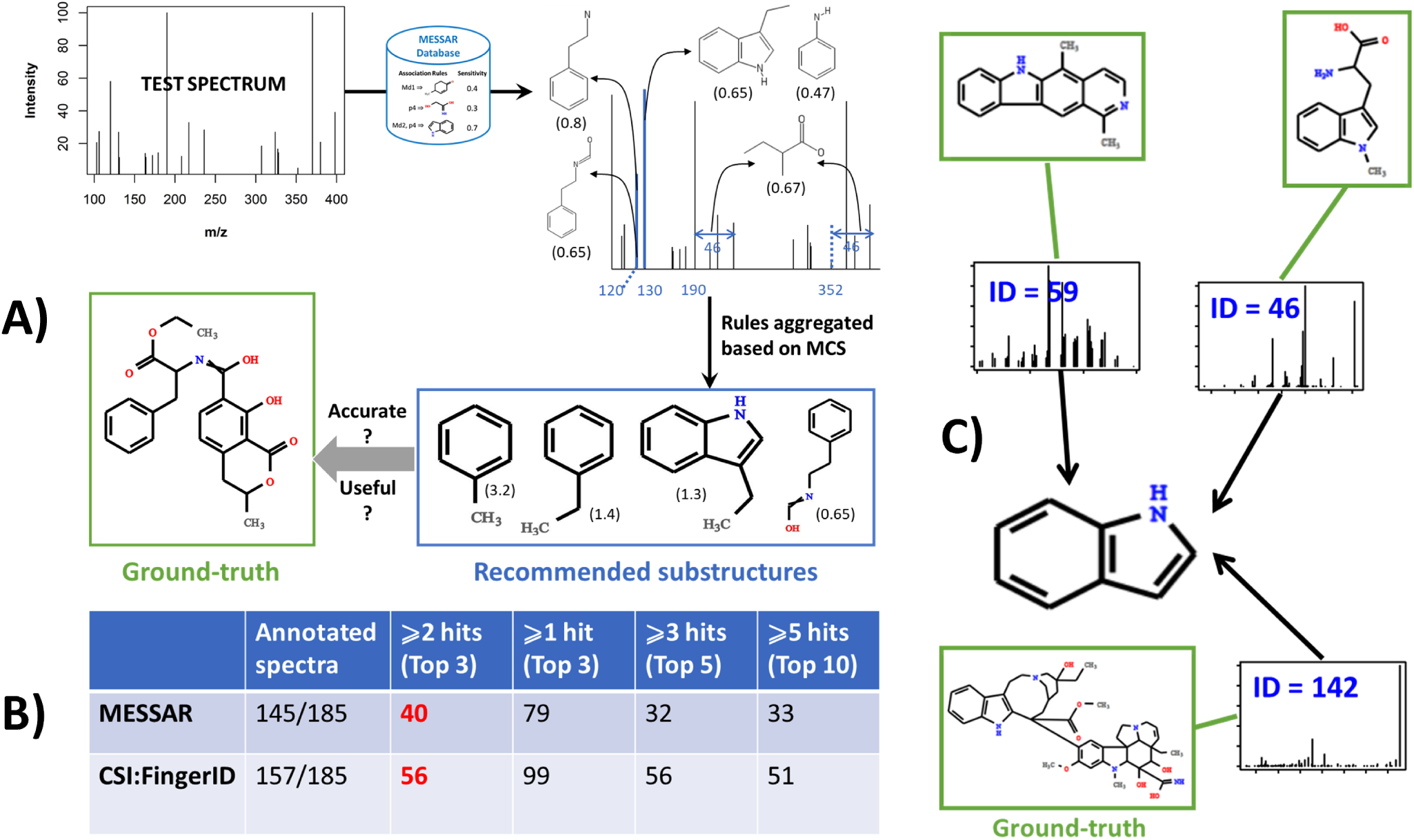
Substructure recommendation and interpretation for unknown spectra. A) Spectral features of the test spectrum are searched against the rule database. The matched rules are aggregated through MCS calculation of analogous rules. The final recommendations are MCSs scored by the sum of *sensitivity*. The top recommendations are verified against the ground-truth in terms of exact structural match and biochemical relevance. B) Number of test spectra annotated by MESSAR or CSI:FingerID (since some spectra cannot be processed by either tool) and number of “good annotations” under four criteria of accuracy. C) Based on top 3 outputs of MESSAR, test spectra 46, 59 and 142 can be grouped for sharing potentially the indole substructure.

In practice, only top-ranked substructures are recommended to end users for *de novo* identification or approximate characterization of unknown metabolites. In other words, the top candidates of a good substructure prediction tool should contain both accurate and meaningful (e.g. biochemically-relevant) structural knowledge. Therefore, both quantitative (accuracy of tools) and qualitative (meaningfulness) aspects were considered when comparing MESSAR output (top-ranked substructures aggregated from matched rules) with two other substructure recommendation tools i.e. CSI:FingerID and MS2LDA.

The 185 test spectra from “MASSBANK CASMI” (S2 Data) were submitted to all three software. The accuracy of MESSAR was only compared with CSI:FingerID since MS2LDA output was text descriptions of substructures thus not eligible for quantitative comparison. Small substructures with fewer than 5 non-hydrogen atoms were discarded. Based on the output of both tools, we retrieved the number of hits (when the predicted substructure is an exact match of ground-truth) among top 3, 5 and 10 recommended substructures. We counted the number of “properly annotated” test spectra under four fixed criteria (Fig 5B): i) at least one hit among top 3; ii) 2 or 3 hits among top 3; iii) at least 3 hits among top 5 and iv) at least 5 hits among top 10 candidates (Fig 5B). These criteria reflect how trustworthy the tools are if users only look at the top substructure recommendations. Among spectra that CSI:FingerID or MESSAR were able to process, around 30% and 60% were correctly annotated by both software according to the first and second criterion, respectively. In other words, 60% of spectra received at least one exact substructure annotation by taking top 3 substructures from either tool.

Based on independent spectra, CSI:FingerID had a slightly better performance over MESSAR in terms of accuracy. However, an exact substructure match to test compounds does not mean that the output is meaningful or useful to reveal the ground-truth. For instance, a substructure with the SMILES code “CC(CCC)CC” is probably an exact match to diverse metabolites, but it does not help understanding the biochemical origin of unknowns. Therefore, a qualitative evaluation of software output by an external expert is preferred. Here we collected the output of all three software for the 185 spectra as well as the ground-truth in S3 Table. MESSAR and CSI:FingerID outputs were top 3 substructure candidates, and MS2LDA output was the interpretation of matched GNPS motif. The external expert performed a blinded evaluation and suggested for each spectrum the appropriate tool(s) based on: i) how useful the predicted substructures are (e.g. indication of chemical family); ii) biochemical relevance of output with regard to ground-truth. In the example of Ochratoxin B ethyl ester, MESSAR and CSI:FingerID were equally considered useful since both correctly predicted the aromatic ring and amine group (S3 Table).

According to the expert, 126 out of 185 test spectra received reliable and useful annotations from at least one software (S3 Table). MESSAR, CSI:FingerID and MS2LDA reliably predicted substructures for 68, 65 and 32 times, and they were single appropriate tool for 41, 41 and 6 test spectra, respectively. These comparisons indicate that all three tools could add value to a meaningful substructure prediction. Specifically, MESSAR was most powerful for capturing polycyclic aromatic (e.g. Challenge ID: 3,9,16,36,58,78...), indole (ID: 6, 46, 59, 60, 142...) and chlorobenzene (ID: 40, 66) substructures. CSI:FingerID provided reliable prediction of amino acids (ID: 25, 41, 119, 153, 179) and other nitrogen-containing functional groups such as pyrimidine (ID: 13, 94) and benzenesulfonylamide (93, 182). MS2LDA was able to elucidate sterone-related (ID: 3, 36, 84, 146) and conjugated sugar (ID: 76, 169). Combined use of three tools would allow a broader coverage of metabolite families and increased reliability of substructure prediction. On the other hand, one tool can be more preferable if user has prior knowledge about biochemical origins of the unknown.

## Conclusions

MESSAR was inspired by the idea that the presence of spectral features in an MS/MS spectrum is linked to substructures of the metabolite. We have implemented a data-driven approach to unravel such relations from public spectral library. Our approach was inspired from association rule mining. Statistical evaluation of target/decoy rules and validation on independent spectra characterized such relations as “good *sensitivity* but lower *precision*”. It means that some spectral features are constantly present for compounds containing a certain substructure. In fact, rules with high *sensitivity* in S1 File reveal characteristic ions for several important substructure. However, learning a specific substructures from one or a few spectral features could be challenging due to the low *precision*.

Although individual MESSAR rules have a low predictive power in terms of *precision*, the strength of our approach lies in two aspects: i) rules with higher *sensitivity* predict meaningful substructures, useful for *de novo* identification; ii) the sparsity of rules enables the prediction of diverse substructures, making our tool a good structural hypothesis generator. On the other hand, rules matched to an unknown spectrum can usually be aggregated through MCS search, leading to accurate and reliable substructure recommendation.

We developed MESSAR web tool to assist the *de novo* annotation of unknown metabolites, for example, to identify functional classes of unknown spectra that share substructures (Fig 5C), to corroborate results from other chemical identification tools, etc. Our tool and CSI:FingerID work on a similar scope, but the machine-learning model behind is fundamentally different. First, CSI:FingerID starts by converting the training spectra into fragmentation trees before learning implicit rules between fragmentation tree similarity and substructure presence. In contrast, MESSAR directly explores the relationships between mass features and molecular substructures. Second, CSI:FingerID relies on molecular fingerprints to train SVM (support vector machine) models, while MESSAR employs both predefined substructures (molecular fingerprints) and less common ones by breaking chemical bonds of training compounds. Third, the main objective of CSI:FingerID is to identify compounds via candidate reranking, whereas MESSAR focuses on the partial identification of unknowns via substructure prediction. Therefore, one tool might be more suitable than the other depending on the type of compounds, and a joint use is recommended.

MESSAR is inherently complementary to recently-published software MS2LDA as MS2LDA extracts co-occurring spectral features while MESSAR provides an automated structural annotation of these features. We envisage an improvement in substructure discovery by the combined use of expert-driven M2M annotations and data-driven MESSAR rules. Similarly, we anticipate that MESSAR will be useful for the functional analysis of complex biological matrices as it can quickly recognize substructure patterns.

## Supporting information

**S1 Text. Supporting method.**

**S1 Fig. Supplementary statistical evaluation of rules.** A) Distribution of target/decoy rule *specificity*. B) Dependency between *sensitivity* of target rules and *support*.

**S2 Fig. Examples of GNPS training spectra** The peak 172.075 is present in all examples, while the underlying training molecules were structurally similar – little or no substructure difference was observed.

**S3 Fig. Filtering target rules via FDR control** A) *Sensitivity* distribution of target and decoy rules. B) FDR score of rules as a function *sensitivity*. The red vertical line in both plots indicates the *sensitivity* threshold estimated for 1% FDR.

**S1 Table. Annotation of 26 M2M motifs by MESSAR rules and comparison with ground-truth**

**S2 Table. Examples of unexplained positive ion mode M2Ms annotated by MESSAR rules.**

**S3 Table. Direct comparison between MESSAR, CSI:FingerID and MS2LDA for 185 testing spectra, including ground-truth and expert’s decision of meaningful output**

**S1 Data. MESSAR training data and molecular fingerprints used for substructure generation** GNPS spectral library and its decoy version in mgf format. The datasets are derived from https://bio.informatik.uni-jena.de/passatutto. The molecular fingerprints are retrieved from the SIRIUS-4.0.1.

**S2 Data. MESSAR testing data**

**S3 Data. MESSAR rules that significant overlapped with M2M motifs**

**S1 File. FDR-filtered target MESSAR rule database and statistical metrics based on training data**

**S2 File. Overlapping statistics between rules and motifs** i) target MESSAR rules and motifs, ii) target MESSAR rules and “random” motifs, iii) decoy MESSAR rules and motifs.

**S3 File. Statistical evaluation of MESSAR rules on independent testing data**

**S4 File. MESSAR substructure recommendation for a testing spectrum** i) all matched rules sorted by *sensitivity*. ii) rules were aggregated and obtained MCSs were scored and ranked.

## Data availability

The source code is publicly available: https://github.com/daniellyz/MESSAR.

## Acknowledgments

Special thanks goes to Sara Forcisi for evaluating the output of substructure recommendations. We also thank Danh Bui-Thi and Charlie Beirnaert for insightful discussions.

