## Supplementary material for "MESSAR: Automated recommendation of metabolite substructures from tandem mass spectra": S3 Table

**Table S3** Substructure prediction by MESSAR, CSI:FingerID and MS2LDA for 185 challenge spectra along with the ground-truth. The external expert decided for each challenge the meaningful and relevant substructure(s) compared to the ground-truth (without knowing the name of software used to generate each substructure). The selected substructures were inside blue rectangles. We report for each challenge the tool(s) that generate the selected substructure(s).

*Example:*

| ID | MESSAR | CSI:FingerID | MS2LDA | Ground-truth | Selected method |
|---|---|---|---|---|---|
| 38 | 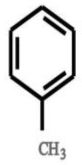 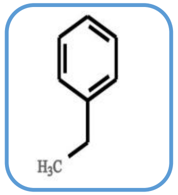 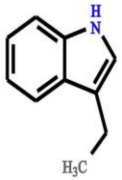 | 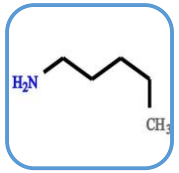 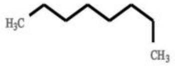 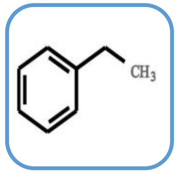 | CO loss - indicative for presence of ketone/aldehyde/lactone group (C=O) | 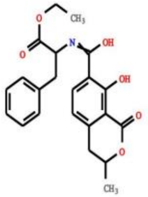 | MESSAR & CSI:FingerID |

| | | | | | |
|---|---|---|---|---|---|
| 1 | 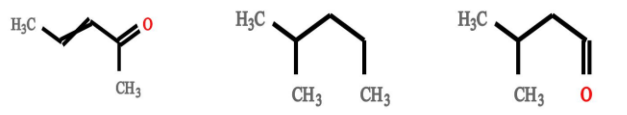 | 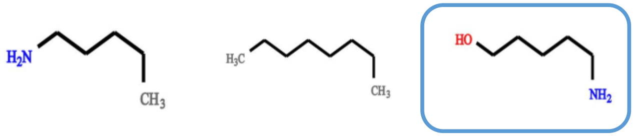 | None | 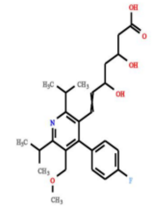 | CSI:FingerID |
| 2 | 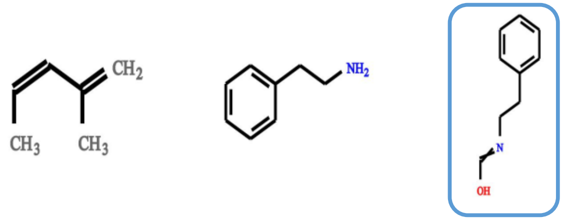 | 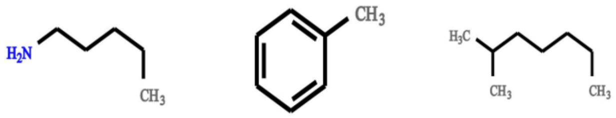 | Fragment indicative for aromatic compounds related to methylbenzene substructure ($C_7H_7$ fragment) | 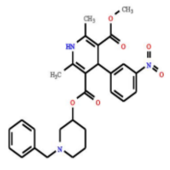 | MESSAR |
| 3 | 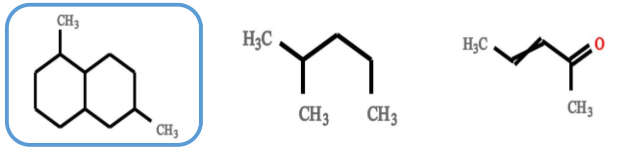 | 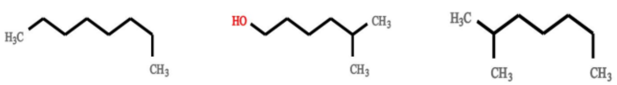 | Steroid core related ($C_{18}H_{21}$ and smaller fragments thereof - with $C_{12}H_{13}$, $C_{11}H_{11}$, and $C_{11}H_{13}$ most probable) | 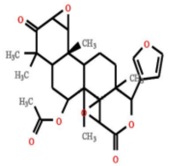 | MESSAR & MS2LDA |
| 4 | 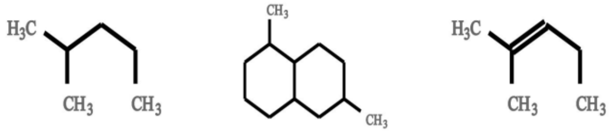 | 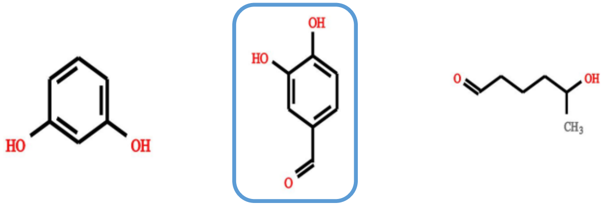 | CO loss - indicative for presence of ketone/aldehyde/lactone group (C=O) | 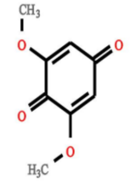 | CSI:FingerID |
| 5 | 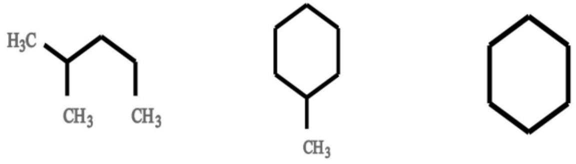 | None None None | None | 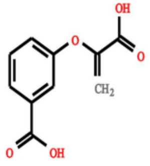 | None |

| | | | | | | | | | |
|---|---|---|---|---|---|---|---|---|---|
| 6 | 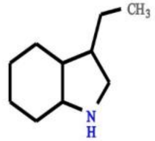 | 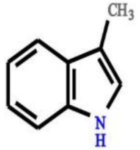 | 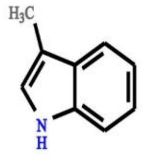 | 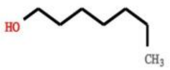 | 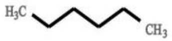 | 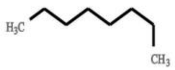 | [illegible][7.3.1.0~2,7~]trideca-2,4 substructure | 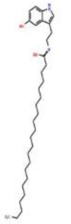 | MESSAR |
| 7 | 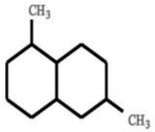 | 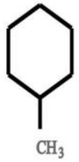 | 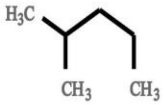 | 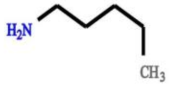 | 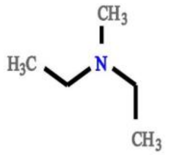 | 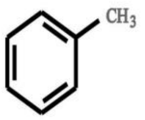 | Fragment indicative for aromatic compounds related to methylbenzene substructure (C7H7 fragment) | 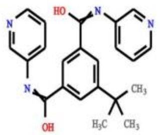 | CSI:FingerID |
| 8 | None | None | None | None | None | None | None | 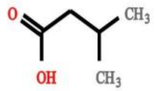 | None |
| 9 | 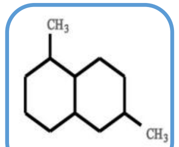 | 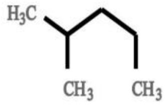 | 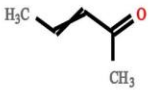 | 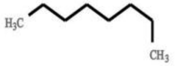 | 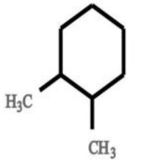 | 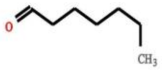 | Water loss - indicative of a free hydroxyl group â€“ (in beer often seen in sugary structures) | 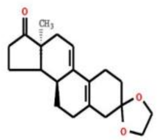 | MESSAR |
| 10 | 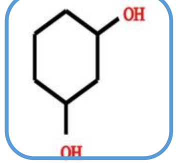 | 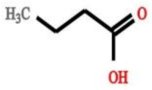 | 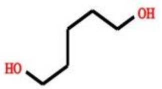 | 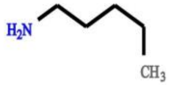 | 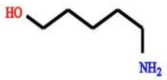 | 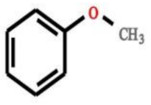 | None | 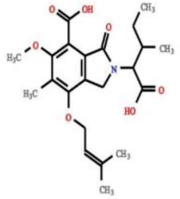 | MESSAR<br>&CSI:FingerID |

| | | | | | | | | | |
|---|---|---|---|---|---|---|---|---|---|
| 11 | 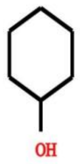 | 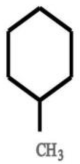 | 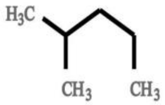 | 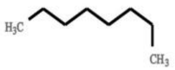 | 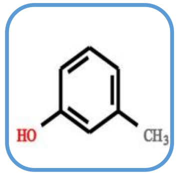 | 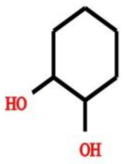 | None | 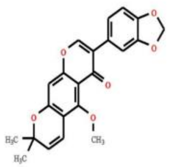 | CSI:FingerID |
| 12 | 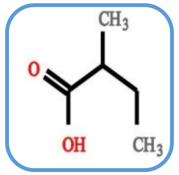 | 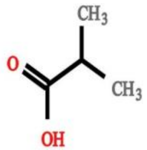 | None | 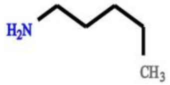 | 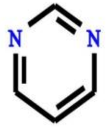 | None | None | 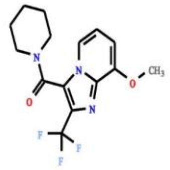 | MESSAR |
| 13 | 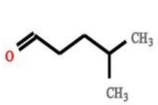 | 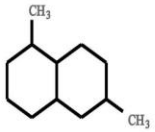 | 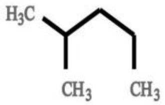 | 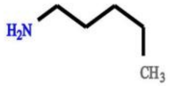 | 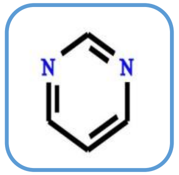 | 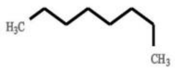 | None | 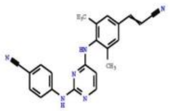 | CSI:FingerID |
| 14 | None | None | None | 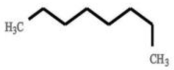 | 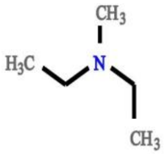 | 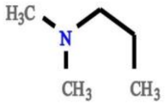 | 4-keto-chlorobenzene substructure | 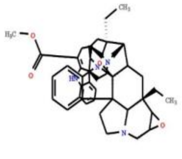 | None |
| 15 | 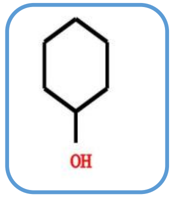 | 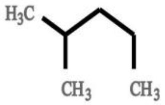 | 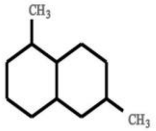 | 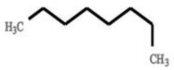 | 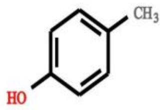 | 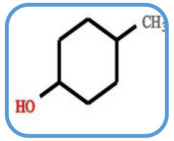 | Fragments indicative for cinnamic/hydroxycinnamic acid substructure | 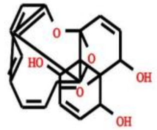 | MESSAR &CSI:FingerID |

| | | | | | | | | |
|---|---|---|---|---|---|---|---|---|
| 16 | 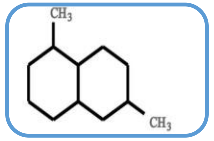 | 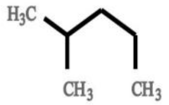 | 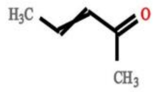 | 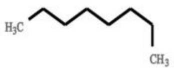 | 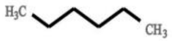 | None | None | 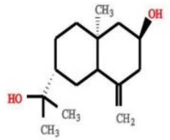 | MESSAR |
| 17 | None | None | None | 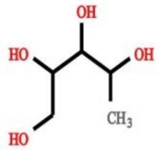 | 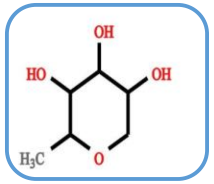 | 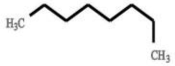 | Methoxylated benzene substructure [ClassyFire - Relevant terms - Substituents: Methoxybenzene, Phenoxy compounds - Taxa: O-methylated flavonoids] | 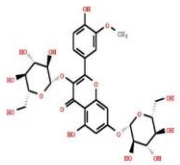 | CSI:FingerID &MS2LDA |
| 18 | 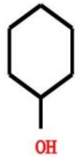 | 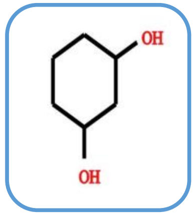 | 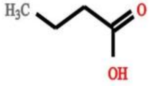 | 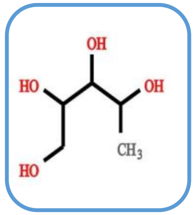 | 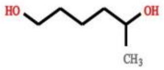 | 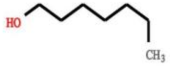 | phthalate substructure | 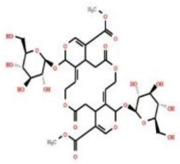 | MESSAR &CSI:FingerID |
| 19 | 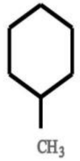 | 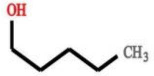 | 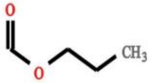 | None | None | None | xyphenyl)-2-oxo-2H-chrom relared substructure (or isomeric variants) | 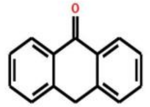 | None |
| 20 | 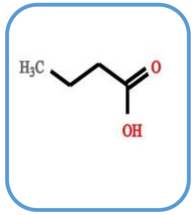 | 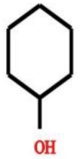 | 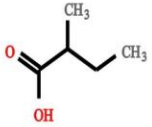 | 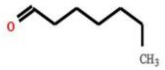 | 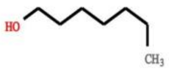 | 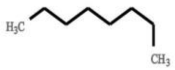 | Fragments indicative for cinnamic/hydroxycinnamic acid substructure | 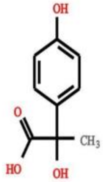 | MESSAR &MS2LDA |

| | | | | | | | | | |
|---|---|---|---|---|---|---|---|---|---|
| 21 | 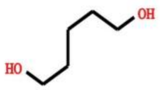 | 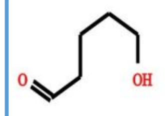 | 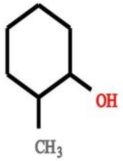 | 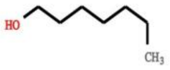 | 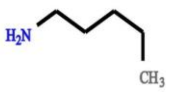 | 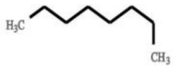 | None | 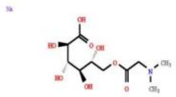 | MESSAR &CSI:FingerID |
| 22 | 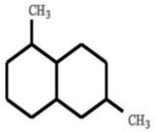 | 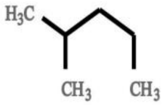 | 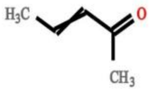 | 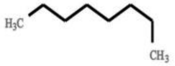 | 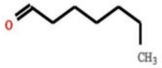 | 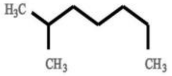 | None | 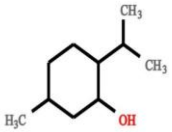 | None |
| 23 | 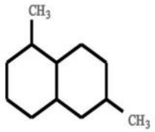 | 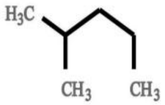 | 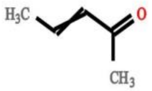 | 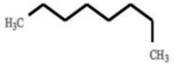 | 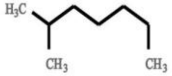 | None | Steroid core related (C18H21 and smaller fragments thereof - with C12H13, C11H11, and C11H13 most probable) | 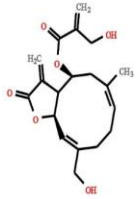 | MESSAR |
| 24 | 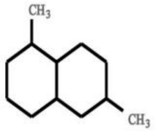 | 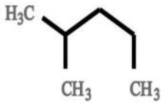 | 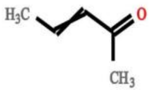 | 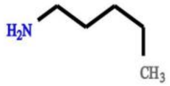 | 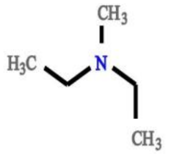 | 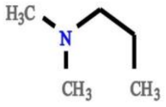 | None | 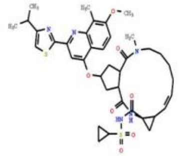 | CSI:FingerID |
| 25 | 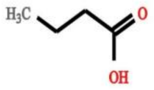 | 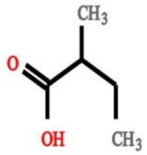 | 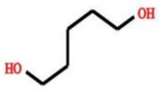 | 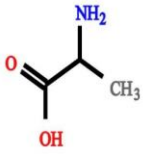 | 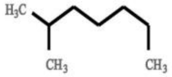 | 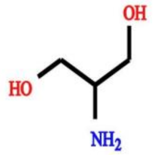 | None | 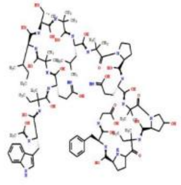 | None |

| | | | | | | | | | |
|---|---|---|---|---|---|---|---|---|---|
| 26 | 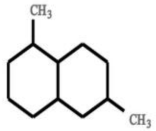 | 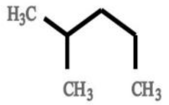 | 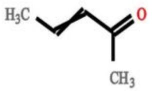 | 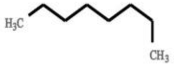 | 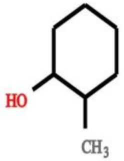 | 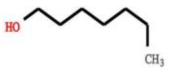 | 2-oxochromen-7-yl [mainly trimethylated] related substructure | 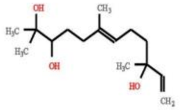 | MESSAR |
| 27 | None | None | None | None | None | None | cinchonan-9-ol substructure | 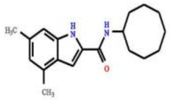 | MS2LDA |
| 28 | 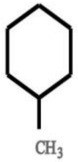 | 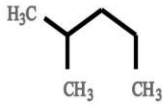 | 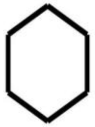 | 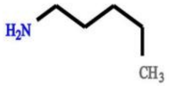 | 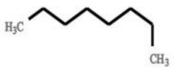 | 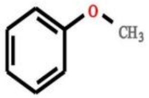 | Sterone related | 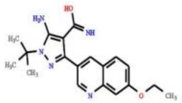 | CSI:FingerID |
| 29 | 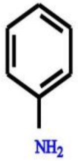 | 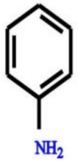 | 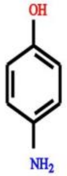 | 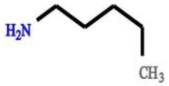 | None | None | None | 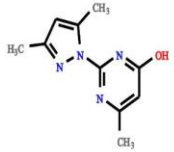 | None |
| 30 | 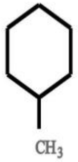 | 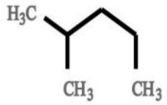 | 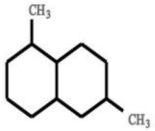 | 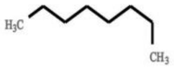 | 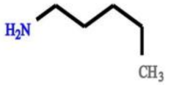 | None | None | 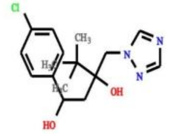 | None |

| | | | | | | | | | |
|---|---|---|---|---|---|---|---|---|---|
| 31 | 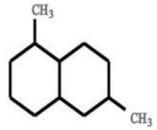 | 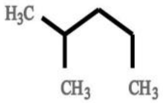 | 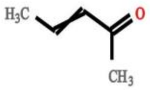 | 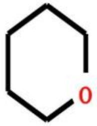 | 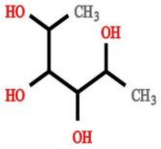 | 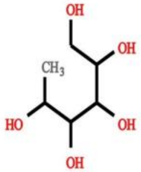 | Fragments indicative of a glycosylation â€“ i.e. indicative for a sugar conjugation (in beer often related to glucose) | 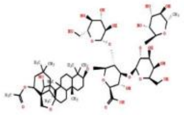 | CSI:FingerID &MS2LDA |
| 32 | 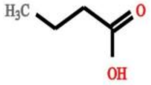 | 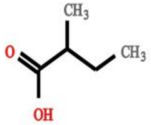 | 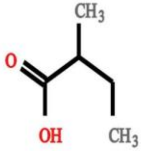 | 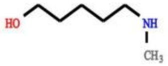 | 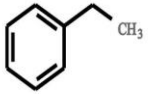 | 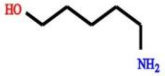 | None | 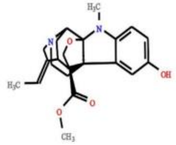 | None |
| 33 | 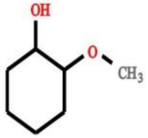 | 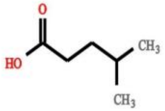 | 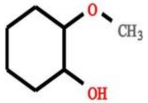 | 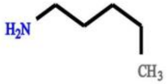 | None | None | None | 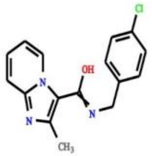 | None |
| 34 | None | None | None | None | None | None | None | 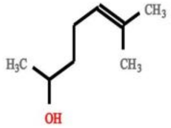 | None |
| 35 | None | None | None | None | None | None | Aliphatic amine (NH3 loss indicates free NH2 group coupled to aliphatic chain) | 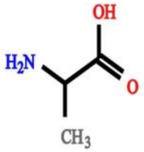 | MS2LDA |

| | | | | | | | | | |
|---|---|---|---|---|---|---|---|---|---|
| 36 | 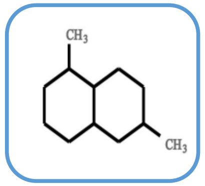 | 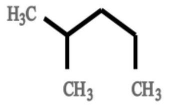 | 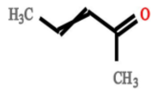 | 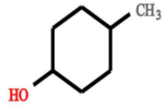 | 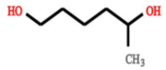 | 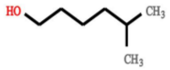 | Hydroxy-Pregnenolone related fragments (not very specific - indicative for presence of steroid backbone) | 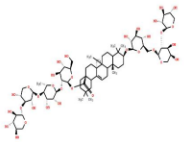 | MESSAR &MS2LDA |
| 37 | 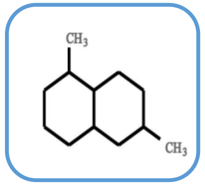 | 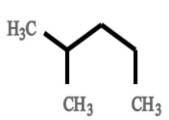 | 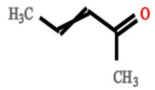 | 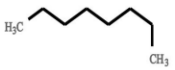 | 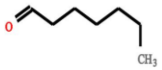 | None | Sterone related | 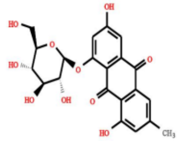 | MESSAR &MS2LDA |
| 38 | 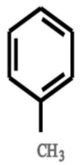 | 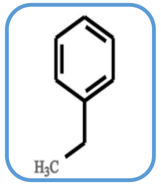 | 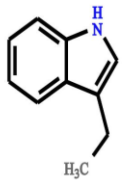 | 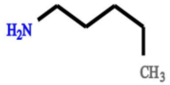 | 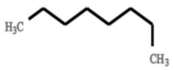 | 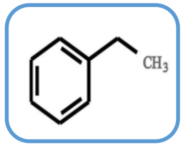 | CO loss - indicative for presence of ketone/aldehyde/lactone group (C=O) | 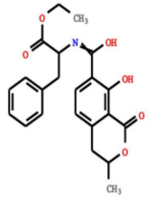 | MESSAR &CSI:FingerID |
| 39 | 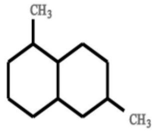 | 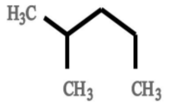 | 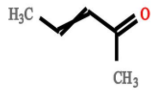 | 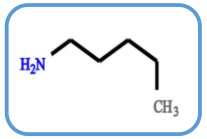 | 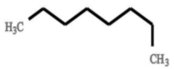 | 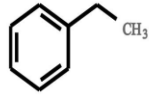 | nitrogen containing substructure [C5H12N] (in beer related to Leucine) | 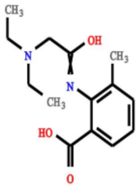 | CSI:FingerID &MS2LDA |
| 40 | 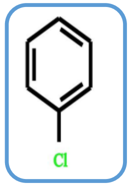 | 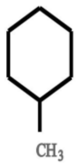 | 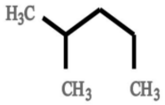 | 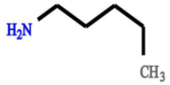 | None | None | CO loss - indicative for presence of ketone/aldehyde/lactone group (C=O) | 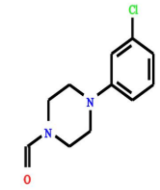 | MESSAR |

| | | | | | | | | | |
|---|---|---|---|---|---|---|---|---|---|
| 41 | 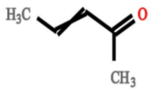 | 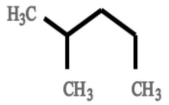 | 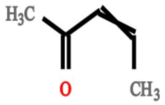 | 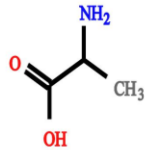 | 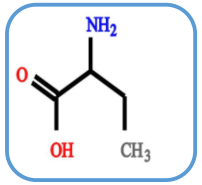 | 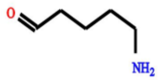 | Fragment and loss of [proline-H2O] - indicative for conjugated proline or arginine/ornithine â€“ EF fits | 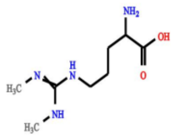 | CSI:FingerID & MS2LDA |
| 42 | None | None | None | 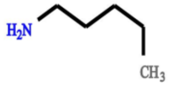 | 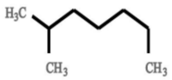 | None | [illegible] substructure (or formed after fragmentation event of dihydro analogue) [ClassyFire: Relevant terms - Substituents: Quinoxaline - [illegible] | 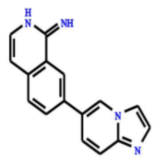 | MS2LDA |
| 43 | 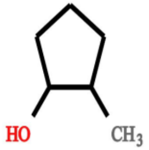 | 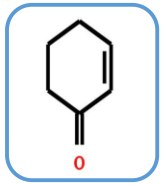 | 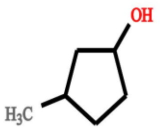 | 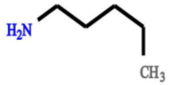 | 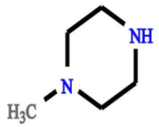 | 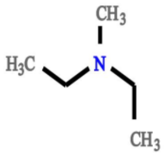 | None | 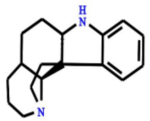 | MESSAR |
| 44 | 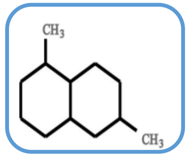 | 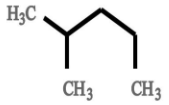 | 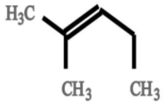 | 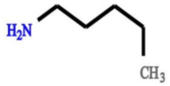 | 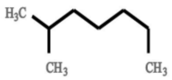 | 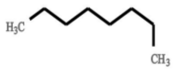 | None | 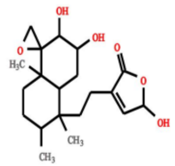 | MESSAR |
| 45 | 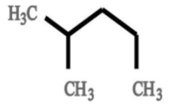 | 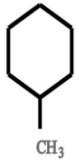 | 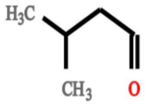 | 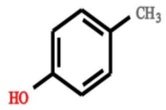 | 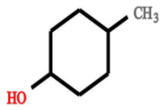 | 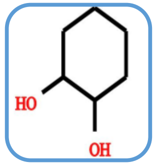 | Fragments indicative for ferulic acid based substructure (MzCloud) | 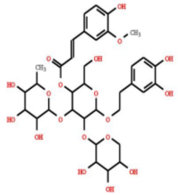 | CSI:FingerID & MS2LDA |

| | | | | | | | | | |
|---|---|---|---|---|---|---|---|---|---|
| 46 | 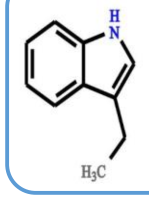 | 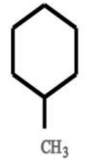 | 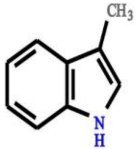 | 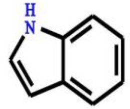 | 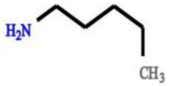 | 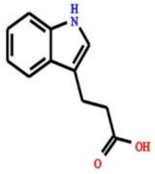 | Amine loss - Indicative for free NH2 group in fragmented molecule | 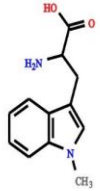 | MESSAR & CSI:FingerID |
| 47 | 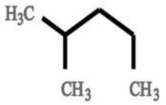 | 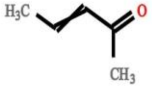 | 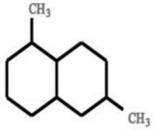 | 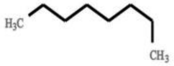 | 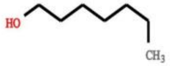 | None | None | 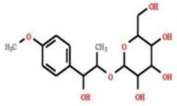 | None |
| 48 | None | None | None | 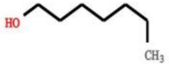 | 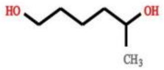 | 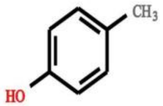 | None | 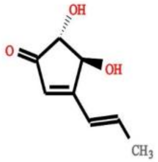 | None |
| 49 | 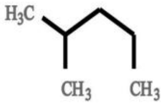 | 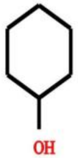 | 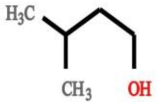 | 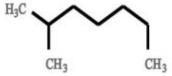 | 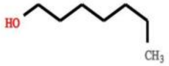 | 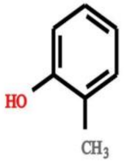 | 2-oxochromen-7-yl [mainly dimethylated] related substructure | 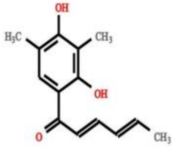 | CSI:FingerID |
| 50 | 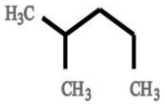 | 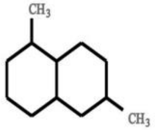 | 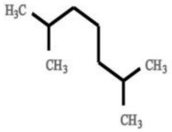 | 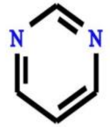 | 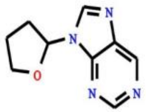 | 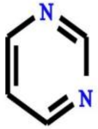 | Fragments indicative adenine (C5H6N5) substructure â€“ most prevalent in Beer3 | 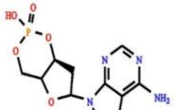 | CSI:FingerID |

| | | | | | | | | | |
|---|---|---|---|---|---|---|---|---|---|
| 51 | 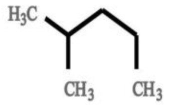 | 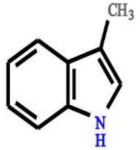 | 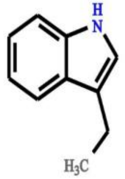 | 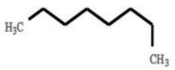 | 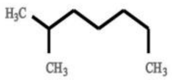 | 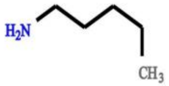 | None | 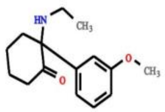 | None |
| 52 | 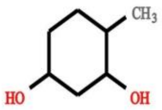 | 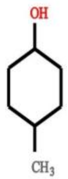 | 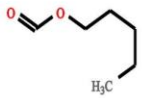 | None | None | None | Small nitrogen containing fragment ion â€“ often proline or ornithine derived â€“ most abundant fragment in beer data | 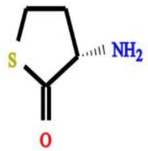 | None |
| 53 | None | None | None | 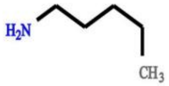 | 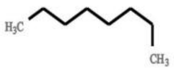 | 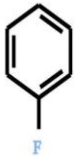 | None | 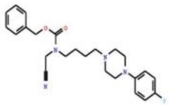 | CSI: FingerID |
| 54 | 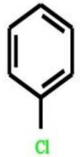 | 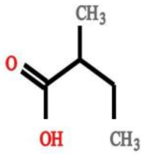 | 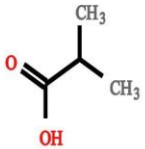 | None | None | None | Water loss - indicative of a free hydroxyl group â€“ (in beer often seen in sugary structures) | 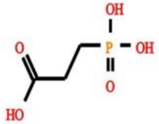 | None |
| 55 | 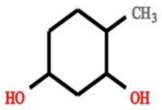 | 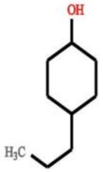 | 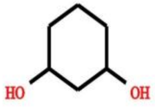 | 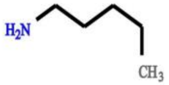 | 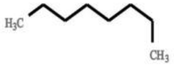 | None | nitrogen containing substructure [C5H12N] (in beer related to Leucine) | 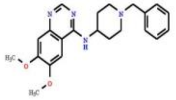 | MESSAR |

| | | | | | | | | |
|---|---|---|---|---|---|---|---|---|
| 56 | 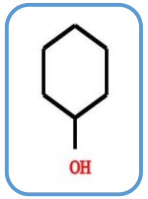 | 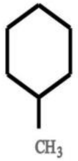 | 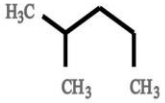 | 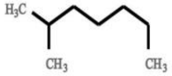 | None | None | None | 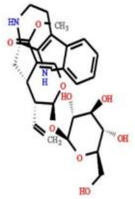 | MESSAR |
| 57 | 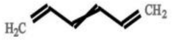 | 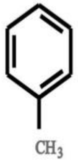 | 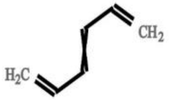 | None | None | None | None | 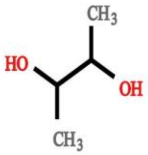 | None |
| 58 | 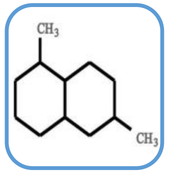 | 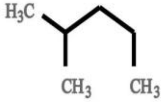 | 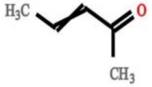 | 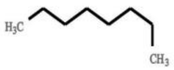 | 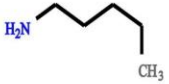 | 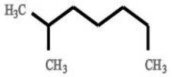 | Steroid core related (C18H21 and smaller fragments thereof - with C12H13, C11H11, and C11H13 most probable) | 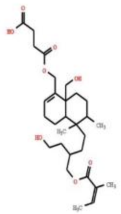 | MESSAR & MS2LDA |
| 59 | 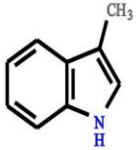 | 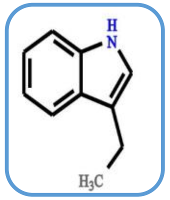 | 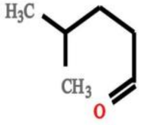 | 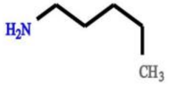 | 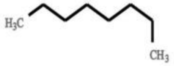 | 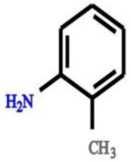 | None | 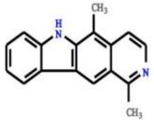 | MESSAR |
| 60 | 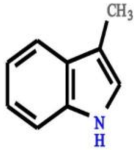 | 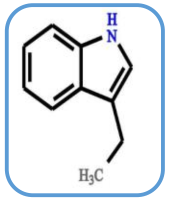 | 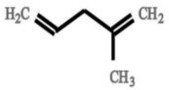 | 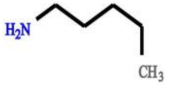 | 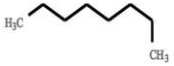 | 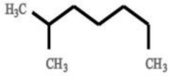 | C4H8 loss indicative for saturated C4-alkyl substructure (often tert-butyl group or loss from 9,10-dihydro-2H,8H-pyran substructure) | 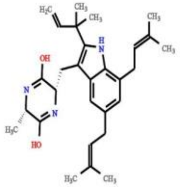 | MESSAR |

| | | | | | | | | | |
|---|---|---|---|---|---|---|---|---|---|
| 61 | 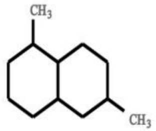 | 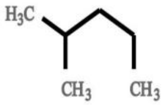 | 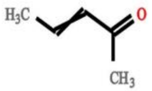 | 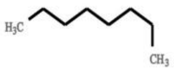 | None | None | None | 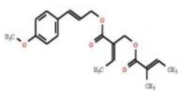 | None |
| 62 | 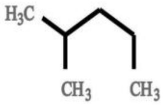 | 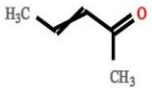 | 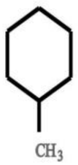 | None | None | None | None | 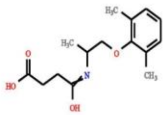 | None |
| 63 | None | None | None | 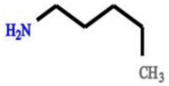 | 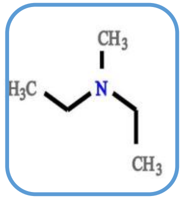 | 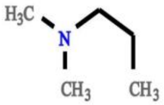 | Fragment ions indicative for alkylamine substructure C5H10N (in beer often pipecolic acid [pipecolate]) | 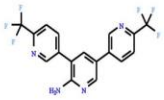 | CSI:FingerID |
| 64 | 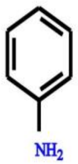 | 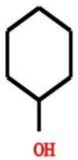 | 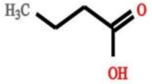 | 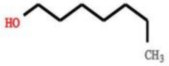 | 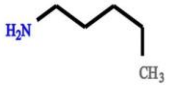 | 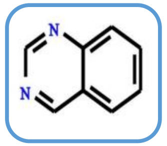 | Amine loss - Indicative for free NH2 group in fragmented molecule | 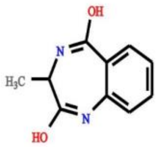 | CSI:FingerID |
| 65 | 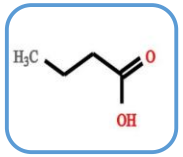 | 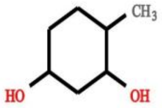 | 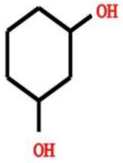 | 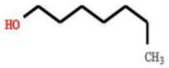 | 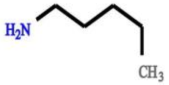 | 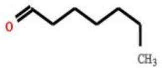 | Loss of CH2O2 - indicative for underivatized carboxilic acid group | 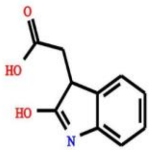 | MESSAR |

| | | | | | | | | | |
|---|---|---|---|---|---|---|---|---|---|
| 66 | 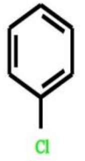 | 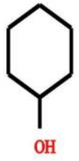 | 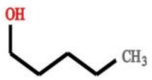 | 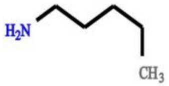 | None | None | indicative of piperazine substructure (or related N-containing ring structures for C2H5N loss only) | 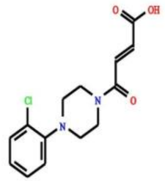 | MESSAR & MS2LDA |
| 67 | 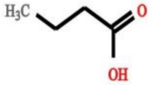 | 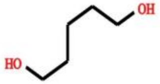 | 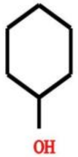 | 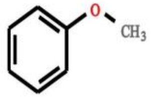 | 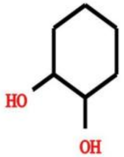 | 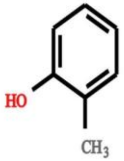 | Methoxylated benzene substructure [ClassyFire - Relevant terms - Substituents: Methoxybenzene, Phenoxy compounds - Taxa: O-methylated flavonoids] | 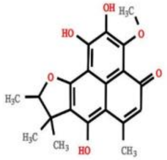 | CSI:FingerID |
| 68 | 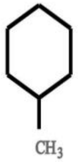 | 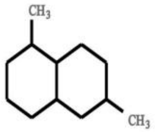 | 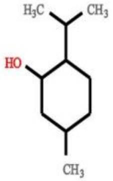 | 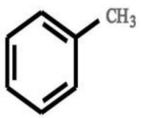 | 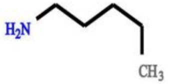 | 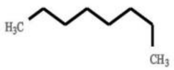 | Fragment indicative for aromatic compounds related to methylbenzene substructure (C7H7 fragment) | 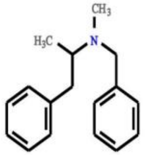 | CSI:FingerID |
| 69 | None | None | None | None | None | None | None | 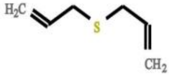 | None |
| 70 | 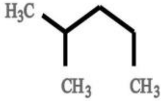 | 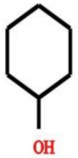 | 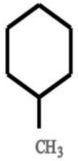 | 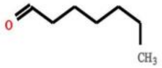 | 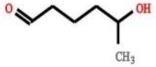 | 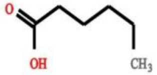 | CO loss - indicative for presence of ketone/aldehyde/lactone group (C=O) | 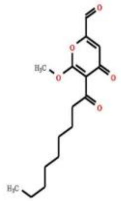 | MESSAR |

| | | | | | | | | | |
|---|---|---|---|---|---|---|---|---|---|
| 71 | 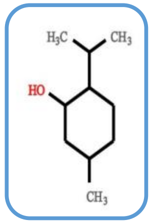 | 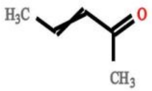 | 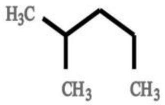 | 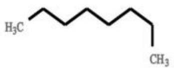 | 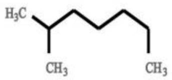 | 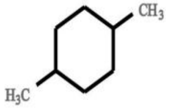 | None | 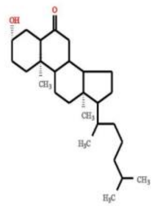 | MESSAR |
| 72 | 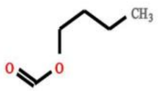 | 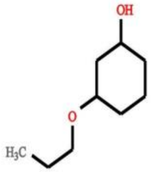 | 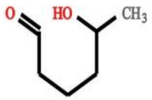 | 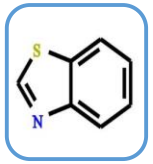 | 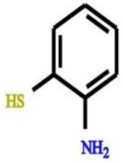 | 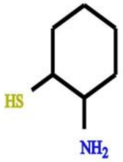 | None | 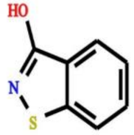 | CSI:FingerID |
| 73 | 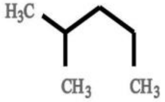 | 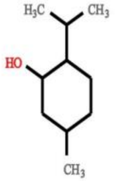 | 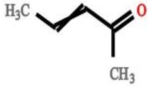 | 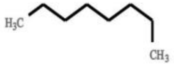 | 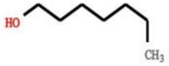 | 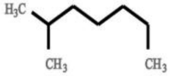 | None | 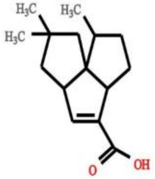 | None |
| 74 | None | None | None | 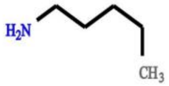 | 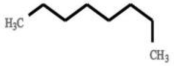 | 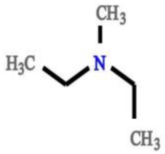 | None | 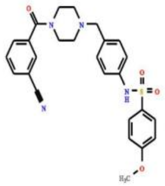 | None |
| 75 | 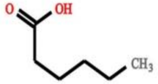 | 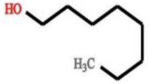 | 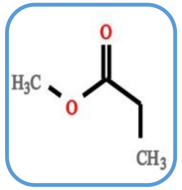 | None | None | None | None | 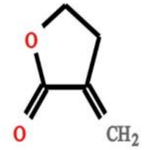 | MESSAR |

| | | | | | | | | | |
|---|---|---|---|---|---|---|---|---|---|
| 76 | None | None | None | 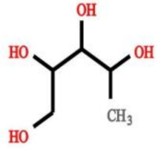 | 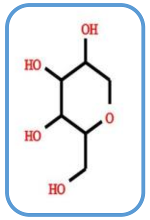 | 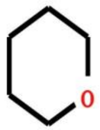 | [Pentose ($C_5$-sugar)-$H_2O$] related loss â€“ indicative for conjugated pentose sugar - EF fits | 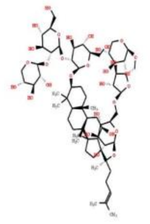 | CSI:FingerID &MS2LDA |
| 77 | 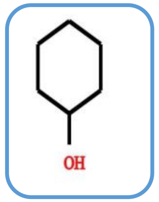 | 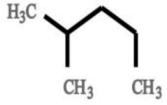 | 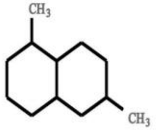 | 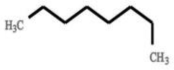 | 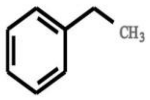 | 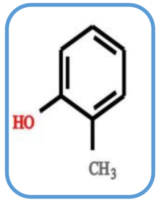 | Fragments indicative for tyrosine related substructure (MzCloud) | 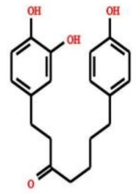 | MESSAR &CSI:FingerID |
| 78 | 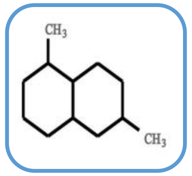 | 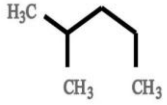 | 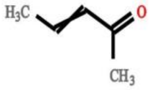 | 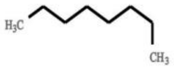 | 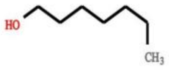 | 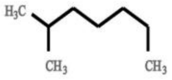 | Sterone related | 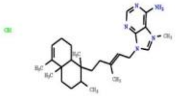 | MESSAR &MS2LDA |
| 79 | 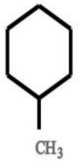 | 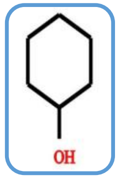 | 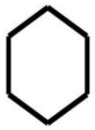 | None | None | None | None | 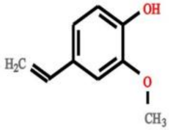 | MESSAR |
| 80 | 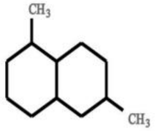 | 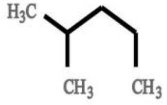 | 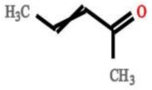 | 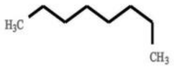 | 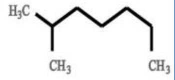 | 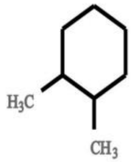 | Steroid core related ($C_{18}H_{21}$ and smaller fragments thereof - with $C_{12}H_{13}$, $C_{11}H_{11}$, and $C_{11}H_{13}$ most probable) | 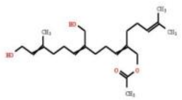 | CSI:FingerID |

| | | | | | | | | | |
|---|---|---|---|---|---|---|---|---|---|
| 81 | 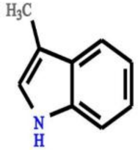 | 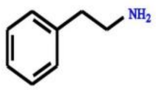 | 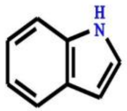 | 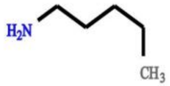 | 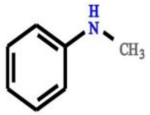 | 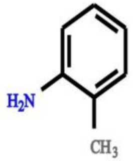 | None | 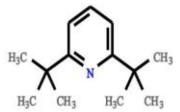 | None |
| 82 | 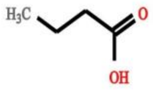 | 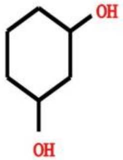 | 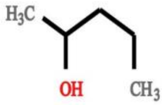 | 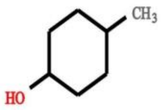 | 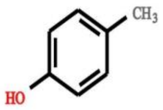 | 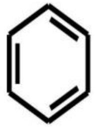 | None | 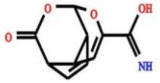 | None |
| 83 | 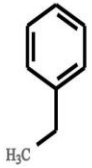 | 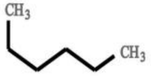 | 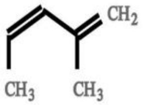 | 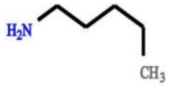 | 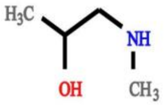 | 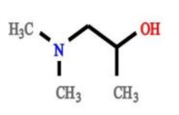 | Loss of CH2O2 - indicative for underivatized carboxilic acid group | 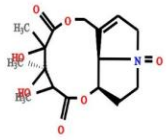 | CSI:FingerID |
| 84 | 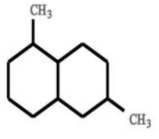 | 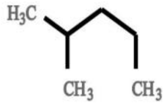 | 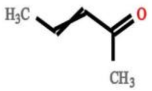 | 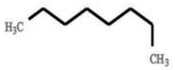 | 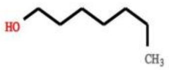 | 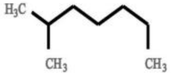 | Sterone related | 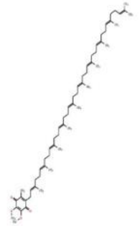 | MESSAR & MS2LDA |
| 85 | 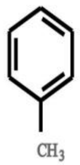 | 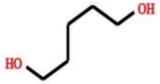 | 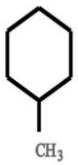 | None | None | None | Fragments indicative for ethylphenol substructure (i.e. resulting from Tyramine â€“ MzCloud) | 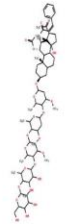 | None |

| | | | | | | | | | |
|---|---|---|---|---|---|---|---|---|---|
| 86 | 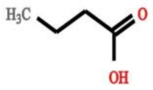 | 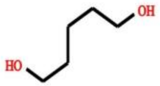 | 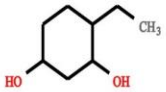 | 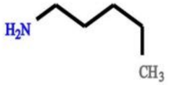 | 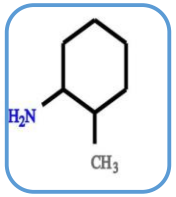 | 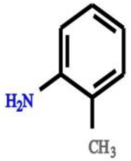 | None | 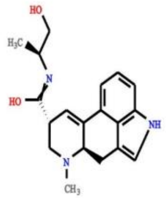 | CSI:FingerID |
| 87 | 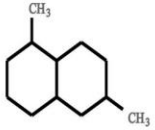 | 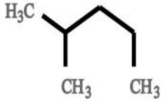 | 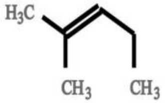 | 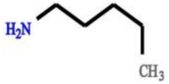 | None | None | Water loss - indicative of a free hydroxyl group â€“ (in beer often seen in sugary structures) | 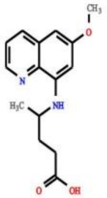 | None |
| 88 | 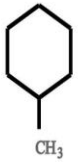 | 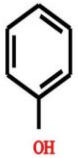 | 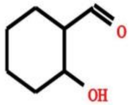 | 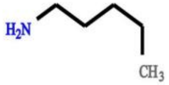 | 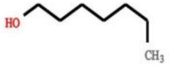 | None | None | 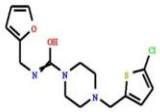 | None |
| 89 | 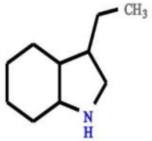 | 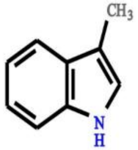 | 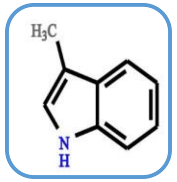 | 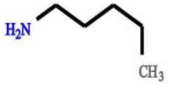 | 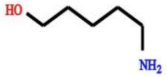 | 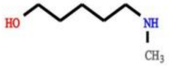 | etrahydro-1H-imidazo[1',5 substructure | 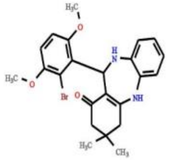 | MESSAR |
| 90 | None | None | None | None | None | None | None | 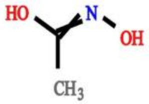 | None |

| | | | | | | | | | |
|---|---|---|---|---|---|---|---|---|---|
| 91 | 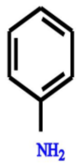 | 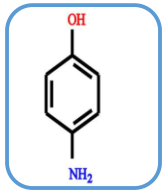 | 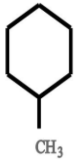 | 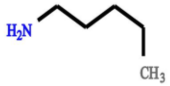 | 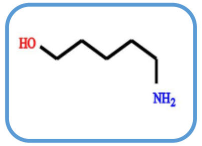 | 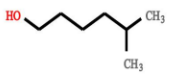 | Loss of CH2O2 - indicative for underivatized carboxilic acid group | 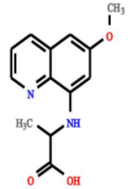 | MESSAR & CSI:FingerID |
| 92 | 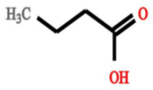 | 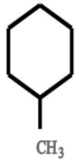 | 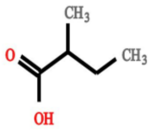 | 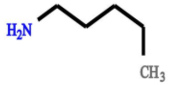 | 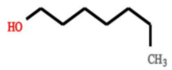 | None | Isopropyl/propylamine substructure (loss based) or isopropyl/propyl side chain | 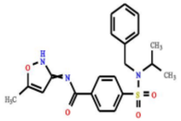 | None |
| 93 | 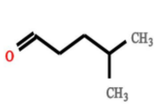 | 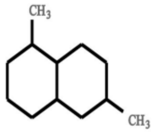 | 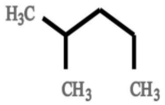 | 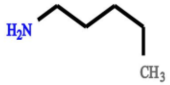 | 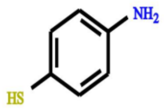 | 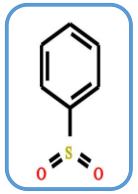 | Benzenesulfonylamide related fragments and losses [ClassyFire - Relevant terms - Substituents: minobenzenesulfonamid Benzenesulfonyl group - Taxa: minobenzenesulfonamide Benzenesulfonamides] | 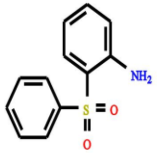 | CSI:FingerID &MS2LDA |
| 94 | None | None | None | 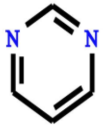 | 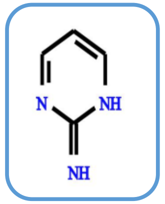 | 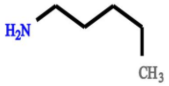 | None | 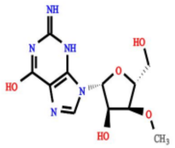 | CSI:FingerID |
| 95 | 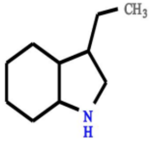 | 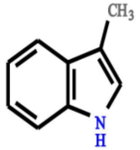 | 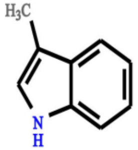 | 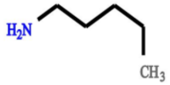 | 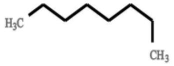 | 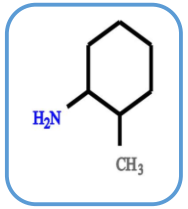 | None | 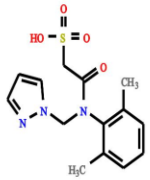 | CSI:FingerID |

| | | | | | | | | | |
|---|---|---|---|---|---|---|---|---|---|
| 96 | 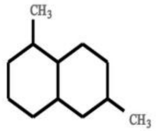 | 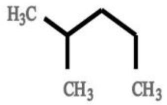 | 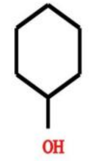 | 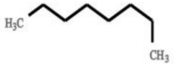 | 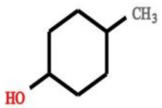 | 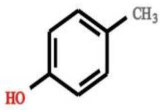 | ethyl-4-oxo-3,4-dihydro-2I substructure | 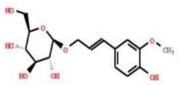 | MESSAR & CSI:FingerID |
| 97 | None | None | None | None | None | None | None | 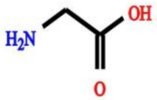 | None |
| 98 | 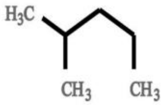 | 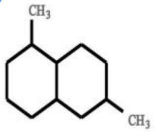 | 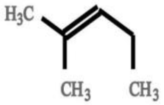 | 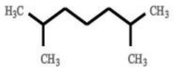 | 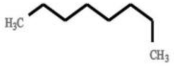 | 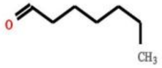 | Water loss - indicative of a free hydroxyl group â€“ (in beer often seen in sugary structures) | 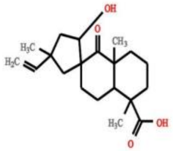 | MESSAR |
| 99 | None | None | None | 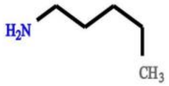 | 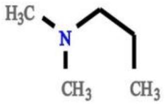 | 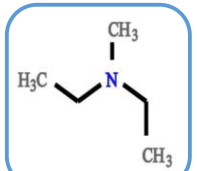 | None | 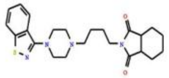 | CSI:FingerID |
| 100 | None | None | None | None | None | None | None | 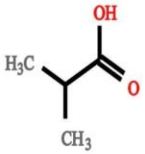 | None |

| | | | | | | | | | |
|---|---|---|---|---|---|---|---|---|---|
| 101 | None | None | None | None | None | None | Amine loss - Indicative for free NH2 group in fragmented molecule | 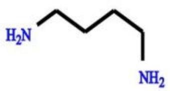 | MS2LDA |
| 102 | 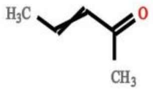 | 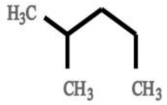 | 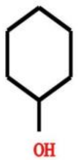 | 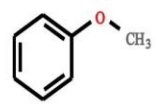 | 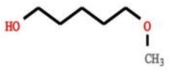 | 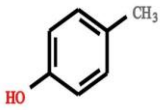 | None | 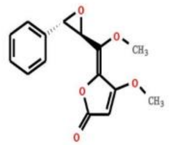 | CSI:FingerID |
| 103 | None | None | None | None | None | None | None | 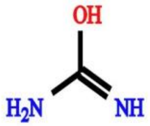 | None |
| 104 | 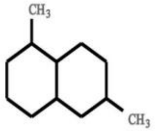 | 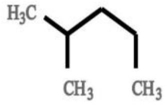 | 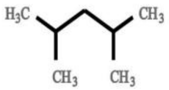 | 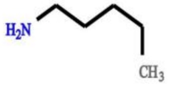 | 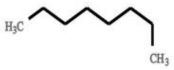 | 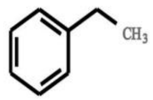 | Fragments indicative for ferulic acid based substructure (MzCloud) | 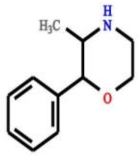 | CSI:FingerID |
| 105 | 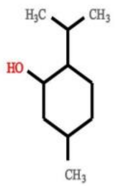 | 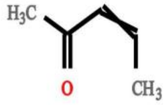 | 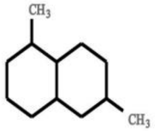 | 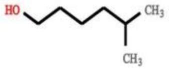 | 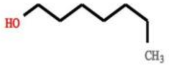 | 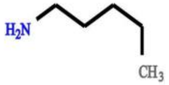 | xy)phenyl)-2-oxo-2H-chrom related substructure (or isomeric variants) | 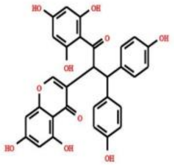 | MESSAR & MS2LDA |

| | | | | | | | | | |
|---|---|---|---|---|---|---|---|---|---|
| 106 | None | None | None | None | None | None | None | 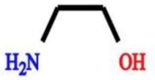 | None |
| 107 | 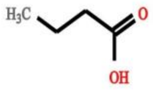 | 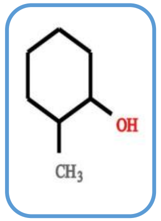 | 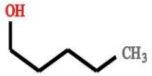 | None | None | None | None | 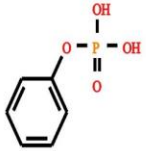 | MESSAR |
| 108 | 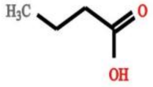 | 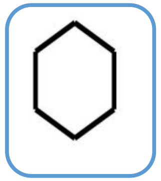 | 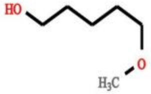 | 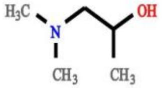 | 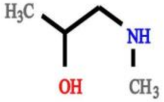 | 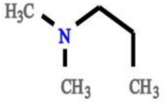 | Sterone steroid related Mass2Motif | 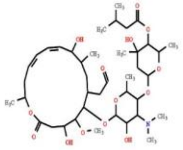 | MESSAR |
| 109 | 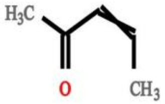 | 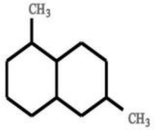 | 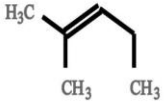 | 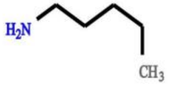 | 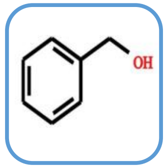 | None | None | 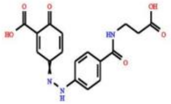 | CSI:FingerID |
| 110 | 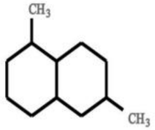 | 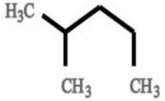 | 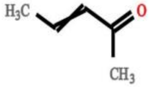 | 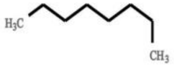 | 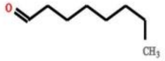 | 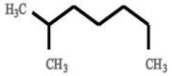 | None | 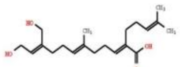 | None |

| | | | | | | | | | |
|---|---|---|---|---|---|---|---|---|---|
| 111 | 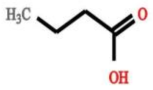 | 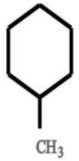 | 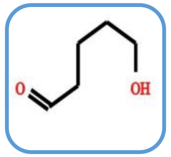 | 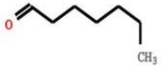 | 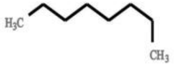 | 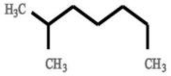 | None | 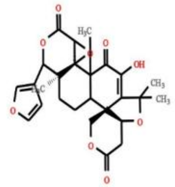 | MESSAR |
| 112 | 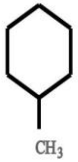 | 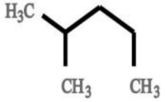 | 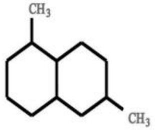 | 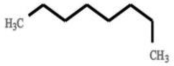 | 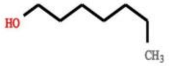 | 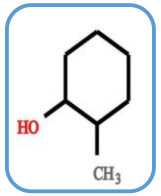 | None | 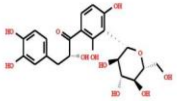 | CSI:FingerID |
| 113 | 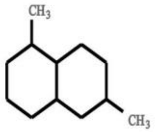 | None | None | 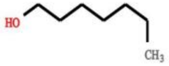 | 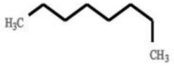 | 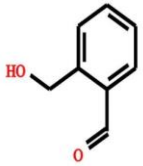 | phthalate substructure | 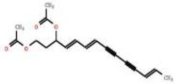 | None |
| 114 | None | None | None | 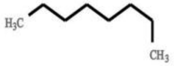 | 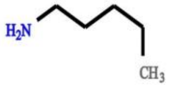 | 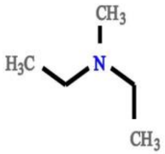 | None | 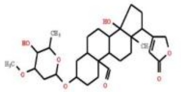 | None |
| 115 | 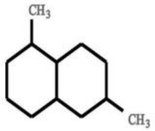 | 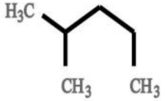 | 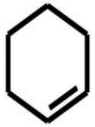 | 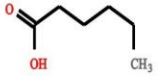 | 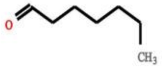 | 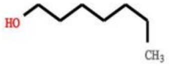 | Double water loss i.e. 2*H20 â€“ Generic feature for metabolites containing several free OH groups attached to a aliphatic chain like sugars. | 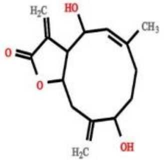 | MS2LDA |

| | | | | | | | | | |
|---|---|---|---|---|---|---|---|---|---|
| 116 | 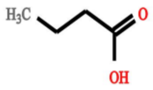 | 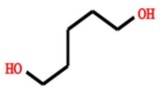 | 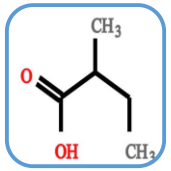 | 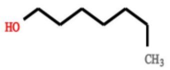 | None | None | phthalate substructure | 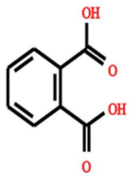 | MESSAR & MS2LDA |
| 117 | 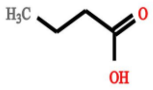 | 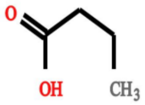 | 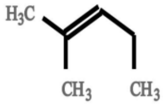 | 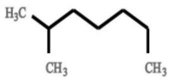 | None | None | None | 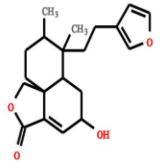 | None |
| 118 | 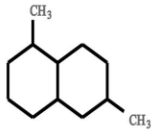 | 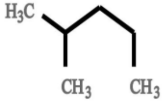 | 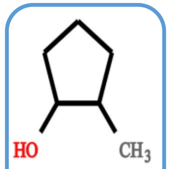 | 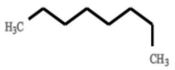 | 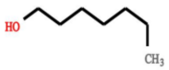 | 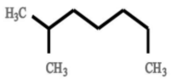 | Double water loss i.e. 2*H20 â€“ Generic feature for metabolites containing several free OH groups attached to a aliphatic chain like sugars. | 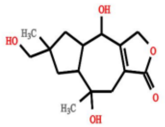 | MESSAR |
| 119 | 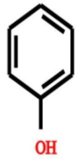 | 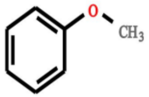 | 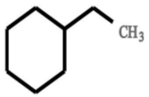 | 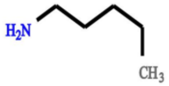 | 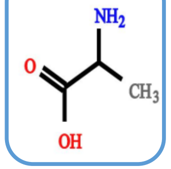 | 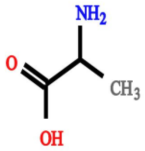 | Fragment indicative for aromatic compounds related to methylbenzene substructure (C7H7 fragment) | 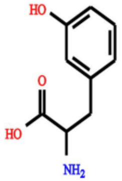 | MESSAR &CSI:FingerID |
| 120 | 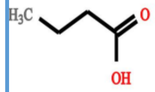 | 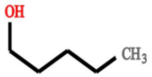 | 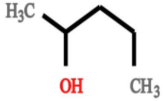 | 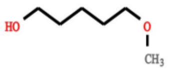 | 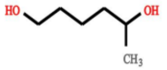 | None | Fragments indicative for cinnamic/hydroxycinnamic acid substructure | 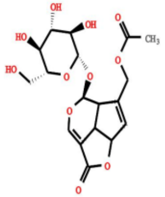 | MESSAR |

| | | | | | | | | | |
|---|---|---|---|---|---|---|---|---|---|
| 121 | 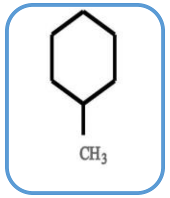 | 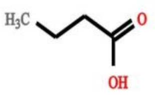 | 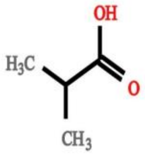 | 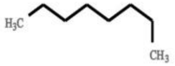 | 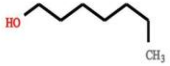 | 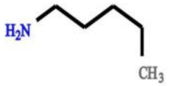 | Loss of CH2O2 - indicative for underivatized carboxilic acid group | 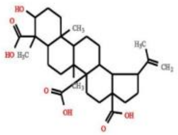 | MESSAR |
| 122 | 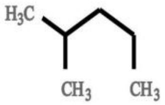 | 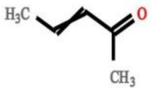 | 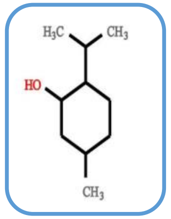 | 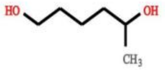 | 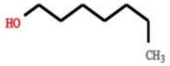 | 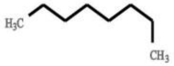 | None | 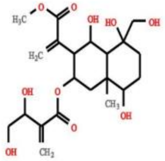 | MESSAR |
| 123 | None | None | None | 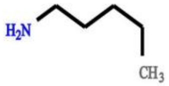 | 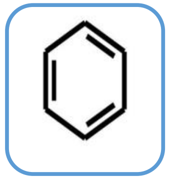 | 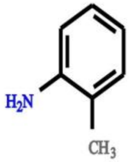 | None | 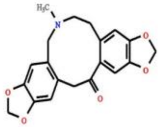 | CSI:FingerID |
| 124 | 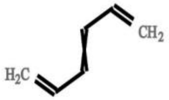 | 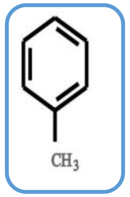 | 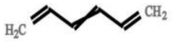 | 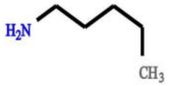 | 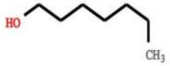 | 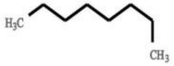 | Double water loss i.e. 2*H20 â€“ Generic feature for metabolites containing several free OH groups attached to a aliphatic chain like sugars. | 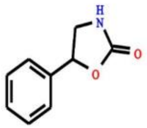 | MESSAR |
| 125 | 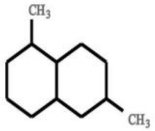 | 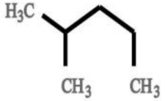 | 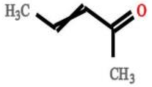 | 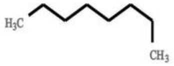 | 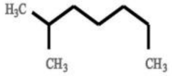 | 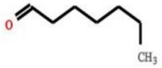 | None | 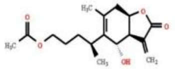 | None |

| | | | | | | | | | |
|---|---|---|---|---|---|---|---|---|---|
| 126 | 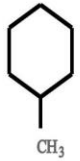 | 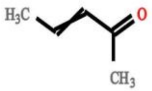 | 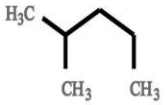 | 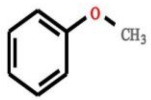 | 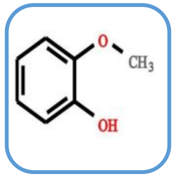 | 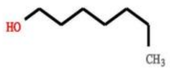 | 2-oxochromen-7-yl [mainly dimethylated] related substructure | 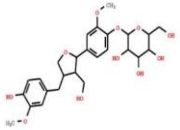 | CSI:FingerID |
| 127 | None | None | None | 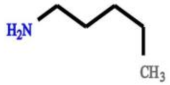 | 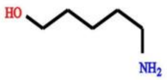 | 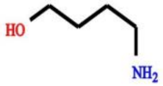 | None | 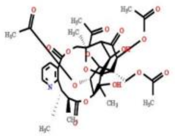 | None |
| 128 | 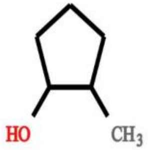 | 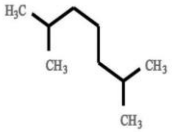 | 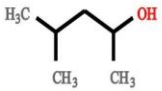 | 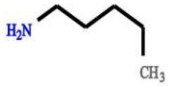 | 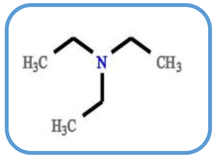 | 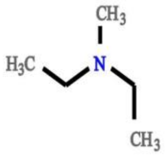 | None | 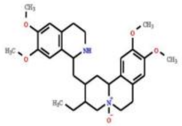 | CSI:FingerID |
| 129 | None | None | None | None | None | None | None | 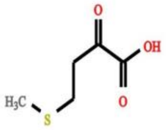 | None |
| 130 | None | None | None | None | None | None | None | 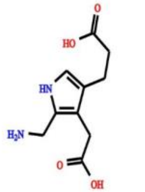 | None |

| | | | | | | | | | |
|---|---|---|---|---|---|---|---|---|---|
| 131 | 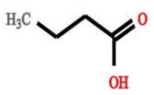 | 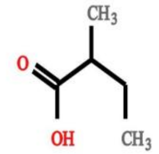 | 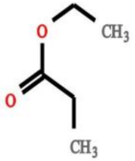 | 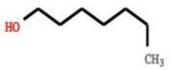 | 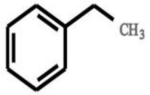 | 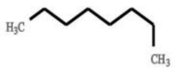 | None | 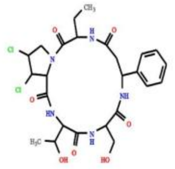 | CSI:FingerID |
| 132 | 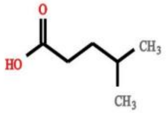 | 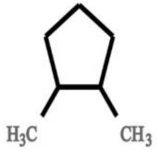 | 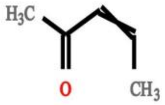 | 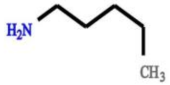 | None | None | None | 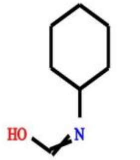 | None |
| 133 | 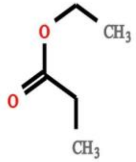 | 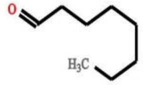 | 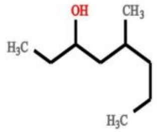 | None | None | None | Amine loss - Indicative for free NH2 group in fragmented molecule | 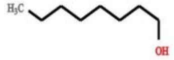 | None |
| 134 | 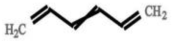 | 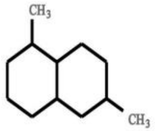 | 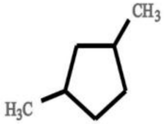 | 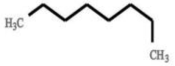 | 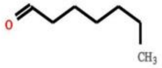 | 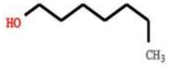 | None | 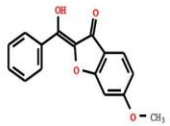 | None |
| 135 | None | None | None | None | None | None | Fragments indicative of a glycosylation â€“ i.e. indicative for a sugar conjugation (in beer often related to glucose) | 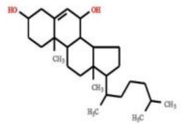 | None |

| | | | | | | | | | |
|---|---|---|---|---|---|---|---|---|---|
| 136 | None | None | None | 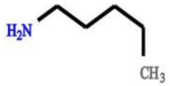 | None | None | None | 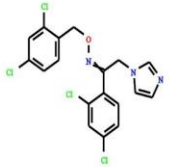 | None |
| 137 | 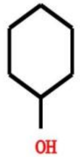 | 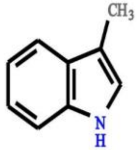 | 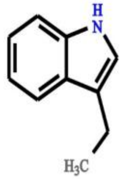 | 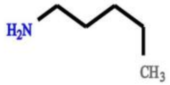 | 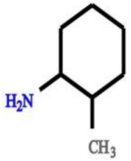 | 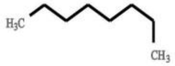 | None | 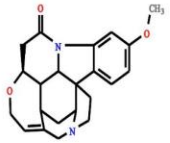 | MESSAR |
| 138 | 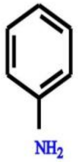 | 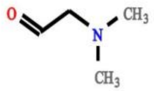 | 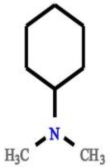 | 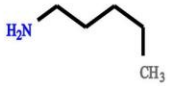 | 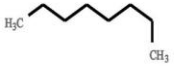 | 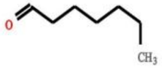 | Small nitrogen containing fragment ion â€“ often proline or ornithine derived â€“ most abundant fragment in beer data | 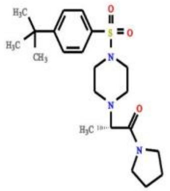 | MESSAR |
| 139 | None | None | None | None | None | None | None | 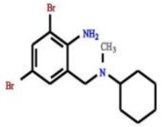 | None |
| 140 | 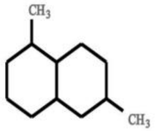 | 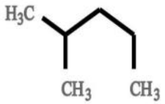 | 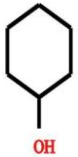 | 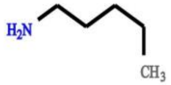 | 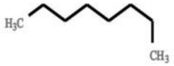 | 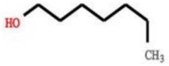 | amino)methyl]cyclohexane acid related Mass2Motif (losses) | 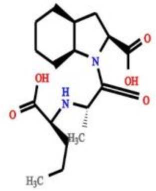 | MESSAR |

| | | | | | | | | | |
|---|---|---|---|---|---|---|---|---|---|
| 141 | 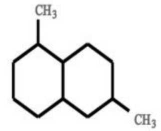 | 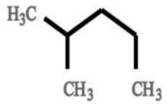 | 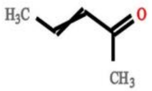 | 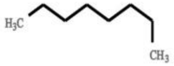 | 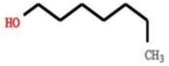 | 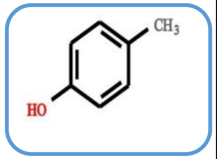 | None | 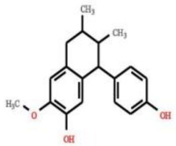 | CSI:FingerID |
| 142 | 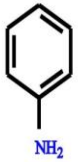 | 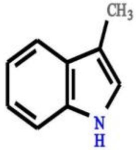 | 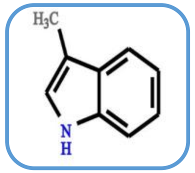 | 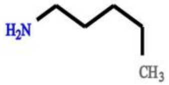 | 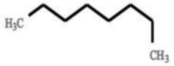 | 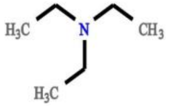 | None | 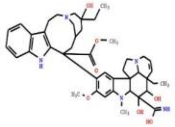 | MESSAR |
| 143 | None | None | None | 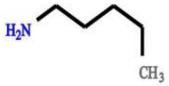 | 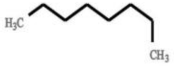 | 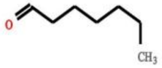 | None | 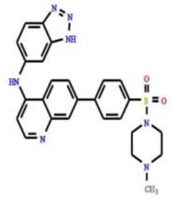 | None |
| 144 | 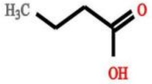 | 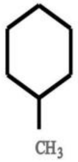 | 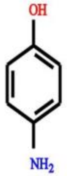 | 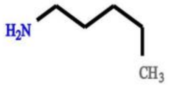 | 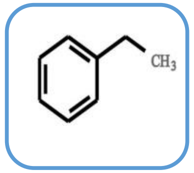 | 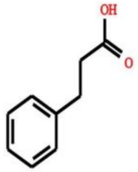 | CO loss - indicative for presence of ketone/aldehyde/lactone group (C=O) | 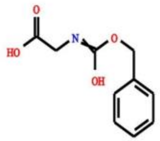 | CSI:FingerID |
| 145 | 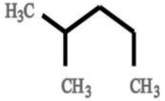 | 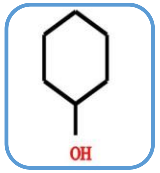 | 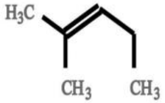 | None | None | None | None | 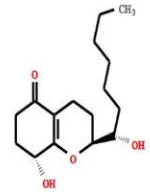 | MESSAR |

| | | | | | | | | | |
|---|---|---|---|---|---|---|---|---|---|
| 146 | 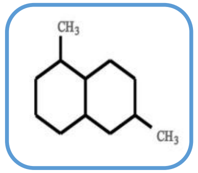 | 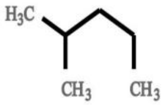 | 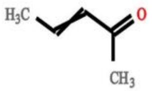 | 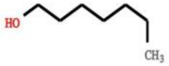 | 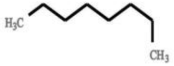 | 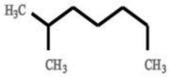 | Hydroxy-Pregnenolone related fragments (not very specific - indicative for presence of steroid backbone) | 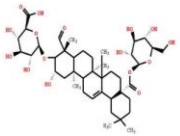 | MESSAR & MS2LDA |
| 147 | 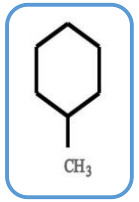 | 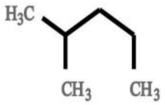 | 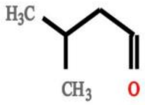 | 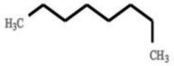 | 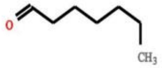 | 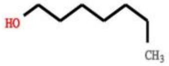 | None | 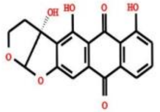 | MESSAR |
| 148 | 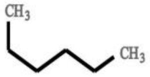 | 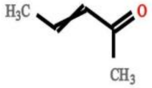 | 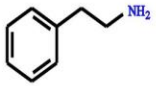 | 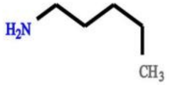 | 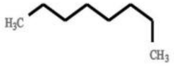 | 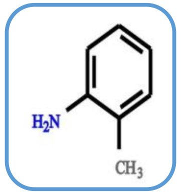 | Indole substructure | 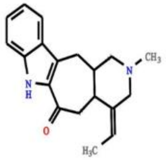 | CSI:FingerID & MS2LDA |
| 149 | None | None | None | None | None | None | Loss possibly indicative of carboxylic acid group with 1-carbon attached. | 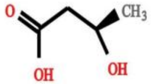 | MS2LDA |
| 150 | None | None | None | 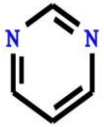 | None | None | None | 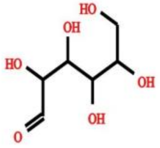 | None |

| | | | | | | | | | |
|---|---|---|---|---|---|---|---|---|---|
| 151 | 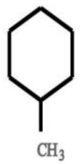 | 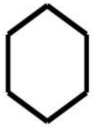 | 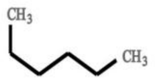 | 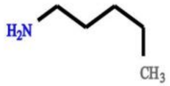 | 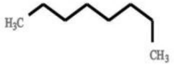 | 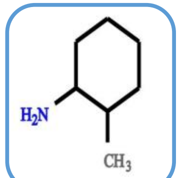 | tetrahydroisoquinoline substructure | 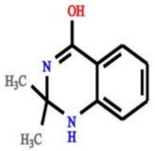 | CSI:FingerID & MS2LDA |
| 152 | 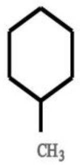 | 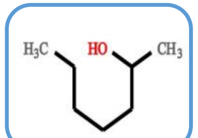 | 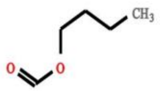 | 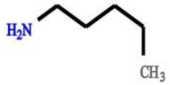 | 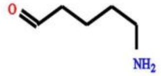 | None | Fragment ions indicative for C6H12NO substructure (in beer related to N-acetylputrescine - MzCloud) | 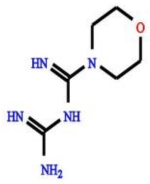 | MESSAR & MS2LDA |
| 153 | 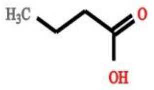 | 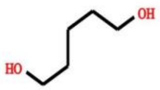 | 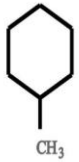 | 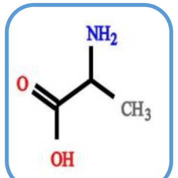 | 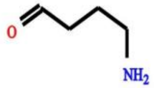 | None | Amine loss - Indicative for free NH2 group in fragmented molecule | 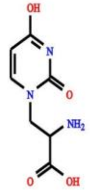 | CSI:FingerID & MS2LDA |
| 154 | 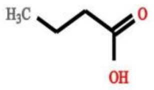 | 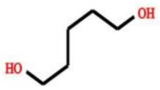 | 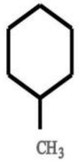 | 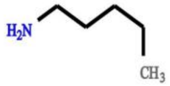 | 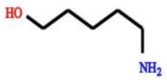 | 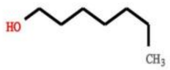 | None | 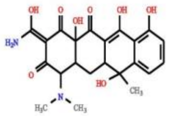 | None |
| 155 | 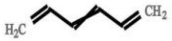 | 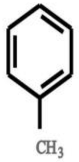 | 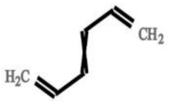 | 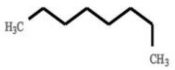 | None | None | None | 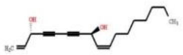 | CSI:FingerID |

| | | | | | | | | | |
|---|---|---|---|---|---|---|---|---|---|
| 156 | None | None | None | 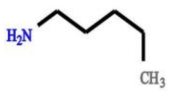 | None | None | None | 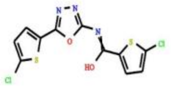 | None |
| 157 | 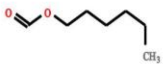 | 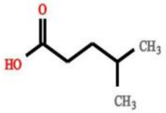 | 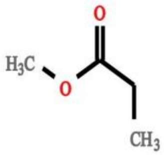 | 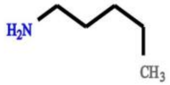 | 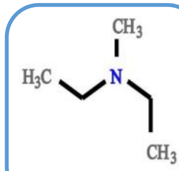 | 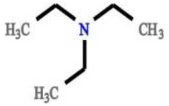 | None | 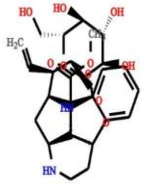 | MESSAR & CSI:FingerID |
| 158 | 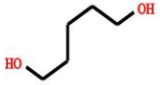 | 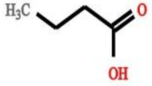 | 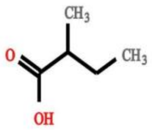 | 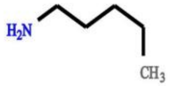 | 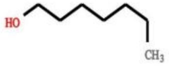 | 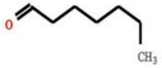 | None | 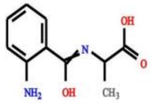 | MESSAR & CSI:FingerID |
| 159 | None | None | None | None | None | None | None | 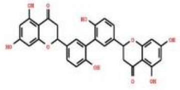 | None |
| 160 | 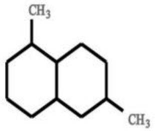 | 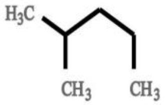 | 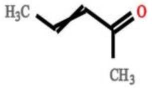 | 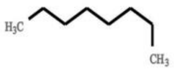 | 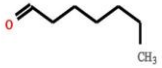 | 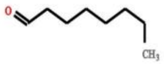 | Hydroxy-Pregnenolone related fragments (not very specific - indicative for presence of steroid backbone) | 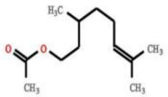 | MESSAR |

| | | | | | | | | | |
|---|---|---|---|---|---|---|---|---|---|
| 161 | 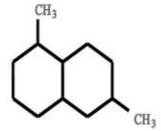 | 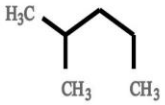 | 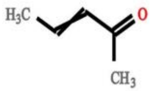 | 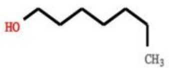 | 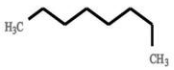 | 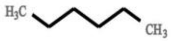 | None | 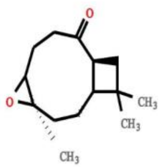 | None |
| 162 | 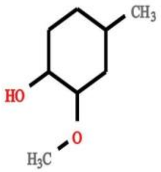 | 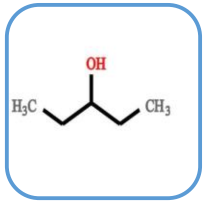 | 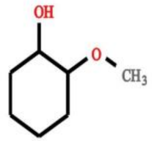 | 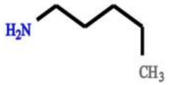 | 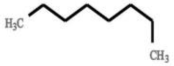 | None | None | 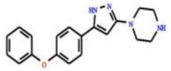 | MESSAR |
| 163 | 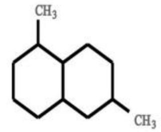 | 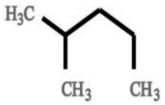 | 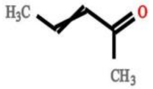 | 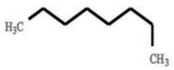 | 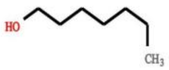 | 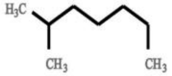 | Pregn-4 or 5-ene-3-dione substructure | 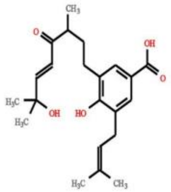 | None |
| 164 | 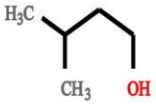 | 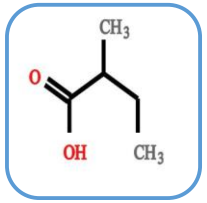 | 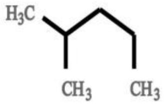 | 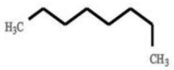 | 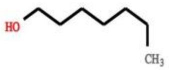 | 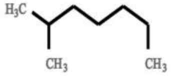 | None | 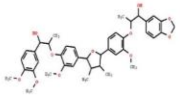 | MESSAR |
| 165 | 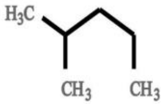 | 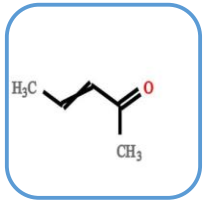 | 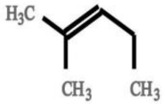 | 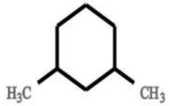 | 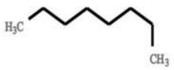 | 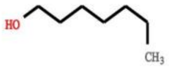 | None | 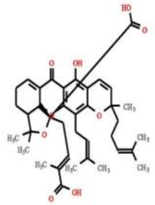 | MESSAR |

| | | | | | | | | | |
|---|---|---|---|---|---|---|---|---|---|
| 166 | 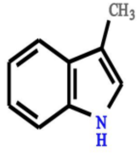 | 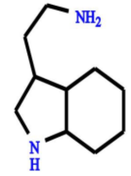 | 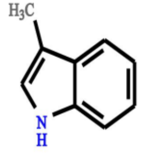 | 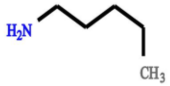 | 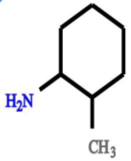 | 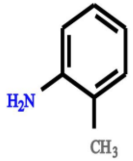 | None | 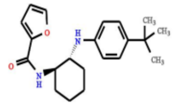 | CSI:FingerID |
| 167 | 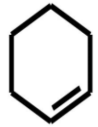 | 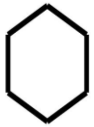 | 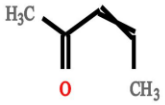 | 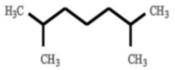 | 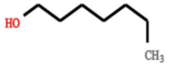 | 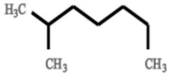 | None | 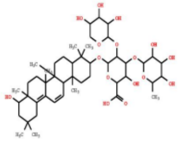 | MESSAR |
| 168 | 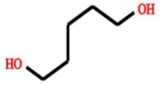 | 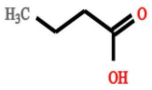 | 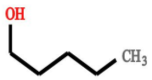 | 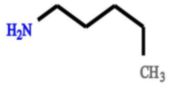 | 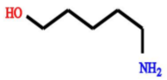 | 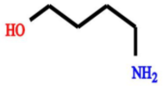 | None | 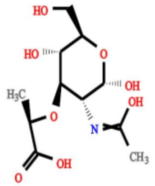 | MESSAR & CSI:FingerID |
| 169 | 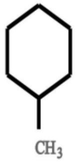 | 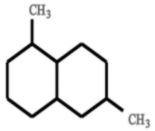 | 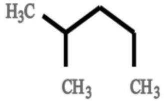 | 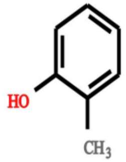 | 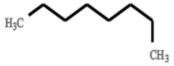 | 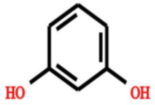 | Loss of [hexose-H2O] â€“ indication of hexose conjugation (for example glucose) [ClassyFire - Relevant terms - Substituents: Hexose monosaccharide [illegible] | 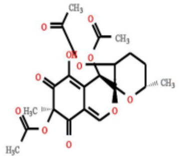 | CSI:FingerID & MS2LDA |
| 170 | None | None | None | 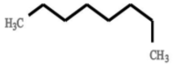 | 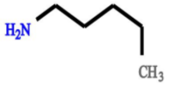 | 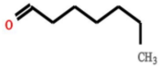 | None | 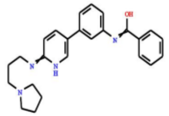 | None |

| | | | | | | | | |
|---|---|---|---|---|---|---|---|---|
| 171 | 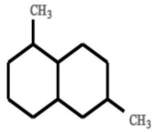 | 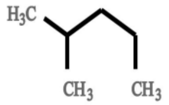 | 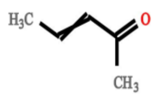 | 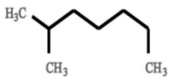 | 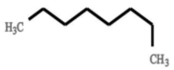 | 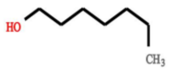 | Steroid core related (C18H21 and smaller fragments thereof - with C12H13, C11H11, and C11H13 most probable) | 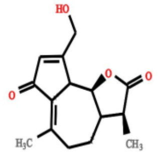 | MESSAR |
| 172 | 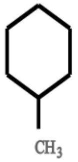 | 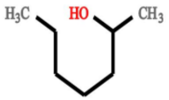 | 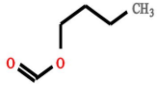 | 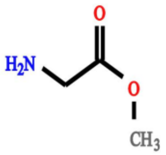 | 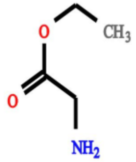 | None | Fragments indicative of a glycosylation â€“ i.e. indicative for a sugar conjugation (in beer often related to glucose) | 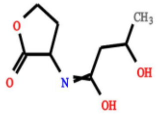 | CSI:FingerID |
| 173 | None | None | None | 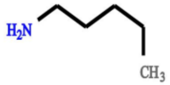 | 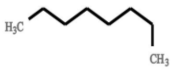 | 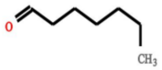 | None | 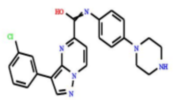 | None |
| 174 | 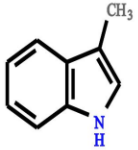 | 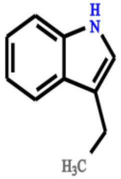 | 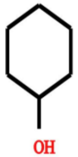 | 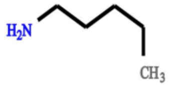 | 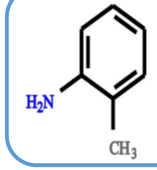 | 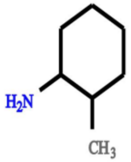 | Indole substructure | 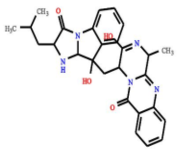 | CSI:FingerID |
| 175 | 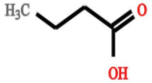 | 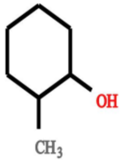 | 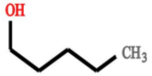 | 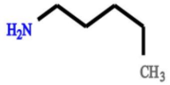 | None | None | xyphenyl)-2-oxo-2H-chrom relared substructure (or isomeric variants) | 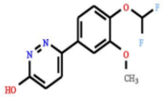 | MESSAR & MS2LDA |

| | | | | | | | | | |
|---|---|---|---|---|---|---|---|---|---|
| 176 | 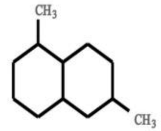 | 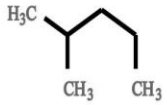 | 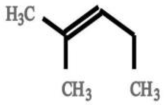 | 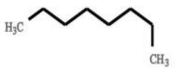 | 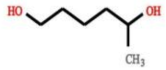 | 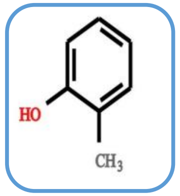 | Fragments indicative for cinnamic/hydroxycinnamic acid substructure | 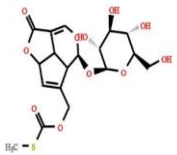 | CSI:FingerID |
| 177 | 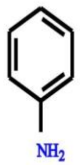 | 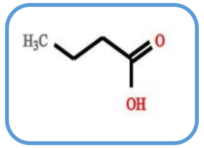 | 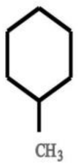 | 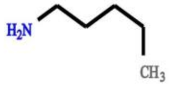 | 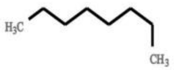 | 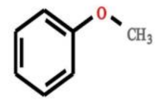 | Indole substructure | 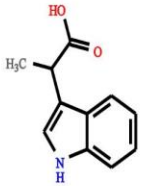 | MESSAR & MS2LDA |
| 178 | 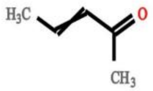 | 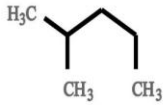 | 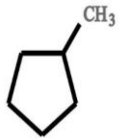 | 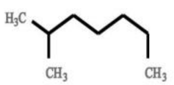 | 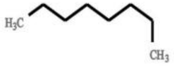 | 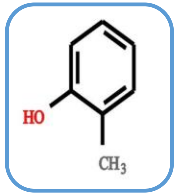 | CO loss - indicative for presence of ketone/aldehyde/lactone group (C=O) | 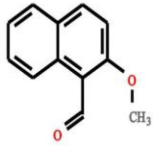 | CSI:FingerID |
| 179 | 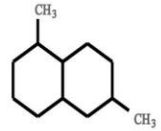 | 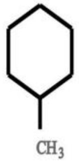 | 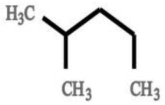 | 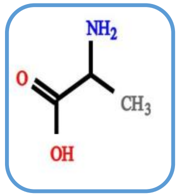 | 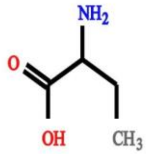 | 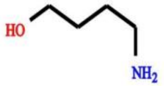 | None | 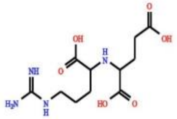 | CSI:FingerID |
| 180 | None | None | None | 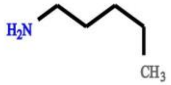 | None | None | None | 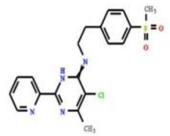 | None |

| | | | | | | | | | |
|---|---|---|---|---|---|---|---|---|---|
| 181 | None | None | None | 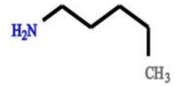 | None | None | None | 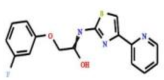 | None |
| 182 | 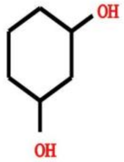 | 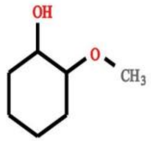 | 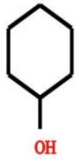 | 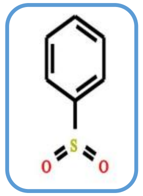 | 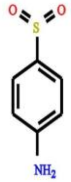 | 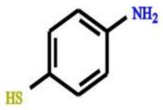 | None | 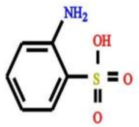 | CSI:FingerID |
| 183 | None | None | None | 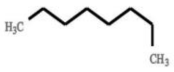 | 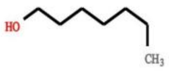 | 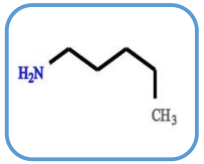 | None | 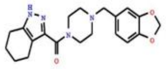 | CSI:FingerID |
| 184 | None | None | None | 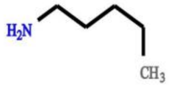 | 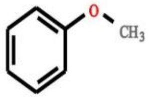 | 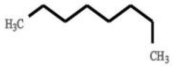 | None | 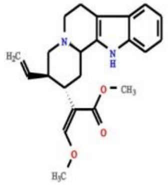 | None |
| 185 | 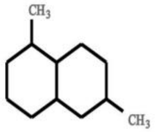 | 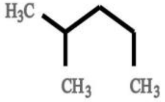 | 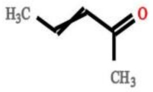 | 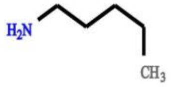 | 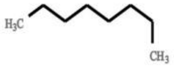 | 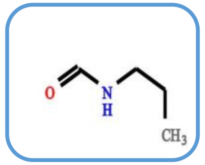 | Sterone related | 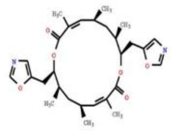 | CSI:FingerID |
