## Supplementary material for "MESSAR: Automated recommendation of metabolite substructures from tandem mass spectra": S2 Table

**Table S2** Examples of unexplained positive ion mode GNPS M2Ms annotated by MESSAR rules

| **M2M** | **Matched MESSAR Target Rules** | | | |
| --- | --- | --- | --- | --- |
| ID | Feature Type | Rule Feature(s) | Substructure(s) | Interpretation |
| **140** | Mass | 159.0804 | 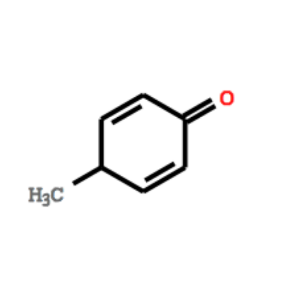 | Aminophenol |
|  | Mass | 145.0648 |  |  |
|  | Mass, Mass | 159.0804, 145.0648 |  |  |
|  | MDiff | 57.0215 | 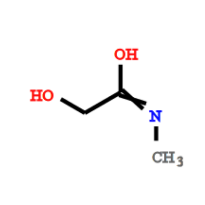 |  |
|  | Mass | 107.0491 | 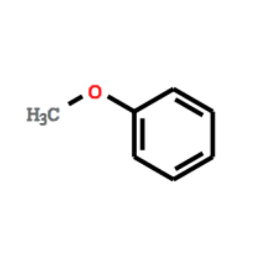 |  |
|  | Mass | 133.0648 | 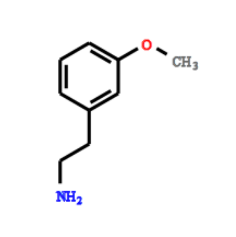 |  |
| **167**  **218**  **264** | Mass | 147.0804 | 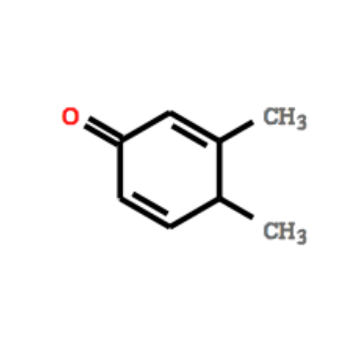 | Dimethyl  -benzene  (ortho-xylene) |
|  | Mass | 135.0804 |  |  |
|  | Mass | 171.0804 |  |  |
|  | Mass | 161.0961 |  |  |
|  | Mass | 185.0961 | 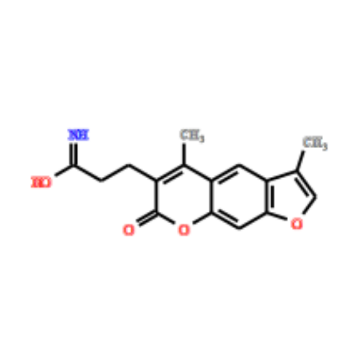 |  |
|  | Mass | 173.0961 | 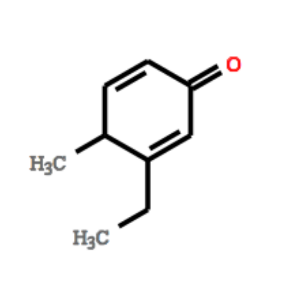 |  |
| **361** | Mass | 117.0573 | 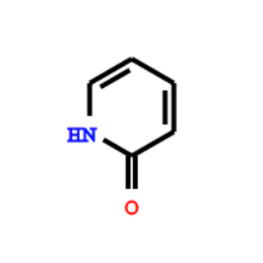 | Hydroxypyridine |
|  | Mass | 134.06 | 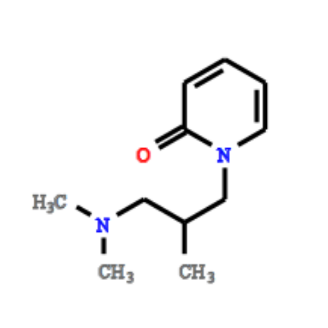 |  |
|  | Mass | 149.0597 | 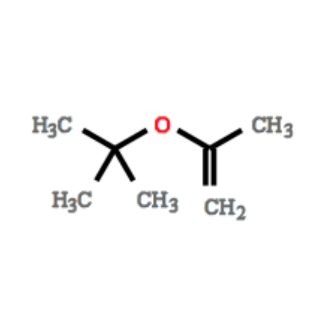 |  |
|  | Mass | 107.0491 | 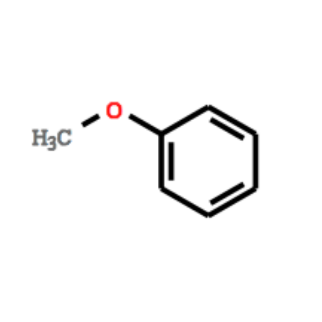 |  |
|  | MDiff | 71.0371 | 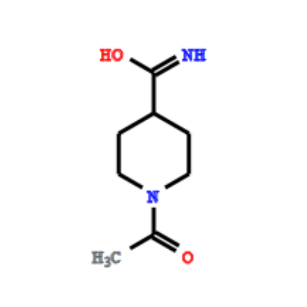 |  |
|  | MDiff | 31.0422 | 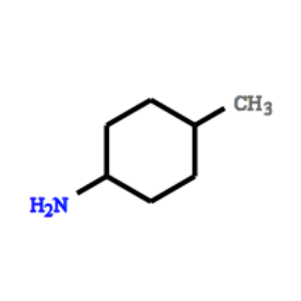 |  |
| **400** | Mass | 157.1012 | 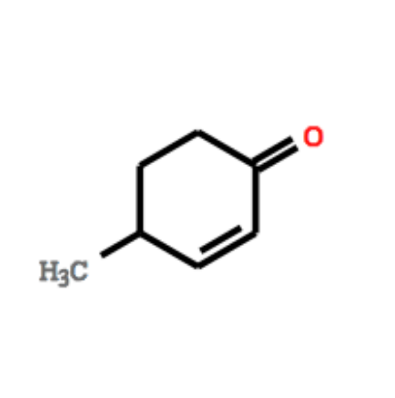 | Coumarin |
|  | Mass | 145.1012 |  |  |
|  | Mass, Mass | 145.1012, 131.0855 |  |  |
|  | Mass | 141.0699 | 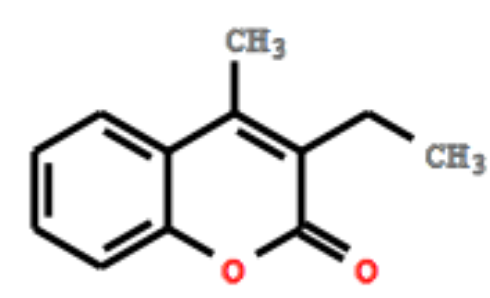 |  |
|  | Mass | 128.062 |  |  |
|  | Mass, Mass | 141.0699, 128.062 |  |  |
