## Supplementary material for "MESSAR: Automated recommendation of metabolite substructures from tandem mass spectra": S1 Text

**S1 Text: Supporting Method**

**Association rule mining**

ARM is a pattern mining technique that detects substructures consistently related to the occurrence or co-occurrence of spectral features. ARM algorithm is usually applied on a list of transactions, with each transaction a set of items. In our approach, each item represents a spectral feature *x_i_* or a molecular substructure *y_j_*. A transaction is then the set of peaks, mass differences and non-redundant substructures, {*x_1_, x_2_, x_3_… y_1_, y_2,_ y_3…_*}. Overall, we have a list of 3146 transactions from either target or decoy spectral library. We use PyFIM package in Python for association rule mining [1].

Patterns are mined separately for target and decoy libraries. A pattern is called an itemset and it can contain one or more items e.g. {*x_2_*}, {*x_2_, x_4_, y_8_*}. *Support* is defined as the number of transactions in which that itemset appears. An itemset is considered frequent if its *support* is higher than a user-defined threshold. For example, the frequent itemset X = *{x_2_, x_4_}* with a *support* of 30 means that spectral features *x_2_* and *x_4_* appear together in 30 out of 3146 mass spectra.

Frequent itemset shows interesting patterns but does not express the link between spectral features and substructures. ARM further searches relationships between frequent item sets expressed as association rules: X → Y. X is called the body of the rule and it should only contain spectral features, while Y is called the head of the rule and it must be a single substructure. The *support* of the association rule X → Y is equal to the *support* of the itemset X ∪ Y. The *confidence* is the frequency that Y is present in a transaction given that X is also present in that transaction, and is defined as:

$$Confidence \left( X\to Y \right)=\frac{Support (X \cup Y)}{Support (X)}$$

When no constraints are applied, the ARM algorithm can detect millions even milliards rules. Such complexity goes far beyond what personal computers can handle. In the training phase of MESSAR, we generated as many rules as possible by controlling three parameters of ARM: the maximum size of itemset X, the minimal *support* of the rule and minimal *confidence*. With a 2.80 GHz Intel Core-i7 CPU and 16 GBytes of RAM under the Windows 10 operating system, we applied the lowest possible parameters to mine the spectral library: the *support* threshold at 5 and *confidence* at 0.10, and the body of the rule can contain up to three spectral features. We used the same set of parameters to generate target and decoy rules.

A basic filter of *lift* was applied to generated rules. The *lift* measures how much more often X and Y occur together than expected if they were statistically independent:

$$Lift \left( X\to Y \right)=\frac{Support (X \cup Y)}{Support \left( X \right) \times Support \left( Y \right)}$$

According to its definition, only rules with *lift* > 1 are interesting to explore. After applying this filter, we obtained 20747 rules from target library and 15480 rules from decoy library.

*Support*, *confidence* and *lift* are called *interestingness measures* of association rules [2]. To clearly illustrate the objective of MESSAR (i.e. substructure prediction), we introduce rules as individual binary classifiers (Method section). Rather than using *interestingness measure*, a terminology specific for association rules, statistical measures such as *precision*, *sensitivity* and *specificity* are more adapted to evaluate the prediction power of rules. A confusion matrix was created for each MESSAR rule using training and testing data (Table 1). Statistical measures based on confusion matrix and *interestingness* of association rules are closely related:

$$TP= Support \left( X \cup Y \right)$$

$$TP+FP= Support \left( X \right)$$

$$TP+FN= Support \left( Y \right)$$

$$Precision= \frac{TP}{TP+FP}= \frac{Support \left( X \cup Y \right)}{Support \left( X \right)}(= Confidence \left( X \cup Y \right))$$

$$Sensitivity= \frac{TP}{TP+FN}= \frac{Support (X \cup Y)}{Support (Y)}$$

$$Specificity= \frac{TN}{FP+TN}= \frac{Support (\neg X \cup\neg Y)}{Support (\neg Y)}$$

Target and decoy rules were statistically evaluated and compared. Since both target and decoy rules showed high *specificity* (around 0.95 as shown in S1A Fig), we only discussed and compared *precision* and *sensitivity* in the main manuscript text. We chose *sensitivity* for rule filtering, validation and substructure prediction since this metrics better discriminated target and decoy rules (Fig 2C). Moreover, the two rule-generating parameters, *support* and *confidence* (*precision*), had little influence on *sensitivity* – rules with low *support* or *confidence* can still have high *sensitivity* (Fig 2A, S1B Fig). The rule mining strategy by applying lowest possible *support* and *confidence* thresholds have indeed captured meaningful rules.
