## Supplementary material for "MESSAR: Automated recommendation of metabolite substructures from tandem mass spectra": S1 Table

**Table S1** Comparison M2M motifs - MESSAR rules: 26 out of 90 structurally-annotated positive ion mode M2M motifs are explained by MESSAR rules via mass feature matching (at least 5 common features). Examples of substructure recommendations of M2Ms are provided based on the text descriptions of motifs. If several matched rules are associated with the same (set of) feature(s), only the rule with the highest sensitivity is used for validation. When there is a tie for sensitivity, MCS of top rules is reported as the predicted substructure. Here we list all rules whose substructure recommendation agrees with manual annotation of M2Ms according to experts.

| **Mass2Motif** | | **Related MESSAR Target Rules** | | |
| --- | --- | --- | --- | --- |
| ID | Experts’ annotation | Feature Type | Rule Feature(s) | Predicted Substructures |
| **1** | Sterone-related  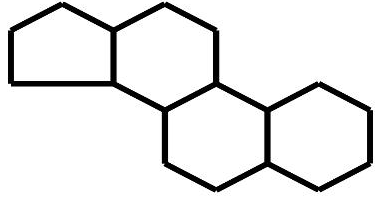 | Mass | 121.0648 | 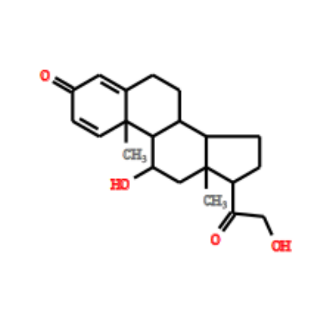 |
|  |  | Mass, Mass | 147.0804, 121.0648 |  |
|  |  | Mass, Mass | 135.0804, 121.0648 | 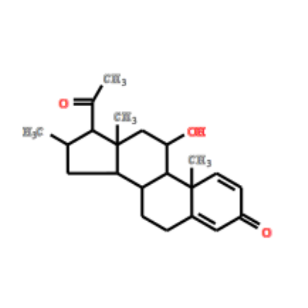 |
|  |  | Mass, Mass | 145.1012, 119.0855 | 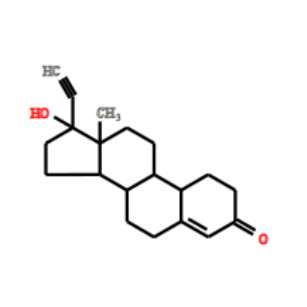 |
| **6** | Diphenyl substructure  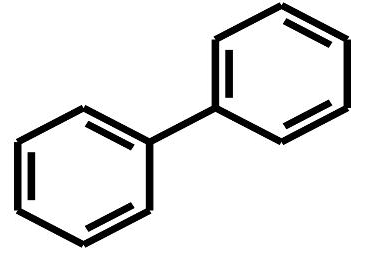 | Mass | 152.062 | 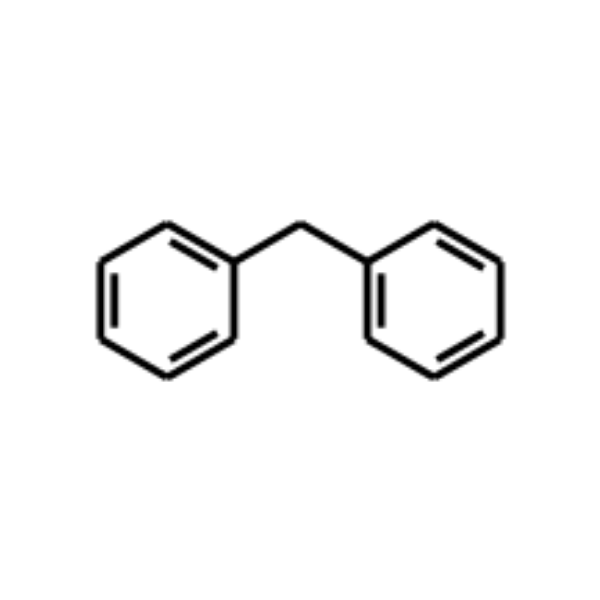 |
|  |  | Mass | 167.0855 |  |
| **21** | Ethyl phenol  (i.e. resulting from Tyramine)  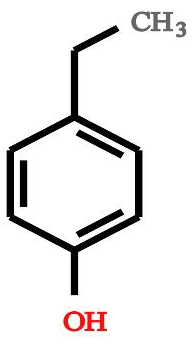 | Mass | 103.0542 | 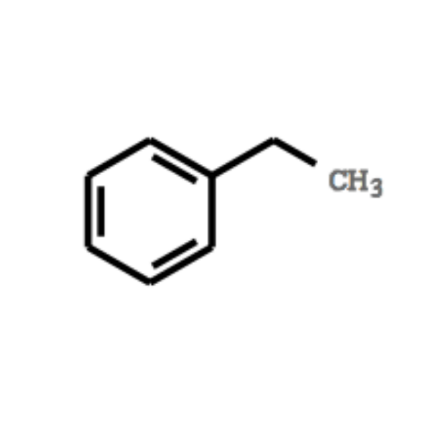 |
|  |  | Mass | 91.0542 |  |
|  |  | Mass, Mass | 121.0648, 103.0542 | 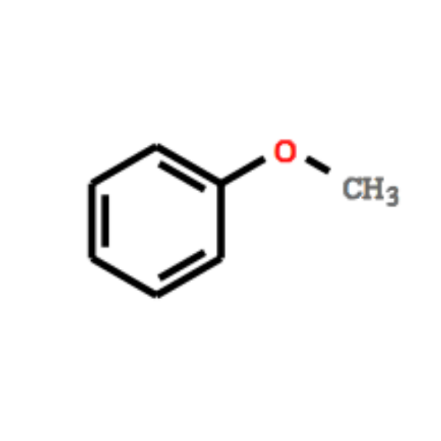 |
| **25** | Indole substructure  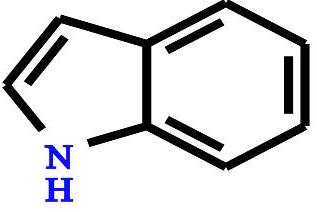 | Mass | 146.06 | 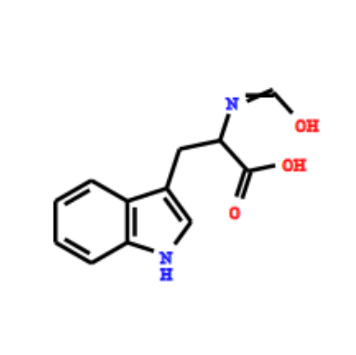 |
|  |  | Mass, Mass | 132.0808, 130.0651 | 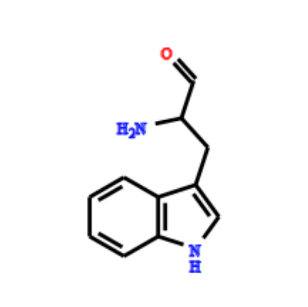 |
|  |  | Mass | 142.0651 |  |
|  |  | Mass, Mass | 144.0808, 130.0651 |  |
|  |  | Mass, Mass | 130.0651, 115.0542 |  |
|  |  | Mass, Mass | 130.0651, 144.0808 | 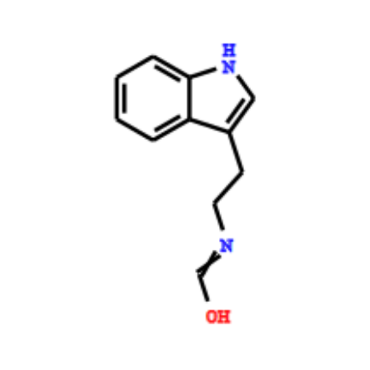 |
|  |  | Mass | 158.06 | 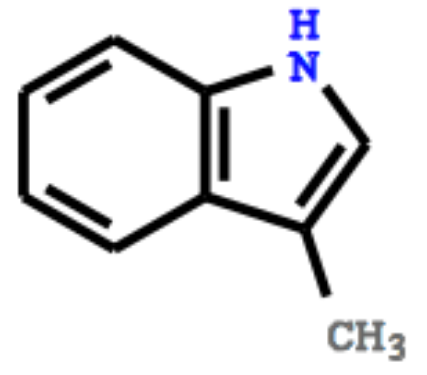 |
| **26** | 6-oxo-7,11-diazatricyclo [7.3.1.0~2,7~] trideca-2,4-diene-11-carboxamide  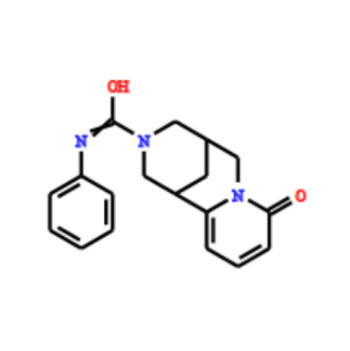  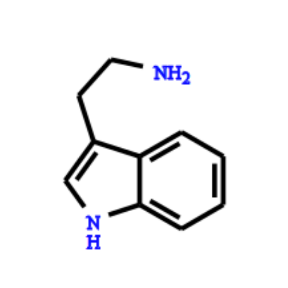 | Mass | 132.0444 | 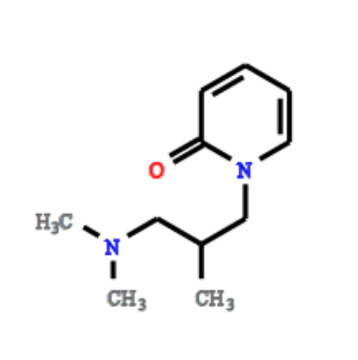 |
|  |  | Mass | 134.06 |  |
|  |  | Mass | 148.0757 |  |
|  |  | Mass | 160.0757 |  |
|  |  | Mass | 172.0757 | 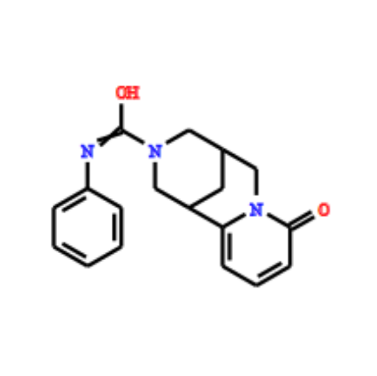 |
|  |  | Mass | 132.0808 | 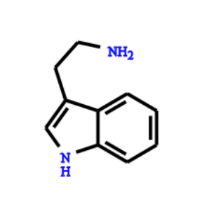 |
| **28** | 2-oxochromen-7-yl [mainly dimethylated] related  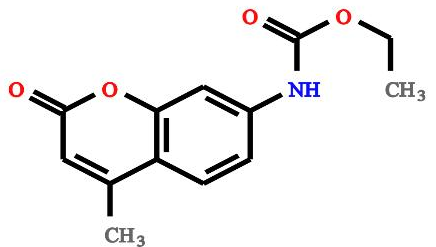 | Mass | 175.0754 | 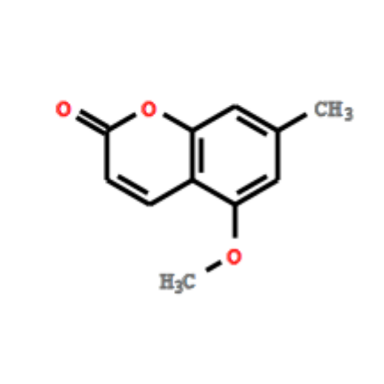 |
|  |  | Mass | 133.0648 | 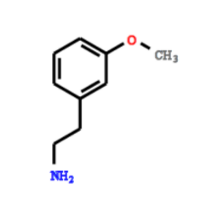 |
| **29** | 1-naphthalenamine  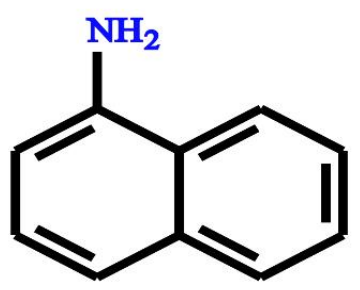 | Mass | 142.0651 | 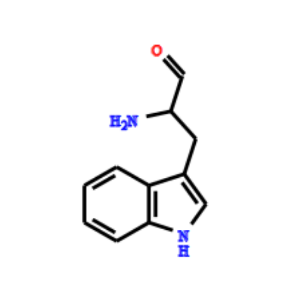 |
|  |  | Mdiff | 78.047 | 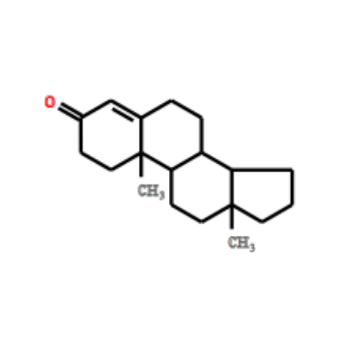 |
| **30** | Methylated [7 or 3,7] 2-(4-Hydroxy-6-oxotetrahydro-2H-pyran-2-yl) ethyl  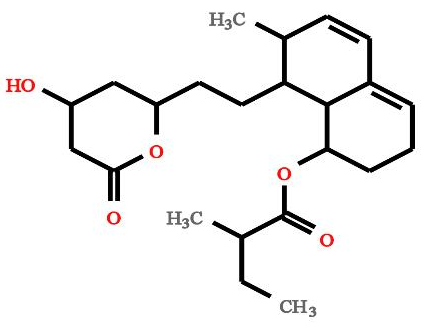 | Mass | 157.1012 | 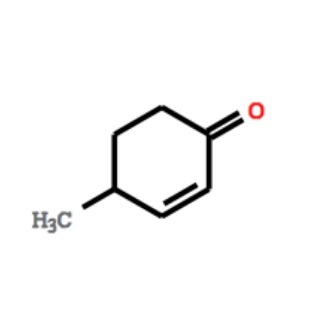 |
|  |  | Mass | 145.1012 |  |
|  |  | Mass, Mass | 145.1012, 131.0855 |  |
|  |  | Mass | 143.0855 | 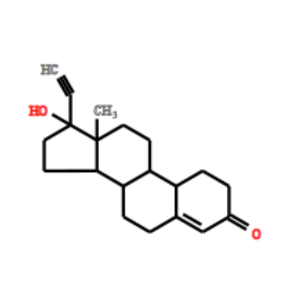 |
|  |  | Mass | 143.0855, 131.0855 |  |
|  |  | Mass, Mass | 155.0855 |  |
| **32** | Pregn-4 or 5-ene-3-dione substructure  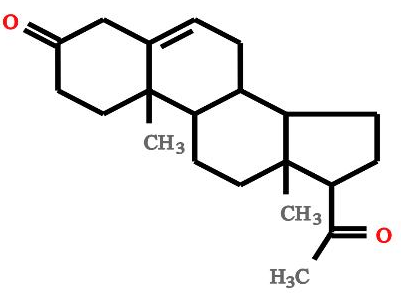 | Mass | 143.0855 | 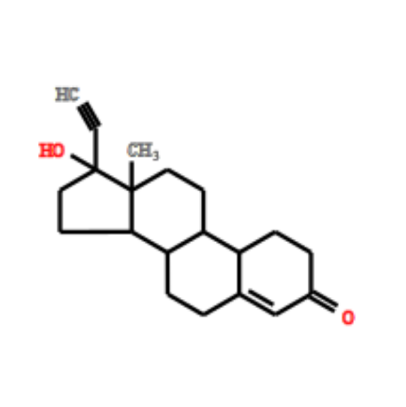 |
|  |  | Mass, Mass | 143.0855, 131.0855 |  |
|  |  | Mass, Mass | 145.1012, 119.0855 |  |
|  |  | Mass | 155.0855 |  |
|  |  | Mass | 131.0855 | 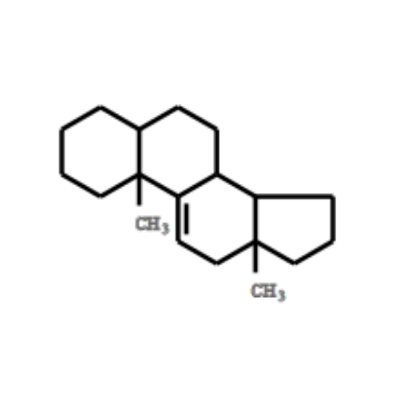 |
|  |  | Mass, Mass | 131.0855, 119.0855 |  |
| **33** | Tetrahydro-isoquinoline substructure  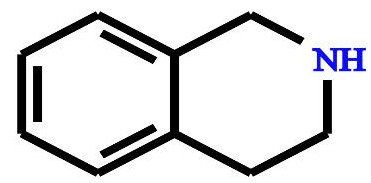 | Mass | 117.0573 | 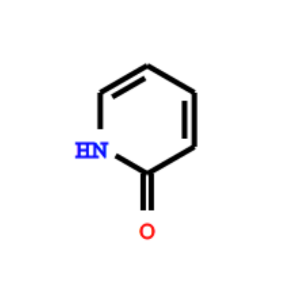 |
|  |  | Mass, Mass | 132.0808, 130.0651 | 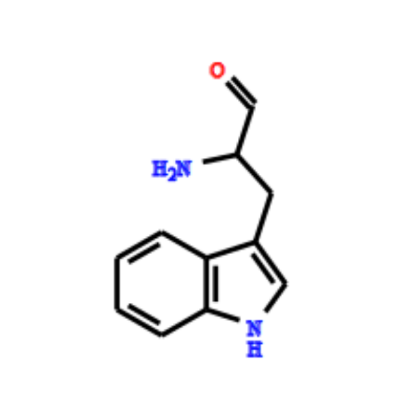 |
|  |  | Mass | 142.0651 |  |
|  |  | Mass, Mass | 144.0808, 130.0651 |  |
|  |  | Mass, Mass | 130.0651, 115.0542 |  |
|  |  | Mass | 132.0808 | 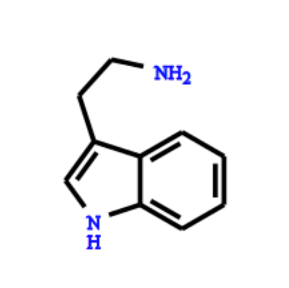 |
|  |  | Mass | 130.0651 | 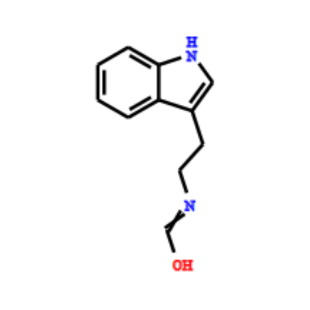 |
|  |  | Mass | 144.0808 |  |
| **36** | 2-oxochromen-7-yl (mainly trimethylated)  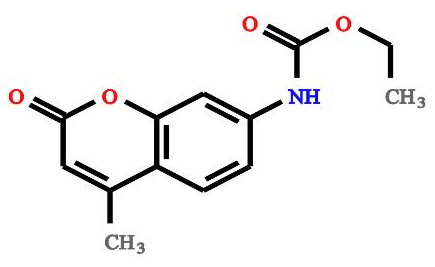 | Mass | 175.0754 | 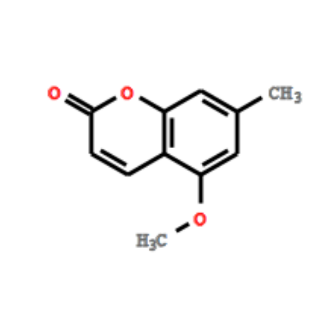 |
|  |  | Mass | 141.0699 | 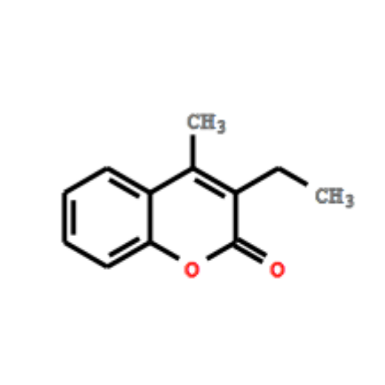 |
|  |  | Mass, Mass | 129.0699, 103.0542 | 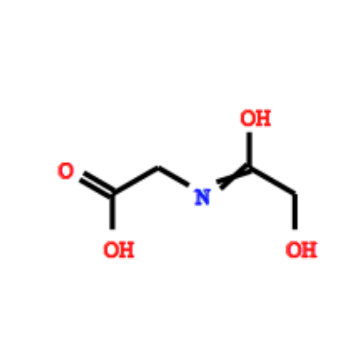 |
| **37** | Fragments indicative for cinnamic or hydroxycinnamic acid substructure  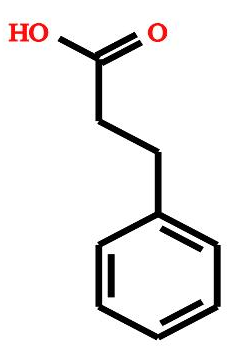 | Mass | 91.0542 | 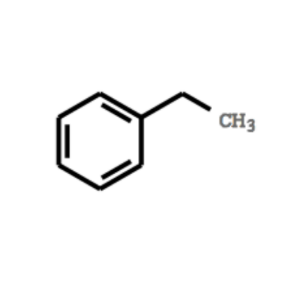 |
| **39** | Hydroxy-Pregnenolone related fragments  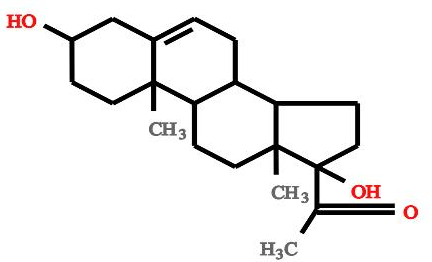 | Mass | 157.1012 | 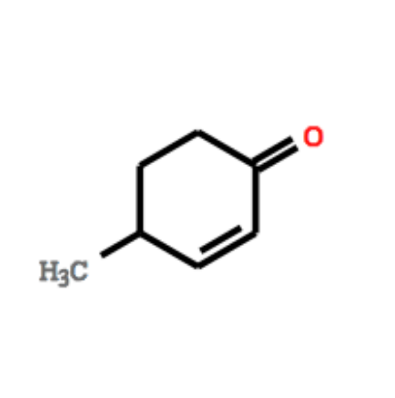 |
|  |  | Mass | 145.1012 |  |
|  |  | Mass | 119.0855 |  |
|  |  | Mass, Mass | 145.1012, 131.0855 |  |
|  |  | Mass | 143.0855 | 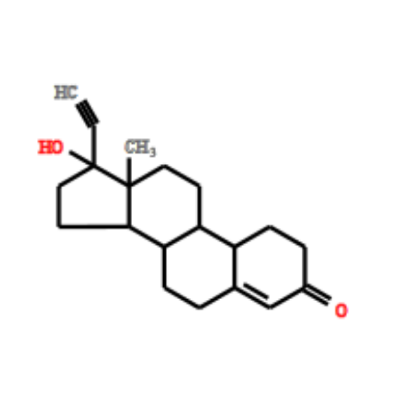 |
|  |  | Mass, Mass | 143.0855, 131.0855 |  |
|  |  | Mass, Mass | 145.1012, 119.0855 |  |
|  |  | Mass, Mass | 143.0855, 129.0699 |  |
|  |  | Mass, Mass | 119.0855, 129.0699 |  |
|  |  | Mass, Mass | 131.0855, 129.0699 |  |
|  |  | Mass | 131.0855 | 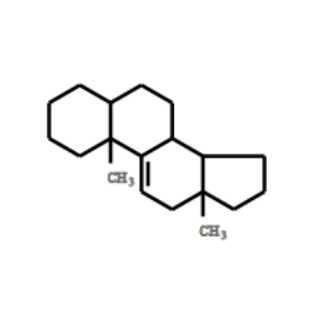 |
|  |  | Mass, Mass | 131.0855, 119.0855 |  |
| **50** | Steroid core related (C18H21 and smaller fragments)  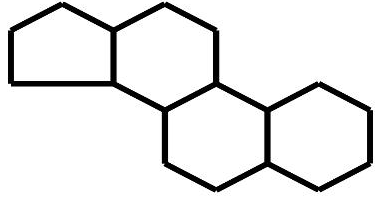 | Mass | 143.0855 | 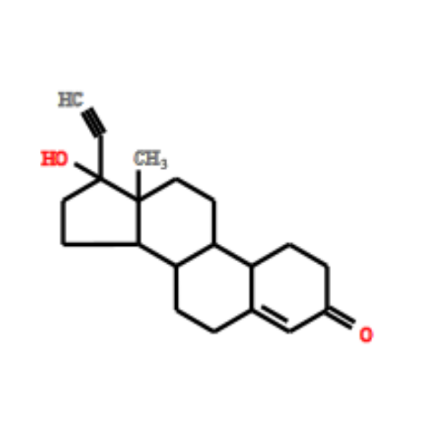 |
|  |  | Mass, Mass | 143.0855, 131.0855 |  |
|  |  | Mass, Mass | 145.1012, 119.0855 |  |
|  |  | Mass, Mass | 143.0855, 129.0699 |  |
|  |  | Mass, Mass | 155.0855 |  |
|  |  | Mass, Mass | 119.0855, 129.0699 |  |
|  |  | Mass, Mass | 131.0855, 129.0699 |  |
|  |  | Mass, Mass | 155.0855, 129.0699 |  |
|  |  | Mass, Mass | 143.0855, 128.062 |  |
|  |  | Mass | 131.0855 | 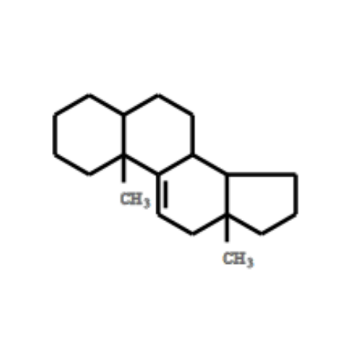 |
|  |  | Mass, Mass | 131.0855, 119.0855 |  |
|  |  | Mass, Mass | 128.062, 129.0699 | 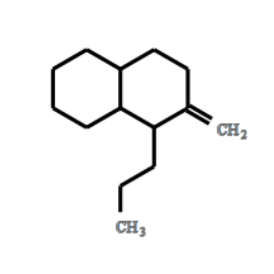 |
| **52** | Aromatic compounds related to methylbenzene  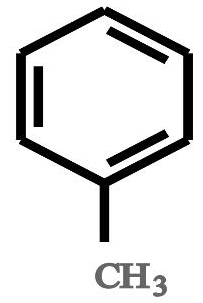 | Mass | 91.0542 | 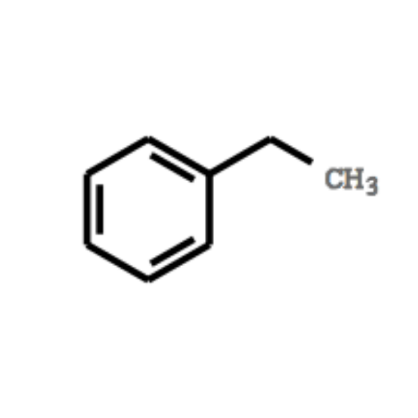 |
| **54** | Ferulic acid-based substructure  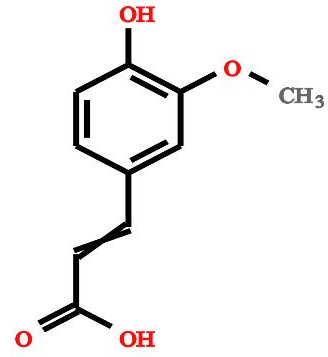 | Mass | 115.0542 | 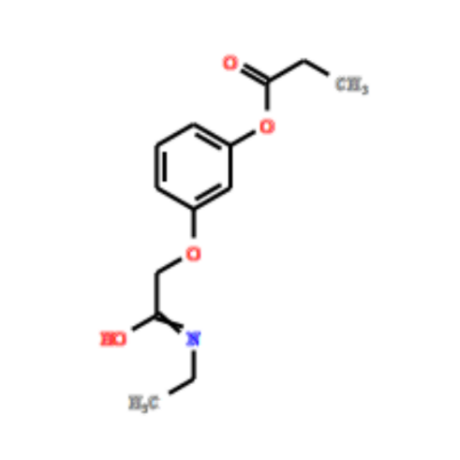 |
| **59** | Fragments indicative for [phenylalanine-CHOOH] based substructure  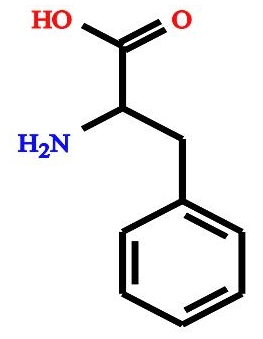 | Mass | 120.0808 | 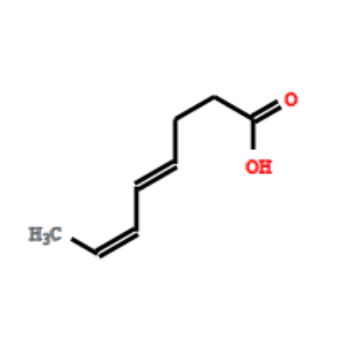 |
|  |  | Mass | 103.0542 | 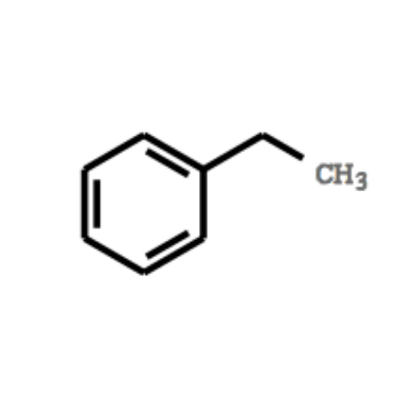 |
|  |  | Mass | 91.0542 |  |
| **65** | Oxo-1,2,3,4-tetrahydrocyclopenta[c]chromen-7-yl substructure  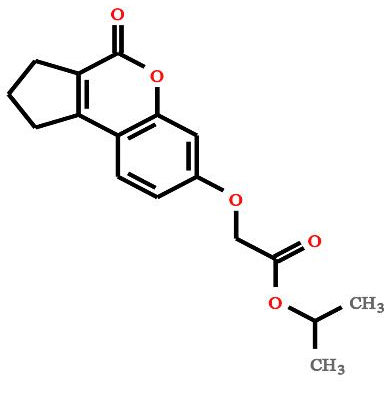 | Mass | 161.0597 | 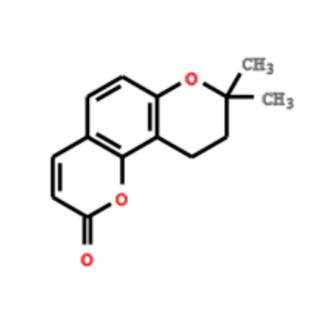 |
|  |  | Mass | 128.062 | 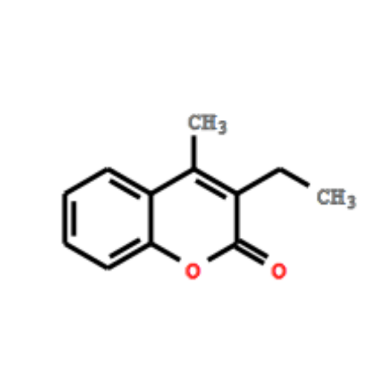 |
|  |  | Mass, Mass | 129.0699, 115.0542 |  |
|  |  | Mass, Mass | 128.062, 115.0542 |  |
|  |  | Mass, Mass | 143.0855, 115.0542 |  |
|  |  | Mass, Mass,  Mass | 128.062, 129.0699, 115.0542 |  |
| **68** | 6-oxo-6H-benzo[c]chromen-3-yl substructure  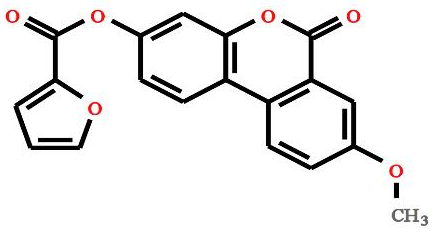 | Mass | 141.0699 | 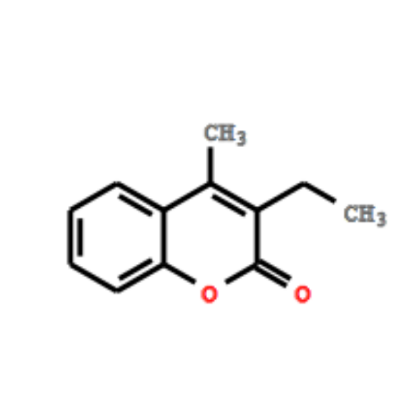 |
|  |  | Mass | 128.062 |  |
|  |  | Mass, Mass | 129.0699, 115.0542 |  |
|  |  | Mass, Mass | 128.062, 115.0542 |  |
|  |  | Mass, Mass | 141.0699, 128.062 |  |
|  |  | Mass, Mass,  Mass | 128.062, 129.0699, 115.0542 |  |
|  |  | Mass, Mass,  Mass | 141.0699, 128.062, 115.0542 |  |
| **69** | H-dibenzo (a,d) cyclohepten-5-ylidene substructure  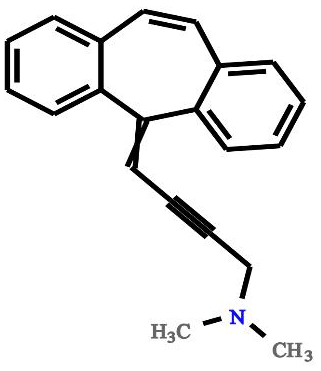 | Mass | 165.0699 | 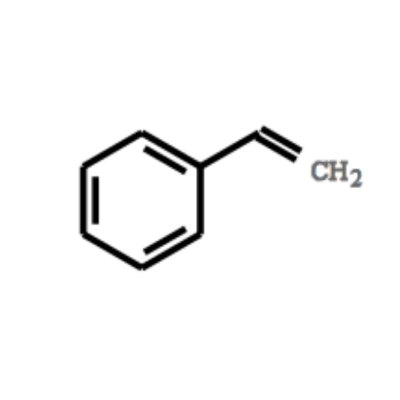 |
|  |  | Mass, Mass | 141.0699, 115.0542 | 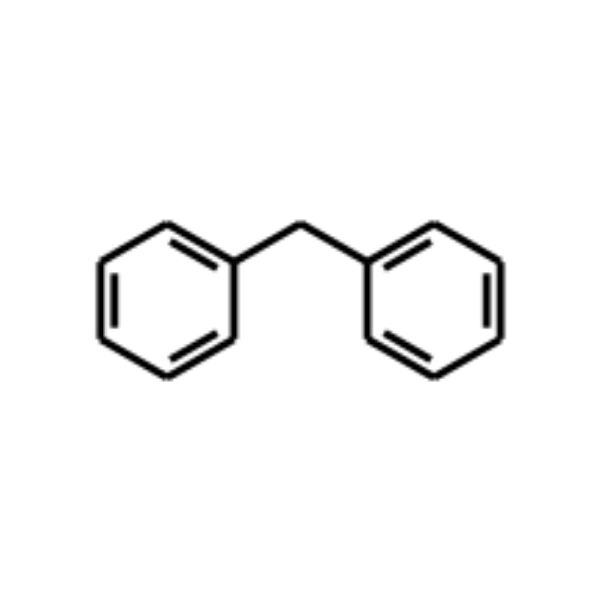 |
| **70** | Piperazine substructure (or related N-containing ring structures for C2H5N loss only)  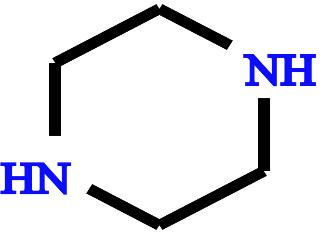 | MDiff | 43.0422 | 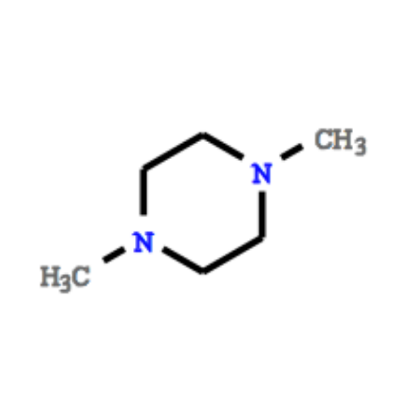 |
| **71** | 4-Methyl-6-oxo-6H-benzo[c]chromen-3-yl  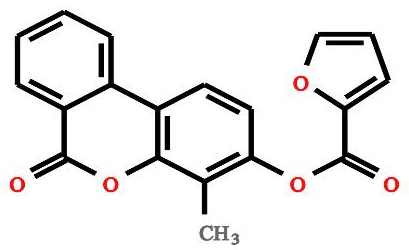 | Mass | 141.0699 | 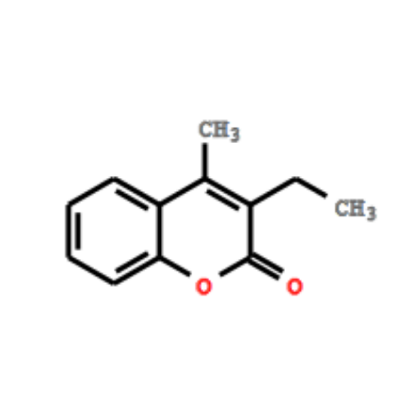 |
| **194** | Indole multi-ring substructure  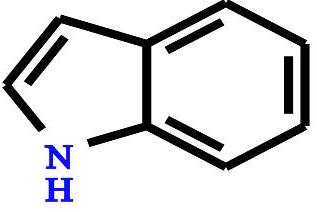 | Mass | 146.06 | 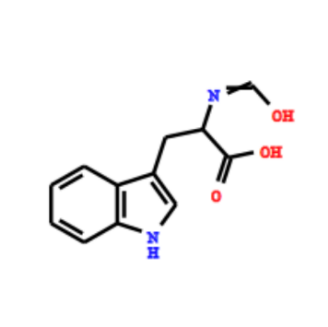 |
|  |  | Mass | 142.0651 | 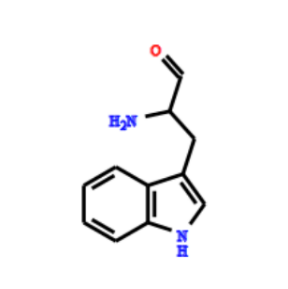 |
|  |  | Mass, Mass | 144.0808, 130.0651 |  |
|  |  | Mass | 130.0651 | 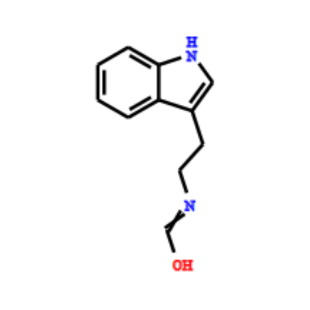 |
|  |  | Mass | 144.0808 |  |
|  |  | Mass | 154.0651 | 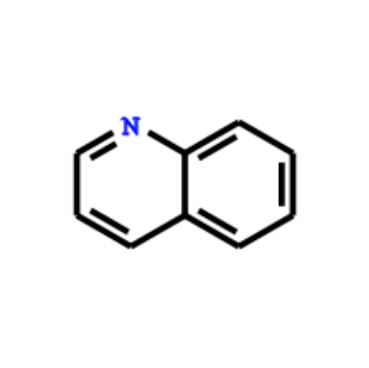 |
| **274** | 3-oxo-delta-1,4-progestogin based steroid related  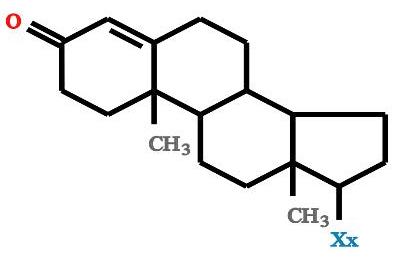 | Mass, Mass | 159.0804, 147.0804 | 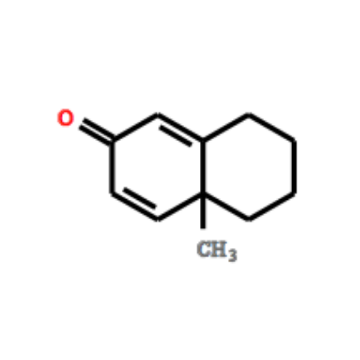 |
|  |  | Mass | 121.0648 | 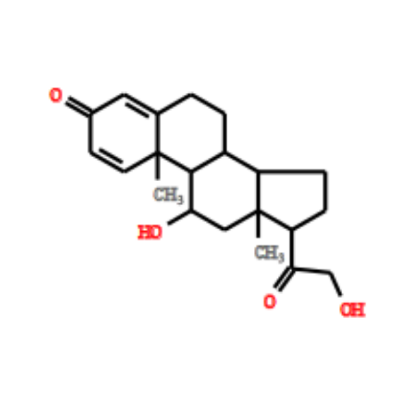 |
|  |  | Mass, Mass | 147.0804, 121.0648 |  |
|  |  | Mass, Mass | 135.0804, 121.0648 | 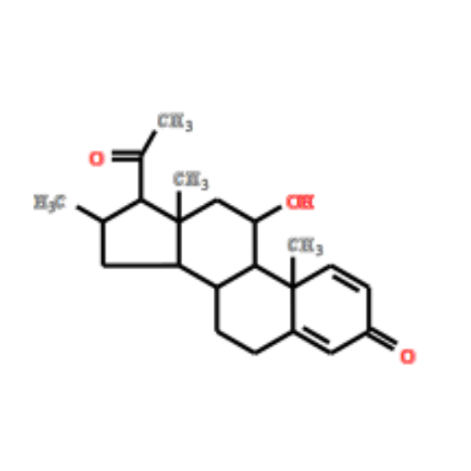 |
| **398**  **457** | 5-Hydroxyindole substructure  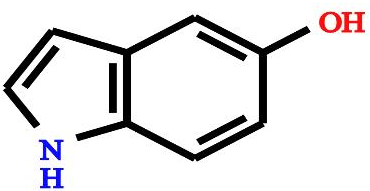 | Mass | 146.06 | 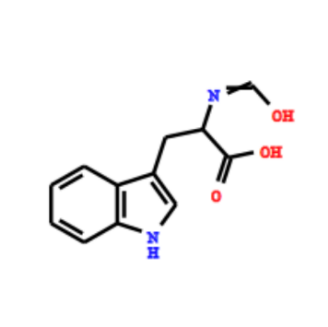 |
|  |  | Mass | 172.0757 | 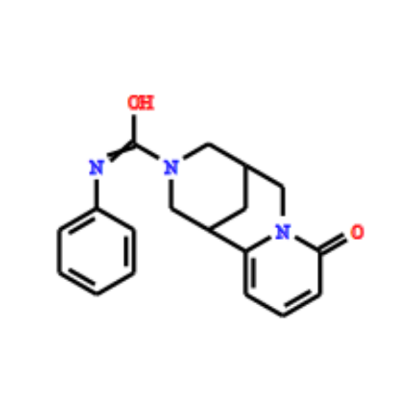 |
