## Supplementary figures and images for "MESSAR: Automated recommendation of metabolite substructures from tandem mass spectra"

### S1 Figure

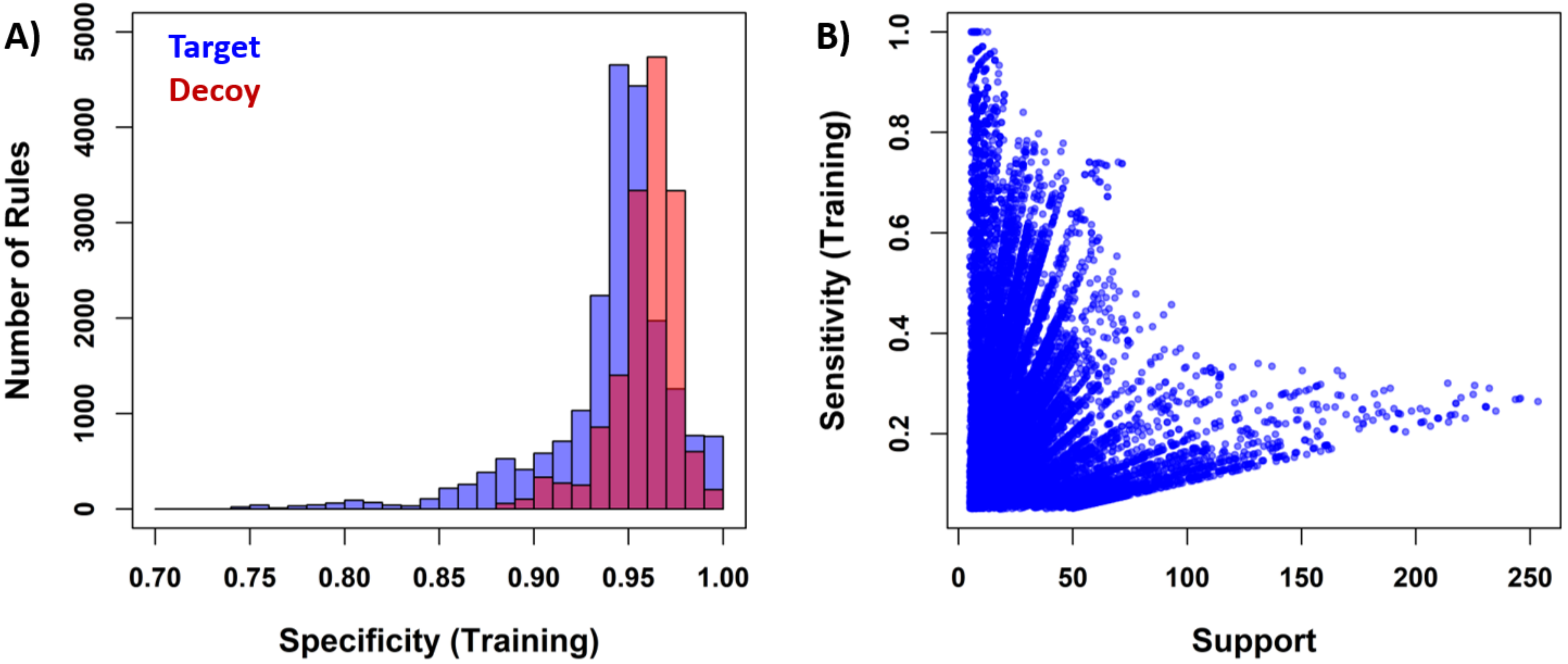

### S2 Figure

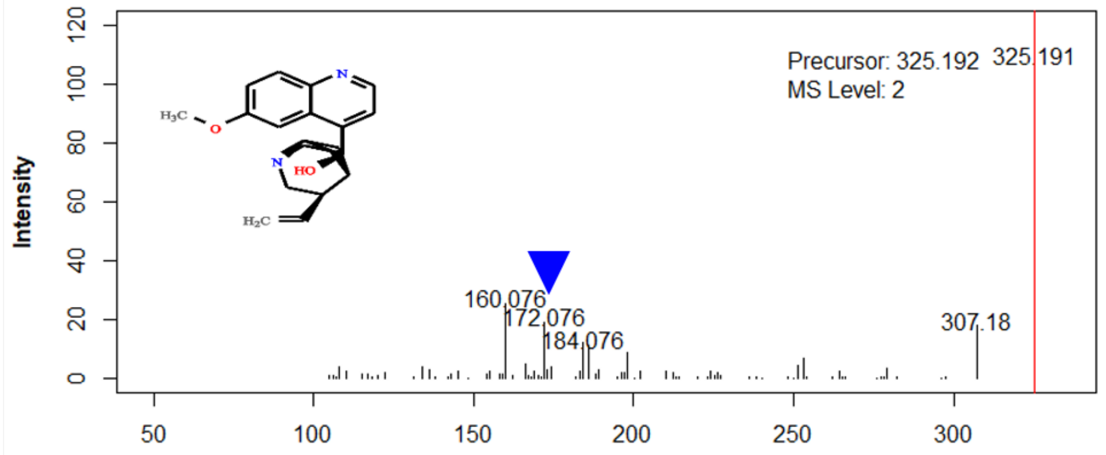

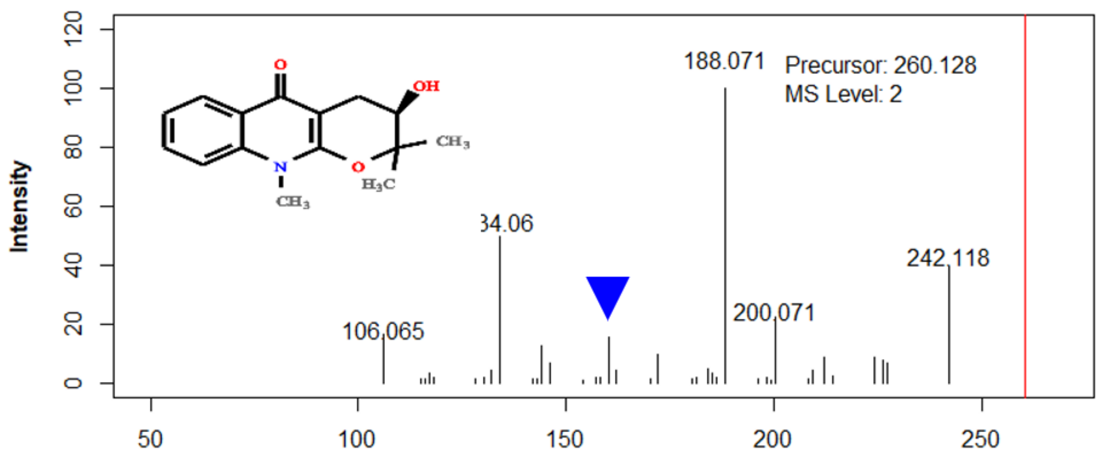

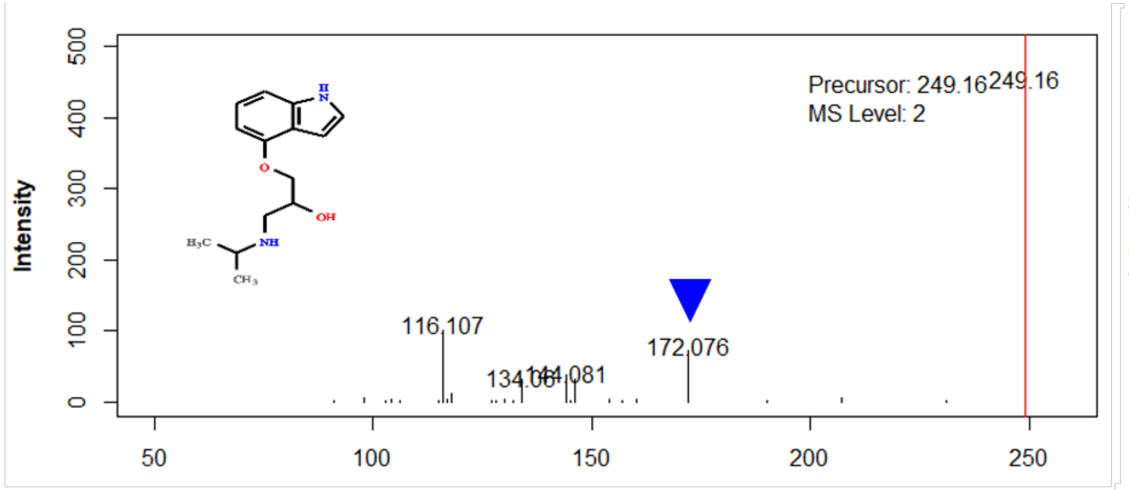

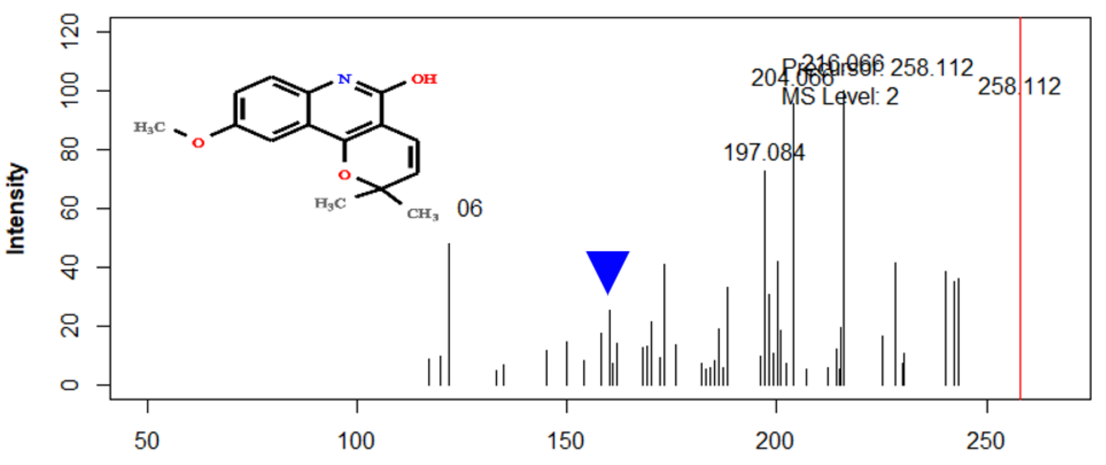

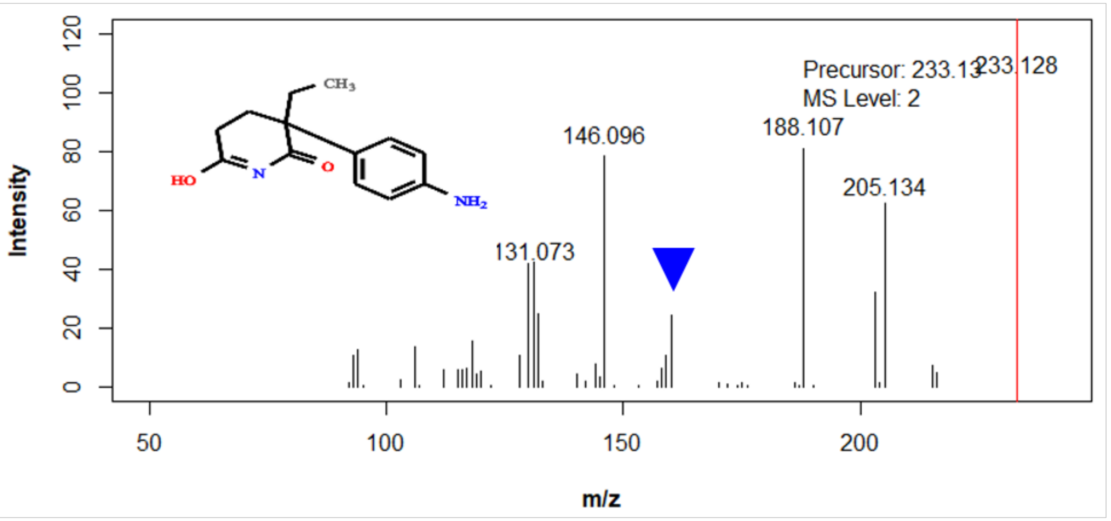

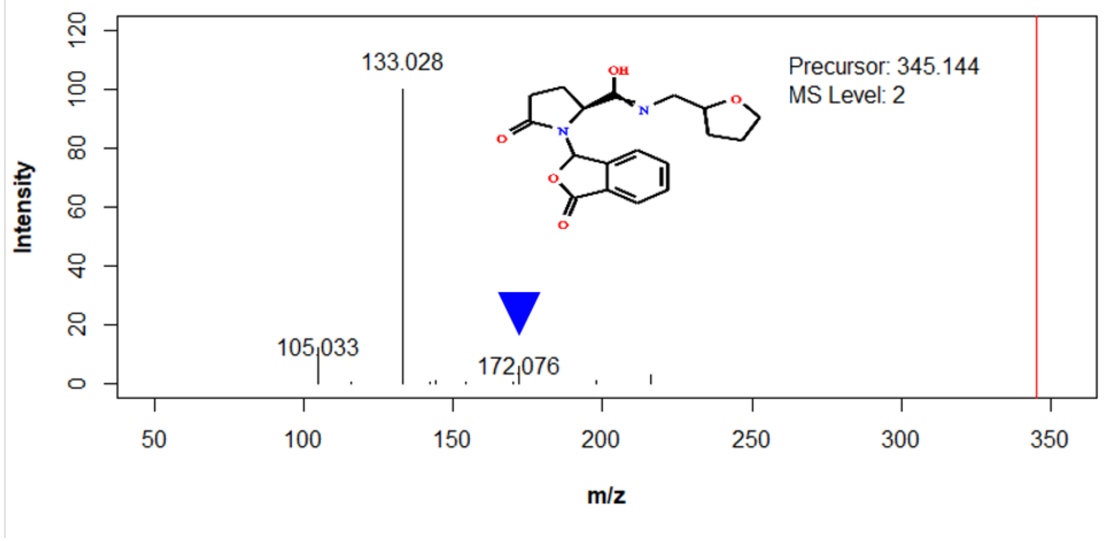

### S3 Figure

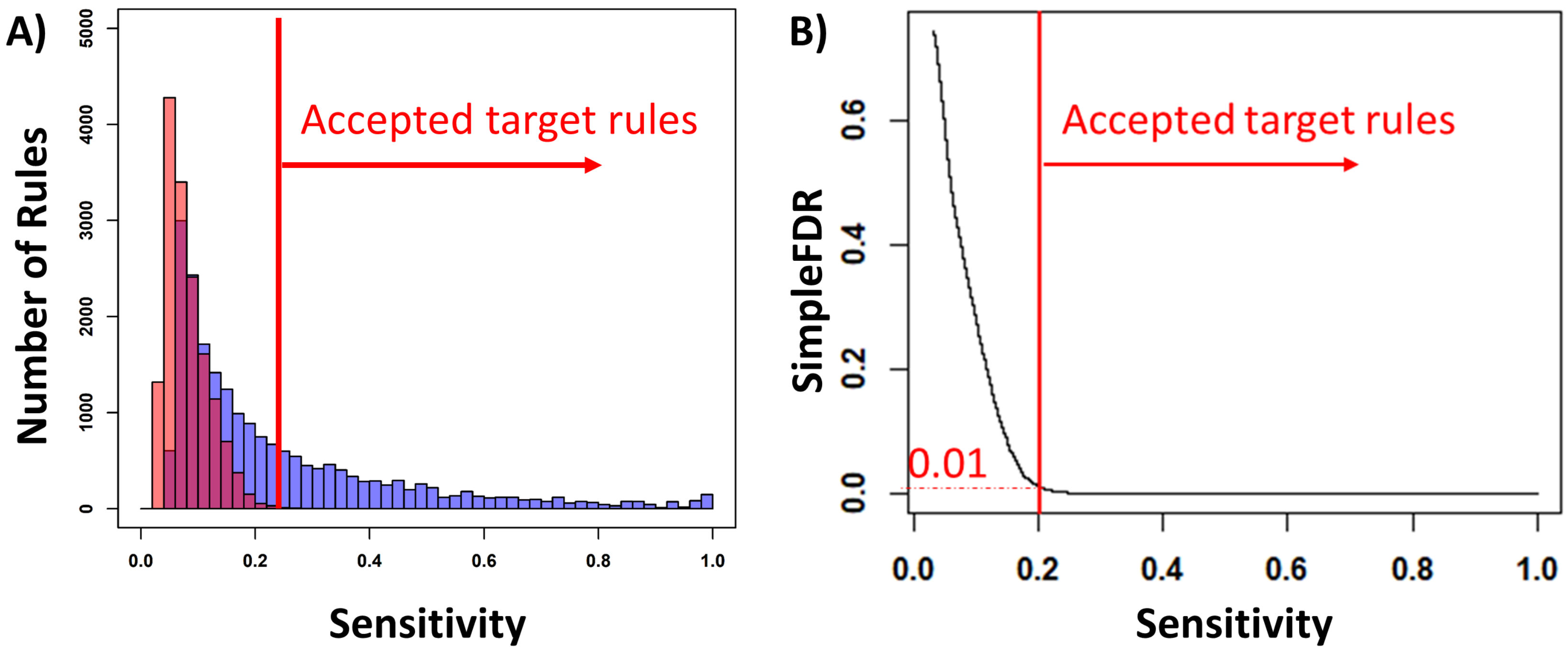
